# Heterotrimeric G proteins regulate planarian regeneration and behavior

**DOI:** 10.1101/2022.03.07.483311

**Authors:** Jennifer E. Jenkins, Rachel H. Roberts-Galbraith

## Abstract

G protein-coupled receptors (GPCRs) play broad roles in development and stem cell biology, but few roles for GPCR signaling in complex tissue regeneration have been uncovered. Planarian flatworms robustly regenerate all tissues and provide a model with which to explore potential functions for GPCR signaling in somatic regeneration and pluripotent stem cell biology. As a first step toward exploring GPCR function in planarians, we investigated downstream signal transducers that work with GPCRs, called heterotrimeric G proteins. Here, we characterized the complete heterotrimeric G protein complement in *Schmidtea mediterranea* for the first time and found that seven heterotrimeric G protein subunits promote regeneration. We further characterized two subunits critical for regeneration, *Gαq1* and *Gβ1-4a*, finding that they promote the late phase of anterior polarity re-establishment, likely through anterior pole-produced Follistatin. Incidentally, we also found that five heterotrimeric G proteins modulate planarian behavior. We further identified a putative serotonin receptor, *gcr052*, that we propose works with *Gβx2* in planarian locomotion, demonstrating the utility of our strategy for identifying relevant GPCRs. Our work provides foundational insight into roles of heterotrimeric G proteins in planarian biology and serves as a useful springboard towards broadening our understanding of GPCR signaling in adult tissue regeneration.

## Introduction

G protein-coupled receptors (GPCRs) represent one of the largest, most highly conserved, and functionally diverse families of cell surface receptors (Krishnan et al. 2012; Anantharaman et al. 2011; Langenhan et al. 2015). GPCRs also comprise ∼30% of drug targets, due to their broad involvement in cell signaling (Hopkins and Groom 2002; Wise et al. 2002; Garland 2013). GPCRs include characteristic seven-pass transmembrane domains, extracellular domains for signal perception, and intracellular domains for interaction with signal transducers (Fig. 1A) (Pierce et al. 2002; Lagerström and Schiöth 2008). Canonically, activation of the receptor initiates dissociation of a heterotrimeric G protein complex (Fig. 1A) into an α subunit and a β/γ subcomplex, both of which impact cellular function (Oldham and Hamm 2008; Smrcka 2008).

**Figure 1.**
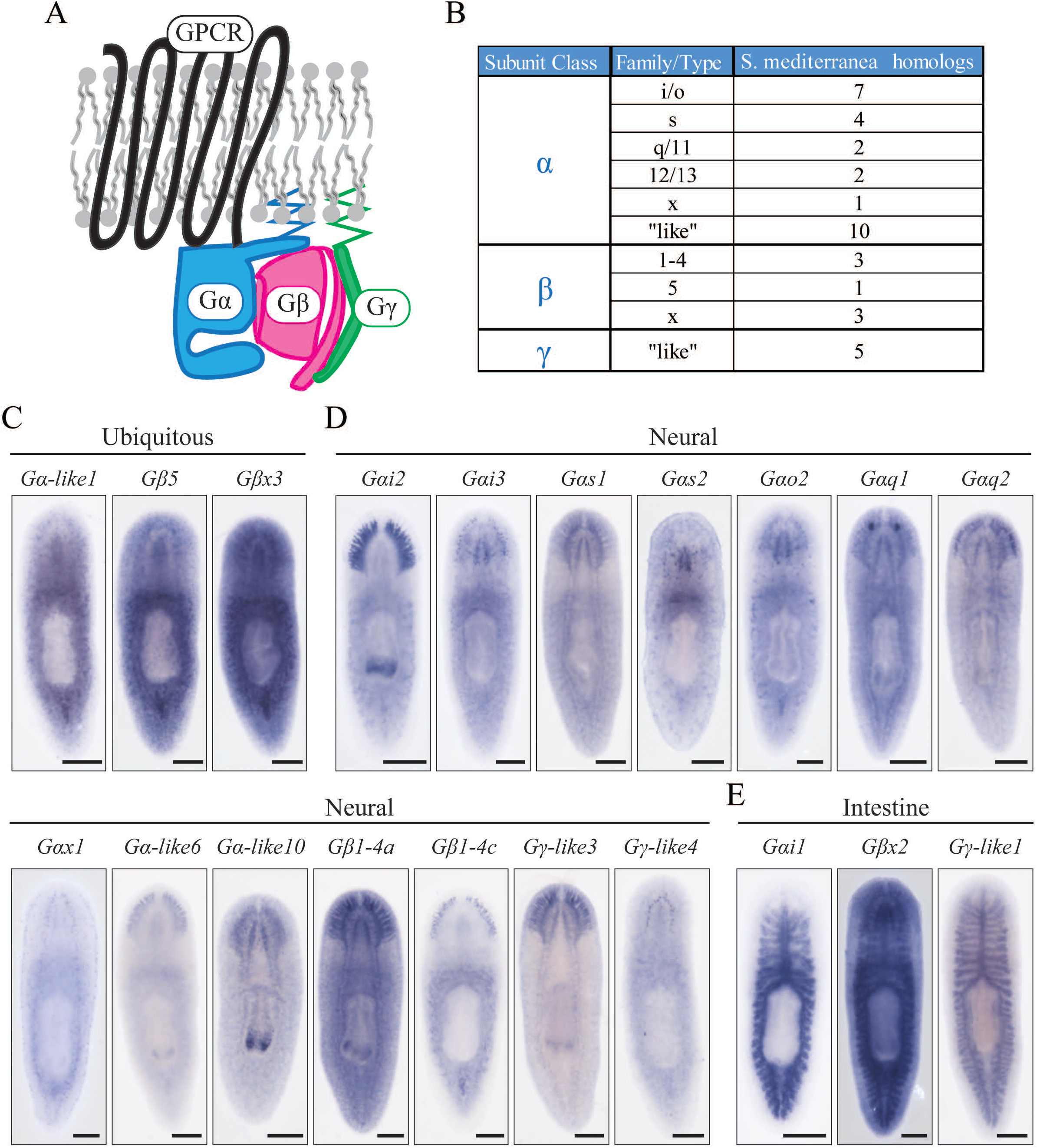
Planarians possess diverse heterotrimeric G proteins. (**A**) Graphical summary of a typical heterotrimeric G protein complex associated with a GPCR (black). The heterotrimer is composed of Gα (blue), Gβ (magenta), and Gγ (green) subunits that are activated upon ligand binding. (**B**) Table depicting *S. mediterranea* homologs for heterotrimeric G proteins. Images of genes that display ubiquitous (**C**), neural (**D**), and intestinal (**E**) expression are shown. Regions of particularly enriched or specific expression are displayed in insets. Scale = 200μm. Anterior is toward the top of the page in all figures.

Importantly, GPCR signaling regulates regeneration and wound responses throughout the animal kingdom including nematodes, fruit flies, and mammals (Ziegler et al. 2009; Doze and Perez 2013; Kiseleva et al. 2014; Zugasti et al. 2014; Choi et al. 2015; Guo et al. 2019; O’Connor et al. 2021). For example, the Protease Activated Receptor 1 (PAR1) GPCR promotes wound healing in murine skin by stimulating production of keratinocytes (Kiseleva et al. 2014). Furthermore, heterotrimeric G proteins themselves also modulate regeneration. Alpha subunits from several families have been shown to promote *or* inhibit axon regeneration in vertebrates (Bates and Meyer 1996; Li et al. 2016) and *C. elegans* (Shimizu and Hisamoto 2020). However, roles for GPCR pathways have not yet been explored in organisms that complete robust, whole-body regeneration. Studying GPCR signaling in highly regenerative models could reveal new roles for this receptor family in regeneration of complex tissues.

Freshwater flatworms called planarians provide an appealing model for investigation of mechanisms underlying robust regeneration. Planarians regenerate all tissues and complex organs *de novo*, including a brain. After nearly any injury, planarians produce a blastema in which undifferentiated cells accumulate and mature to reconstruct missing structures (Baguna et al. 1989). Planarian regeneration requires wound detection (Wenemoser et al. 2012; Wurtzel et al. 2015), activation of pluripotent adult stem cells (Wagner et al. 2011; Raz et al. 2021), and polarity re-establishment (Witchley et al. 2013; Reddien 2018). How planarian cells detect injury, reinterpret polarity axes, and mount the correct regenerative response after injury remain key areas of investigation. Because planarians require multifaceted, fine-tuned coordination of regenerative response after injury and because GPCR signaling functions in diverse aspects of cell biology and healing in other animals, we hypothesized that GPCR pathways could play key roles in planarian regeneration.

Currently the genome of *Schmidtea mediterranea* is predicted to contain 566 GPCR-encoding genes (Zamanian et al. 2011; Saberi et al. 2016). Less than 3% of these genes are functionally characterized, with the identified GPCRs promoting posterior identity, supporting planarian locomotion, coordinating germline differentiation and maintenance, facilitating reproductive system development and repair, or impacting eye regeneration (Pascual-Carreras et al. 2021; Zamanian 2011; Saberi et al. 2016; Lozano 2015). Characterization of planarian heterotrimeric G protein subunits is also limited. *gpas* is expressed in the brain branches and pharynx (Cebrià et al. 2002; Iglesias et al. 2011), while 4 other G protein subunit genes (*gna-q, gna-o, gnb*, and *gnc*) are highly expressed in the photoreceptors (Lapan and Reddien 2012). Importantly, to our knowledge, the function of heterotrimeric G proteins has not been studied in planarians.

As an essential first step toward pursuing our hypothesis that GPCR signaling promotes regeneration, we focused on heterotrimeric G proteins. In this work, we identified and characterized 38 predicted heterotrimeric G protein subunit-encoding genes in *S. mediterranea*, which includes highly conserved homologs of described vertebrate G protein families and divergent subunits. We show that 7 G protein subunit-encoding genes—*Gαs1, Gαs2, Gαq1, Gαq2, Gαo2, Gα-like6*, and *Gβ1-4a*—promote planarian regeneration. 2 of the identified genes, *Gαq1* and *Gβ1-4a*, are essential for promoting the late phase of anterior-posterior axis reestablishment, likely by influencing production of *follistatin+* anterior pole cells. We also show that 5 subunit-encoding genes—*Gαs1, Gαs2, Gαq1, Gβ1-4a*, and *Gβx2*—are required for planarian movement. To illustrate the utility of our G protein-centered approach to identifying key GPCRs, we further found a GPCR-encoding gene, *gcr052* (Saberi et al. 2016), as a potential partner of *Gβx2*. Taken together, our results reveal functions for heterotrimeric G protein signaling in the highly regenerative planarian model for the first time. Our data provide a much-needed starting point for identifying GPCRs with roles in regeneration.

## Results

### Identification of the planarian G protein subunit repertoire

To better understand G protein-coupled receptor signaling in planarians, we identified 38 G protein subunit homologs (26 Gα subunits, 7 Gβ subunits, and 5 Gγ subunits) in *S. mediterranea* transcriptomes (Brandl et al. 2016; Rozanski et al. 2019) (Fig. 1B, Supplemental Table S1). This list included all 5 previously identified planarian G protein subunit genes (Cebrià et al. 2002; Iglesias et al. 2011; Lapan and Reddien 2012). Both numbers and proportions of subunits are consistent with those found in other animals, including humans (Syrovatkina et al. 2016), *C. elegans* (Jansen et al. 1999), and *Drosophila* (Malpe et al. 2020). These results suggest that planarians utilize a typical repertoire of heterotrimeric G protein subunits.

We next classified planarian heterotrimeric subunit homologs into families using phylogenetic analysis. We classified 7 *Gαi*/o homologs, 4 *Gαs* homologs, 2 *Gαq*/11 homologs, 2 Gα12/13 homologs, 3 Gβ1-4 subgroup homologs, and 1 Gβ5 homolog (Fig. 1B; Supplemental File. S1; S2). 1 Gα homolog and 3 Gβ homologs contained all functional regions (Supplemental File. S1) but did not cluster with a specific family (Supplemental File. S2). We therefore designated these genes as “Gαx” or “Gβx” (Fig. 1B). Additionally, 10 Gα class homologs retrieved in our search were truncated, preventing accurate classification (Supplemental File. S3). We designated these genes “Gα-like” (Fig. 1B). Lastly, due to the divergent nature of Gγ homologs, we were unable to classify them into families, so we designated them as “Gγ-like” (Fig. 1B; Supplemental File. S1; S2). Our phylogenetic analysis indicates that the Gα class homolog *gpas* (Cebrià et al. 2002; Iglesias et al. 2011) was previously misclassified, and the name *Gαi2* more accurately represents this subunit’s classification.

After defining the *S. mediterranea* heterotrimeric G protein complement, we next sought to characterize the expression patterns of these genes, potentially providing insight into tissue-specific roles and possible heterotrimer combinations. We observed ubiquitous expression for 10 G protein subunit homologs (Fig. 1C; Supplemental Fig. S1). However, many subunits showed tissue-specific expression in the nervous system (Fig. 1D; Supplemental Fig. S1) or the intestine (Fig. 1E). In addition to our observations, we determined that 32 of the 38 subunits are expressed within stem cells based on available transcriptomic resources (Labbé et al. 2012; Fincher et al. 2018; Plass et al. 2018; Zeng et al. 2018) (Supplemental Table S1). Our results suggest that *S. mediterranea* heterotrimeric G proteins likely function in many different tissue types throughout the animal, including stem cells and a diverse set of neural cell types.

### Planarian heterotrimeric G proteins function in regeneration

We then performed the first investigation into roles for planarian heterotrimeric G proteins in regeneration. After screening 36 of the 38 predicted subunit genes, we found 7 genes for which RNAi caused significant reduction in brain regeneration (Fig. 2C-E). Of these candidates, RNAi targeting *Gαs1, Gαs2, Gαo2, Gαq2*, or *Gα-like6* produced a modest effect (Fig. 2C-E). RNAi targeting *Gαq1* or *Gβ1-4a* caused a severe reduction of brain regeneration by ∼67% and ∼86%, respectively, compared to controls (Fig. 2C-E). Interestingly, we detected no regeneration phenotypes after RNAi targeting Gγ subunit genes, potentially indicating functional redundancy among Gγ subunits.

**Figure 2.**
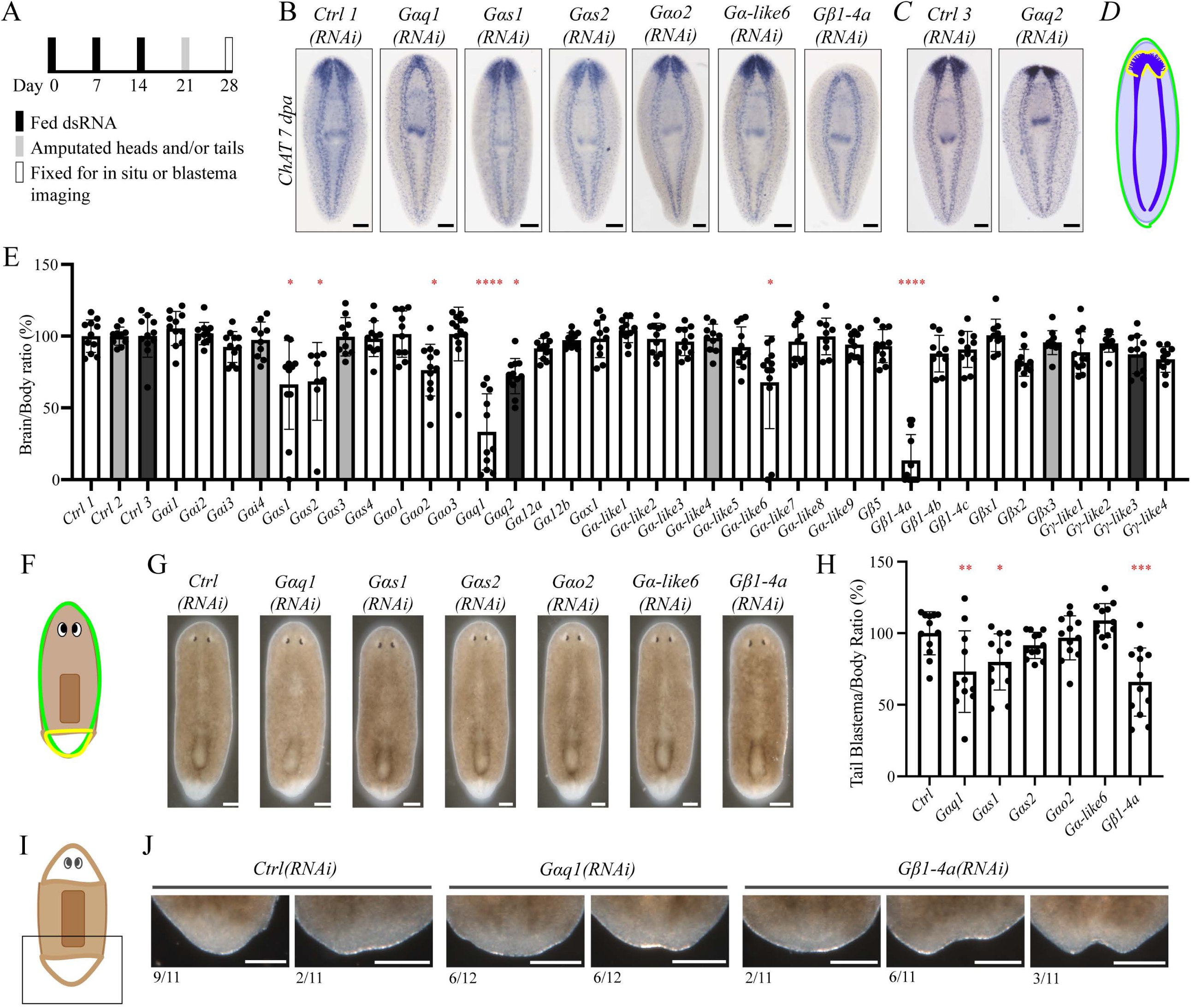
Specific planarian heterotrimeric G protein genes promote brain regeneration. (**A**) RNAi paradigm used for regeneration screens. (**B and C**) Representative images showing animals treated with RNAi targeting genes that reduced brain regeneration, visualized using ISH with *ChAT*. (**D**) Visual schematic displaying brain regeneration quantification. The area of the brain (yellow) and body (green) for each animal are used to calculate brain/body ratios (Roberts-Galbraith et al. 2016). (**E**) Bar graph of data from quantification of brain/body ratios after RNAi, displaying mean and standard deviation. Bars are color coded to match samples and controls from the same experiment. * = P value ≤ 0.05. ** = P value ≤ 0.005. *** = P value ≤ 0.0001. (**F**) Visual schematic displaying quantification of tail blastema formation. The quantification method referred to in (D) was applied to measure tail blastemas. The area of the tail blastema (yellow) and body (green) for each animal were used to calculate blastema/body ratios. (**G**) Representative images showing animals treated with RNAi 7 dpa. (**H**) Bar graph with quantification of tail blastema/body ratios. Data were graphed and analyzed as in Fig. 2E. (**I**) Visual schematic showing image perspective in (J). (**J**) Representative images showing asymmetrical and notched tail blastemas. Scale = 200μm.

Defective brain regeneration could be brain-specific or could reflect reduced regeneration independent of tissue identity. To distinguish between these possibilities, we examined posterior regeneration in knockdown animals. We saw significantly reduced tail blastema size after RNAi targeting *Gαs1, Gαq1*, or *Gβ1-4a* (Fig. 2F-H). Noticeably, tail regeneration was less affected than head regeneration for these genes. We also detected asymmetrical and/or notched tail blastemas in half of *Gαq1(RNAi)* animals and the majority of *Gβ1-4a(RNAi)* animals (Fig. 2I-J). Our results suggest that *Gαs2, Gαo2, Gαq2*, and *Gα-like6* specifically support brain regeneration. In contrast, *Gαs1, Gαq1*, and *Gβ1-4a* likely play roles in whole-body regeneration. Additionally, our results demonstrate an important role for *Gβ1-4a* in tail blastema morphology. In summary, we show that multiple heterotrimeric G proteins promote regeneration, with *Gαq1* and *Gβ1-4a* playing especially critical roles.

### Elucidation of roles for heterotrimeric G proteins in planarian behavior

During our RNAi screen (Fig. 2A), we incidentally observed that knockdown of 5 heterotrimeric G protein genes caused behavioral phenotypes. The strongest behavior we documented was reduced movement and paralysis in *Gαs1(RNAi)* animals. These animals reacted more slowly when on their dorsal side and sometimes displayed no reaction at all (Supplemental Movie S1). We quantified this reduced capacity for *Gas1(RNAi)* animals to right themselves. All control animals righted themselves, taking an average of 27.35 seconds (Fig. 3B-C). In contrast, 5 of 20 *Gαs1(RNAi)* animals failed to flip onto their ventral side within 5 minutes. The remaining *Gas1(RNAi)* animals took an average of 168 seconds to flip (Fig. 3B-C). These results indicate that *Gas1* is required for the righting response in planarians.

**Figure 3.**
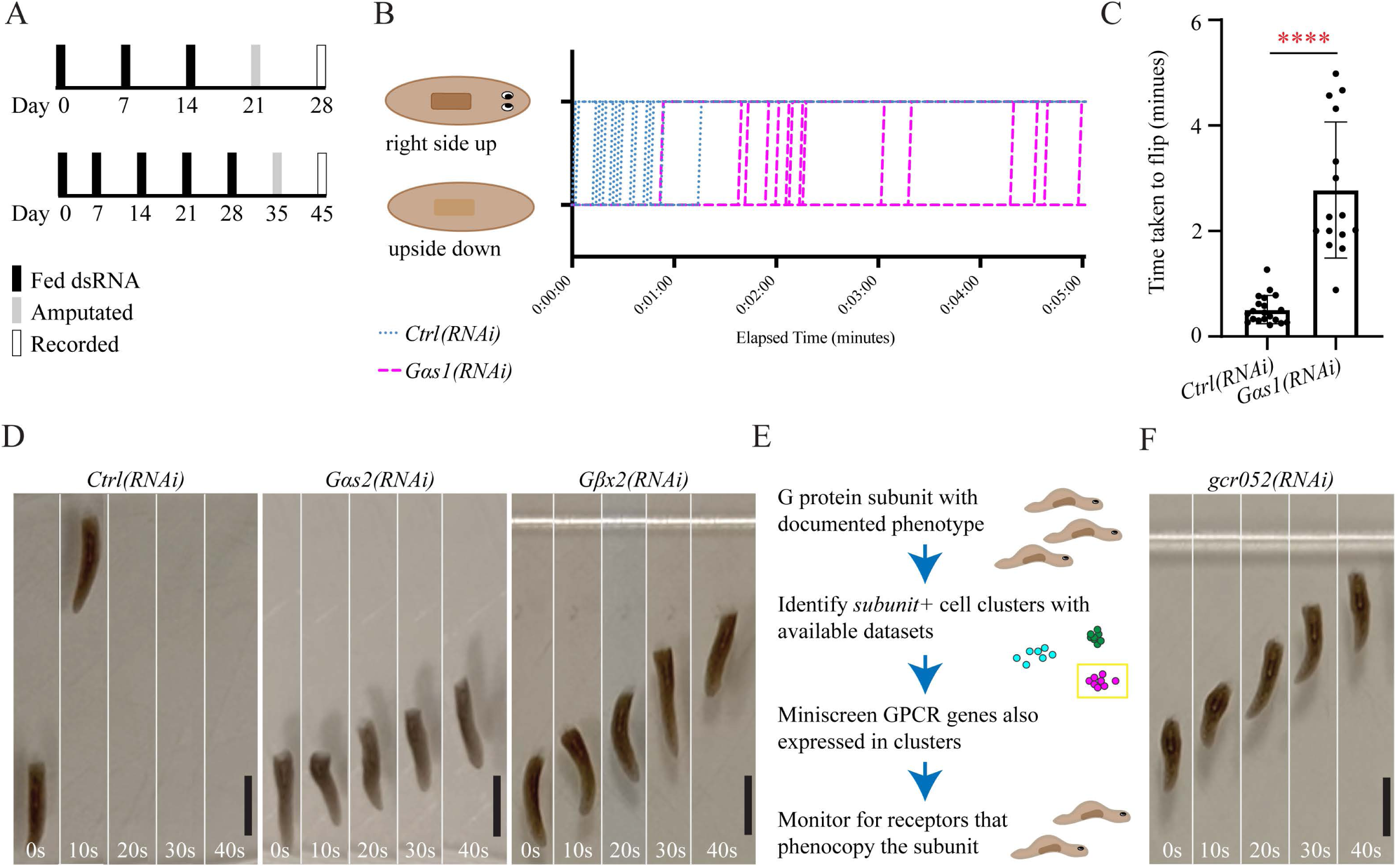
Planarian heterotrimeric G proteins and GCR052 promote animal movement. (**A**) RNAi paradigms used during initial screens (top) and for long-term treatment (bottom). Data from (B-C) resulted from initial screen paradigm, and data from (D-F) resulted from long-term paradigm. (**B**) Flip assay used to document paralysis. The graph includes flipping data for 20 animals per RNAi condition. (**C**) Bar graph showing mean times taken for animals to flip (excluding 5 non-flipping *Gas1(RNAi)* animals). **** = P value ≤ 0.0001. (**D**) Image stills from videos capturing locomotion displayed by control, *Gas2(RNAi)*, and *Gβx2(RNAi)* animals 7 dpa. (**E**) Graphical scheme showing the method used to identify candidate GPCRs for G protein subunits with documented phenotypes. (**F**) Image stills from videos capturing locomotion displayed by *gcr052(RNAi)* animals 7 dpa. Data from (D) and (F) were taken from a single experiment. Scale = 2 mm.

Inhibition of any of 4 genes—*Gαs2, Gβx2, Gβ1-4a, and Gαq1*—resulted in decreased gliding movement, which caused “inching” behavior (Glazer et al. 2010). The quickest effects were seen following RNAi of *Gαs2* or *Gβx2* (Fig. 3D; Supplemental Movies S2-4). Inching was detectable by the third RNAi feeding, resulting in reduced distance traveled over time (Supplemental Fig. S2). After long-term RNAi, *Gβ1-4a(RNAi)* animals alternated between inching and gliding (Supplemental Fig. S3; Supplemental Movies S5-6), and *Gαq1(RNAi)* animals displayed labored inching undetectable in real time (Supplemental Fig. S3; Supplemental Movies S5; S7).

*Gαs2(RNAi)* and *Gβx2(RNAi)* animals were indistinguishable, which led us to hypothesize that these subunits may be operating in the same cells. Available single cell sequencing data (Fincher et al. 2018) indicated coexpression of *Gαs2* and *Gβx2* in a specific group of neurons (neural subcluster 31), which supported this hypothesis (Supplemental Table S3). We then sought to identify the GPCR that works with *Gαs2* and *Gβx2*. We identified and screened 7 GPCR genes coexpressed with *Gαs2* or *Gβx2* (Fincher et al. 2018) (Fig. 3E; Supplemental Table S3). Using this method, we identified a putative serotonin receptor, *gcr052*, for which knockdown caused inching indistinguishable from that displayed by *Gαs2(RNAi)* and *Gβx2(RNAi)* animals (Fig. 3F and Supplemental Movies S2-4; S8). *gcr052* is expressed in neural clusters 8 and 18 with *Gβx2* (Fincher et al. 2018) (Supplemental Table S3). We thus propose that *Gβx*2 might act downstream of the GCR052 receptor to support gliding motion.

Together, our results show that 5 planarian heterotrimeric G proteins are essential for normal animal movement. Additionally, our identification of GCR052 provides proof-of-principle that the heterotrimeric G proteins characterized in this work can accelerate planarian GPCR research. Further, although we saw overlap between regeneration and behavior after perturbation of some genes (*Gαs1, Gαs2, Gαq1* and *Gβ1-4a*), other genes specifically promote regeneration (*Gαo2, Gαq2, and Gα-like6*) or behavior (*Gβx2)* (Fig. 2, 3, and Supplemental Table S2).

### Exploring roles for Gαq1 and Gβ1-4a

After our initial characterization of the role of planarian G proteins in regeneration and behavior, we focused on the 2 subunits with most prominent roles in planarian regeneration, *Gαq1* and *Gβ1-4a*. To determine whether the regeneration defects observed after *Gαq1(RNAi)* or *Gβ1-4a(RNAi)* resulted from perturbed stem cell maintenance or differentiation, we looked at expression of a stem cell marker (*Smedwi-1* [Reddien et al. 2005]) and epidermal progenitor markers (*AGAT-1* and *prog-1* [Eisenhoffer et al. 2008; Tu et al. 2015]) 7 days post amputation (dpa). We did not see loss of stem cells or progenitors through in situ hybridization (ISH) (Fig. 4A-C). However, expression of *Smedwi-1* after *Gαq1(RNAi)* showed a modest reduction through Quantitative Reverse Transcription Polymerase Chain Reaction (RT-qPCR) (Fig. 4D). Altogether, we conclude that *Gαq1* and *Gβ1-4a* are not broadly required for stem cell maintenance or differentiation, but they could play subtle roles in stem cell function.

**Figure 4.**
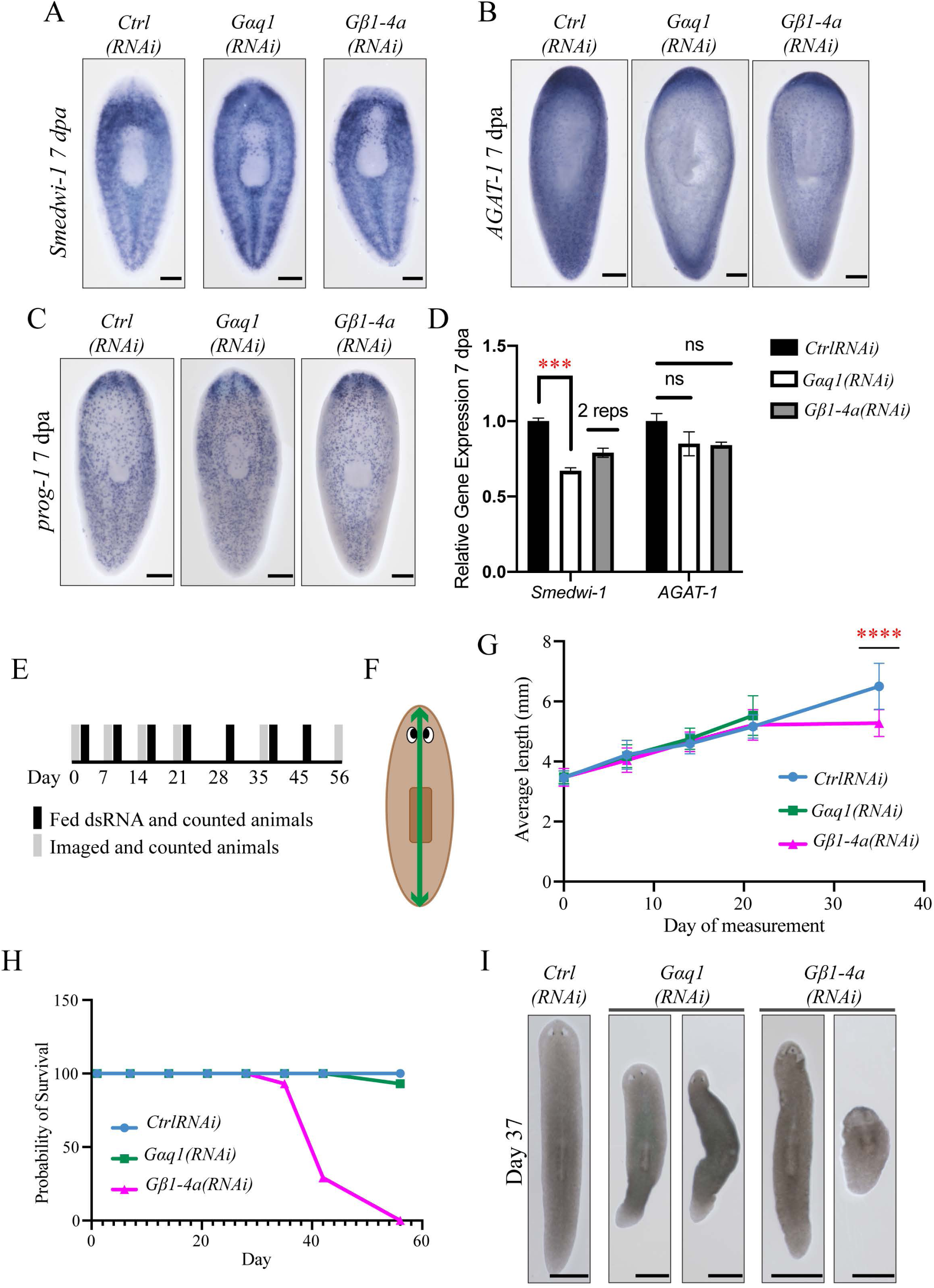
*Gαq1* promotes stem cell marker expression and *Gβ1-4a* supports growth and survival. Representative images of *Smedwi-1* (**A**), *AGAT-1* (**B**), and *prog-1* (**C**) ISH 7 dpa in animals after RNAi targeting *Gαq1*, and *Gβ1-4a*. (**D**) Relative expression levels of stem cell markers measured by RT-qPCR. Error bars represent SEM. *** = P value ≤ 0.0005. (**E**) Homeostatic roles for *Gαq1* and *Gβ1-4a* were determined through a long-term RNAi paradigm. (**F**) Graphical depiction of animal length measurement. (**G**) Growth curve data showing mean animal length over time. (**H**) Survival curve showing the relative percentage of surviving animals after each week of RNAi. (**I**) Representative images of homeostatic phenotypes on day 37. Scale: 200μm (A-C), 1 mm (I).

We next considered whether these genes affected injury response. We first examined their expression after injury. *Gαq1* and *Gβ1-4a* are upregulated at the amputation site at both 6 hours post-amputation (hpa) and 3 dpa (Supplemental Fig. S4A). We also determined that *Gαq1(RNAi)* and *Gβ1-4a(RNAi)* animals had largely unaffected expression of wound-induced genes at 6 hpa (Supplemental Fig. S4B-C). These results suggest that both genes are upregulated during regeneration, but that targeting *Gαq1* and *Gβ1-4a* does not appear to broadly affect injury response.

Finally, we asked whether the roles of *Gαq1* and *Gβ1-4a* were exclusive to regeneration or whether either gene functioned during homeostasis. We adopted a long-term RNAi paradigm and measured animal growth and survival over time (Fig. 4F). *Gβ1-4a(RNAi)* animals ceased growth after day 21 and we halted the growth measurements of *Gαq1(RNAi)* animals because they fissioned (Fig. 4G). Long-term RNAi targeting *Gβ1-4a* was lethal, with animals showing head lysis and dying near day 40 (Fig. 4H-I). We also noted postural changes and no pattern of lethality in *Gαq1(RNAi)* animals (Fig. 4I).

To summarize, our data indicate that *Gαq1* and *Gβ1-4a* are essential for regeneration but not strictly required for stem cell survival and wound response induction. Further, long-term inhibition of *Gβ1-4a*, but not *Gαq1*, is lethal, indicating a key difference in function for these G protein subunits.

### Gαq1 and Gβ1-4a support the late phase of anterior-posterior polarity re-establishment

Early in planarian regeneration, tissues reorganize to pattern the body axes using conserved developmental signaling pathways (e.g., Wnt [Petersen and Reddien 2008; Iglesias et al. 2008; Gurley et al. 2008]). After ruling out global stem cell loss and wound response failure, we next considered whether polarity is perturbed in *Gαq1(RNAi)* and *Gβ1-4a(RNAi)* animals after amputation.

During the early phase of polarity reestablishment, the remaining tissue determines which end of the animal is anterior and which is posterior (Fig. 5A) (reviewed in Owlarn and Bartscherer 2016). To determine whether *Gαq1(RNAi)* and *Gβ1-4a(RNAi)* animals correctly complete the initial anterior-posterior decision, we looked at *notum* and *wnt1* expression 18 hpa. *notum* expression was localized correctly in *Gαq1(RNAi)* and *Gβ1-4a(RNAi)* animals (Fig. 5C). *Gαq1(RNAi)* animals also expressed *wnt1* normally, but most *Gβ1-4a(RNAi)* animals displayed posterior-enriched expression of *wnt1* (Fig. 5C). These results suggest that *Gαq1* is not involved in early polarity decisions, but *Gβ1-4a* might affect anterior wound-induced *wnt1* expression.

**Figure 5.**
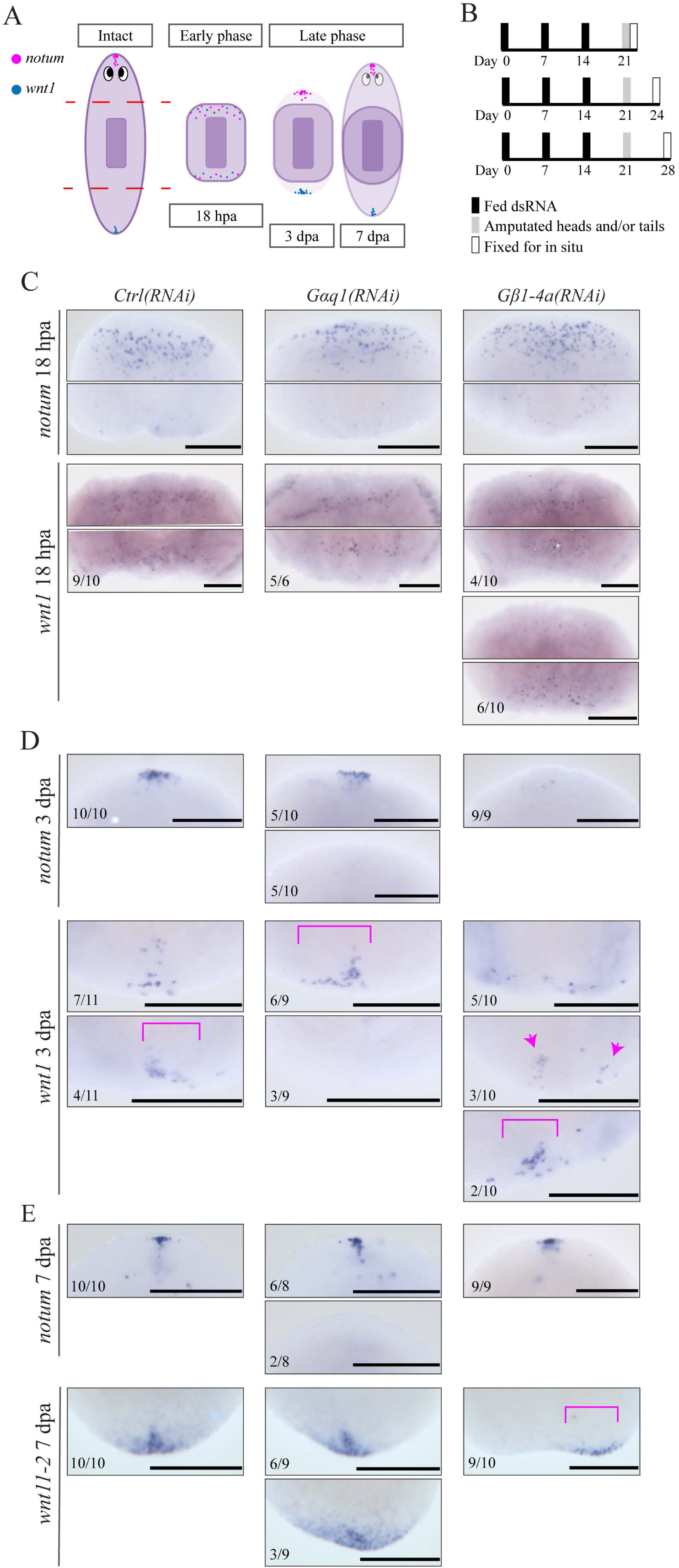
*Gαq1* and *Gβ1-4a* support the late phase of anterior and posterior pole regeneration. (**A**) Graphic summary depicting phases of polarity reestablishment after head and tail amputation, as summarized by (Owlarn and Bartscherer 2016). (**B**) RNAi paradigms for 18 hpa (top), 3 dpa (middle), and 7 dpa (bottom). (**C-E**) The following images are zoomed to focus on the regenerating head or tail blastemas for each stage. Representative images of anterior *notum*, and posterior *wnt1* or *wnt11-2* expression at 18 hpa, 3 dpa, and 7 dpa of heads and/or tails. Magenta brackets denote non-medial expression domains. Magenta arrowheads indicate multiple expression domains. Scale = 200 μm.

After re-initiation of axial polarity, anterior and posterior “poles” form at the distal ends of the planarian body (Fig. 5A). To determine whether pole formation was disrupted in *Gαq1(RNAi)* and *Gβ1-4a(RNAi)* animals, we analyzed *notum* and *wnt1* expression at 3 dpa. 50% of *Gαq1(RNAi)* animals and all *Gβ1-4a(RNAi)* animals lacked anterior *notum* expression (Fig. 5D). Additionally, *Gαq1(RNAi)* animals displayed an asymmetric *wnt1* pattern or absent *wnt1* in the posterior domain, and *Gβ1-4a(RNAi)* animals regenerated with a broadened and/or asymmetrical domain of *wnt1* expression (Fig. 5D). Our results indicate that *Gαq1* and *Gβ1-4a* impact the process of pole formation at both anterior and posterior ends of the animal.

Anterior and posterior poles further coalesce and mature during the late phase of polarity reestablishment (Fig. 5A). To investigate whether *Gαq1* and *Gβ1-4a* support the maturation of the key polarity domains, we examined *notum* or another posterior marker, *wnt11-2* (Gurley et al. 2008) expression in knockdown animals at 7 dpa. We observed a lack of *notum* staining at the anterior pole in ∼25% of *Gαq1(RNAi)* animals (Fig. 5E), and we confirmed this pattern by using a second mature pole marker, *sFRP-1* (Petersen and Reddien 2008; Gurley et al. 2008) (Supplemental Fig. S5A). Posterior pole maturation was also disrupted in ∼33% of *Gαq1(RNAi)* animals, which displayed broader and more diffuse *wnt11-2* expression (Fig. 5E). Surprisingly, all *Gβ1-4a(RNAi)* animals restored anterior *notum* expression by 7 dpa, although the domains appeared less consolidated than in control animals, which could signify slower maturation (Fig. 5E, Supplemental Fig. S5A). Strikingly, most *Gβ1-4a*(RNAi) animals expressed posterior *wnt11-2* asymmetrically, with staining on either side of the animal’s midline (Fig. 5E). Though notched tails were commonly seen after *Gβ1-4a(RNAi)* (Fig. 2J), our data did not support the presence of a secondary anterior domain, as has been seen after other RNAi treatments (Supplemental Fig. S5B-C) (Cloutier et al. 2021).

Taken together, *Gβ1-4a* supports the speed of anterior pole reestablishment and promotes proper midline placement of the posterior pole. Our data also support a role for *Gαq1* in promoting *notum*^+^ anterior pole cell regeneration. Interestingly, while both *Gαq1* and *Gβ1-4a* function during regeneration and influence anteroposterior polarity, the precise phenotypes seen after RNAi of *Gαq1* and *Gβ1-4a* are distinct.

### Gαq1 promotes head regeneration through production and activity of follistatin+ anterior pole cells

The anterior pole is established and maintained through two mutually dependent signaling proteins, Notum and Follistatin (Roberts-Galbraith and Newmark 2013; Gaviño et al. 2013; Petersen and Reddien 2011). *notum* and *follistatin* encode key extracellular inhibitors of posterior-promoting Activin and Wnt pathways, respectively (Nakamura et al. 1990; Kakugawa et al. 2015). We noted several similarities between the phenotypes caused by *follistatin(RNAi)* and those caused by *Gαq1(RNAi)* or *Gβ1-4a(RNAi)*. Similarities include: strong impacts on head and brain regeneration; reduced or delayed *notum* expression in the regenerating head; unaffected expression of early wound response genes; and subtle impacts on stem cells.

Based on phenotypic similarities, we sought to determine whether RNAi of *Gαq1* or *Gβ1-4a* impacts *follistatin* expression during regeneration (Fig. 6A-E). We detected no change in *follistatin* expression at 12 hpa after perturbation of *Gαq1* or *Gβ1-4a* (Fig. 6C). We similarly saw equivalent or higher *follistatin* transcripts 6 hpa through RT-qPCR (Supplemental Fig. S4C). These results indicate that regeneration failure in *Gαq1(RNAi)* and *Gβ1-4a(RNAi)* animals is not correlated with a reduction of wound-induced *follistatin* expression.

**Figure 6.**
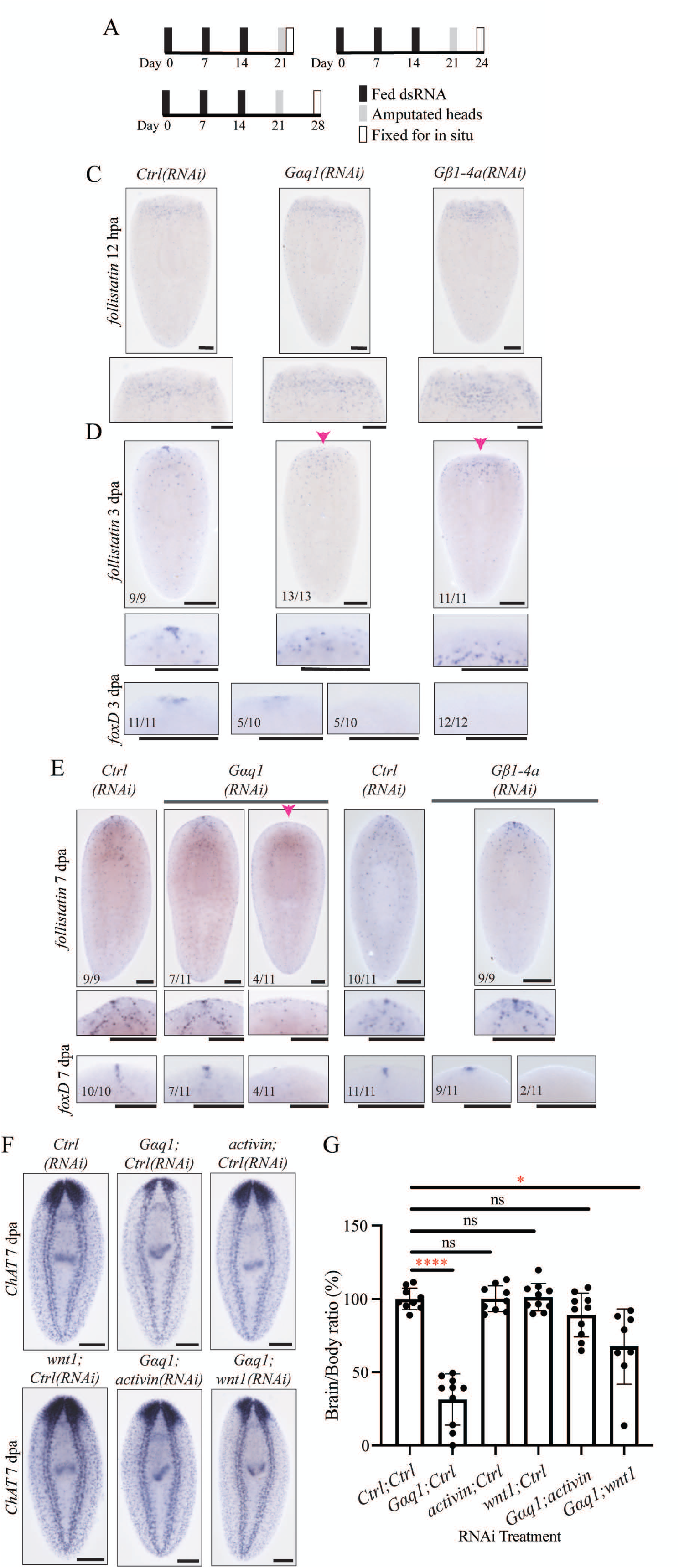
*Gαq1* supports head regeneration through production of *follistatin*^+^ anterior pole cells. (**A**) RNAi paradigms used for data presented 12 hpa (top left), 3 dpa (top right), and 7 dpa (bottom left). Representative images of *follistatin* expression at 12 hpa (**C**) and *follistatin* and *foxD* expression at 3 dpa **(D)** and 7 dpa **(E)** in *Gαq1(RNAi)* and *Gβ1-4a(RNAi)* animals. Magenta arrowheads indicate absent anterior pole expression domain. (**F**) Representative images showing *ChAT* expression from rescue experiments 7 dpa. (**G**) Bar graph showing results from quantification of brain/body ratios in rescue experiments, using the same RNAi paradigm and brain quantification method described in Fig. 2. * = P value ≤ 0.05 and **** = P value ≤ 0.0001. Scale = 200 μm.

To determine whether *Gαq1* and *Gβ1-4a* support *follistatin* expression in the anterior pole, we examined *follistatin* expression during pole formation. At 3 dpa, *Gαq1(RNAi)* and *Gβ1-4a(RNAi)* animals had reduced or absent *follistatin* expression at the anterior pole (Fig. 6D). At 7 dpa, ∼36% of *Gαq1(RNAi)* animals still lacked *follistatin*^*+*^ anterior pole cells (Fig. 6E). However, most *Gβ1-4a(RNAi)* animals established *follistatin*^*+*^ pole cells by 7 dpa (Fig. 6E), reflecting a similar delay in anterior pole formation seen with other markers (Fig. 5E and Supplemental Fig. S5).

Both *notum* and *follistatin* expression in anterior pole cell progenitors relies on a key transcription factor, encoded by *foxD* (Roberts-Galbraith and Newmark 2013; Scimone et al. 2014; Vogg et al. 2014). To investigate whether *Gαq*1 could modulate *follistatin* through FoxD, we examined *foxD* expression following *Gαq1* knockdown. Indeed, anterior *foxD* expression was absent in 50% of *Gαq1(RNAi)* animals at 3 dpa and ∼36% of animals at 7 dpa (Fig. 6D-E; Supplemental Fig. S5D). We confirmed this significant reduction of *foxD* expression through RT-qPCR (Supplemental Fig. S5E). *Gβ1-4a(RNAi)* animals displayed absent *foxD* anterior pole expression 3 dpa and most animals resumed *foxD* expression by 7 dpa (Fig. 6D-E). Thus, our data suggest that impacts on *follistatin* expression could be mediated by *foxD*.

Previous work characterizing the Follistatin/Activin and Notum/Wnt pathways determined that reduction of the antagonistic posterior-promoting ligands rescued head regeneration (Petersen and Reddien 2011; Roberts-Galbraith and Newmark 2013; Gaviño et al. 2013). The similarities between *Gαq1(RNAi)* and *follistatin(RNAi)* phenotypes and the impact of *Gαq1(RNAi)* on *follistatin* expression led us to hypothesize that *Gαq1* functions to promote Follistatin signaling from the pole. To test this hypothesis, we performed RNAi targeting *Gαq1* with *activin(RNAi), wnt1(RNAi)*, or *bmp4(RNAi)* (a TGF-beta ligand that impacts dorsoventral polarity) (Reddien et al. 2007; Roberts-Galbraith and Newmark 2013; Gaviño et al. 2013; Tewari et al. 2018). *activin(RNAi)* significantly rescued *Gαq1(RNAi)*-induced brain regeneration defects (Fig. 6F-G). *wnt1(RNAi)* also partially rescued *Gaq1(RNAi)* (Fig. 6F-G). As expected, *bmp4(RNAi)* failed to rescue regeneration in *Gαq1(RNAi)* animals (Supplemental Fig. S6B-C). Incidentally, though we primarily focused on a functional connection between *Gαq*1 and Follistatin, we also found that *activin* inhibition modestly restored brain regeneration in *Gβ1-4a(RNAi)* animals (Supplemental Fig. S6D-E). We conclude that *Gαq*1 function specifically supports Follistatin signaling from the anterior pole during head regeneration.

## Discussion

The overwhelming number of GPCRs hinders progress in understanding the function of this fascinating receptor family in planarian regeneration and stem cell biology. In this work, we take a first step toward investigating planarian GPCR signaling by identifying and functionally characterizing the heterotrimeric G protein subunit complement in behavior and regeneration. We characterized 38 heterotrimeric G protein homologs, of which 23 were well enough conserved to categorize. Through our functional screens, we identified 5 subunit genes required for proper planarian movement (Fig. 7). Using available single cell sequencing data, we applied our work with G protein subunits and identified a putative serotonin receptor (GCR052) that could function with Gβx2 in locomotion. Through our brain regeneration screen, we identified 7 genes with roles in regeneration, with *Gαq1* and *Gβ1-4a* having especially significant effects (Fig. 7). We determined that *Gαq1* and *Gβ1-4a* promote the regeneration and establishment speed of the anterior pole, respectively. Our findings reveal new mechanisms active in planarian regeneration and behavior and support the hypothesis that GPCR pathways are likely to be involved in signaling events that drive and coordinate planarian regeneration.

**Figure 7.**
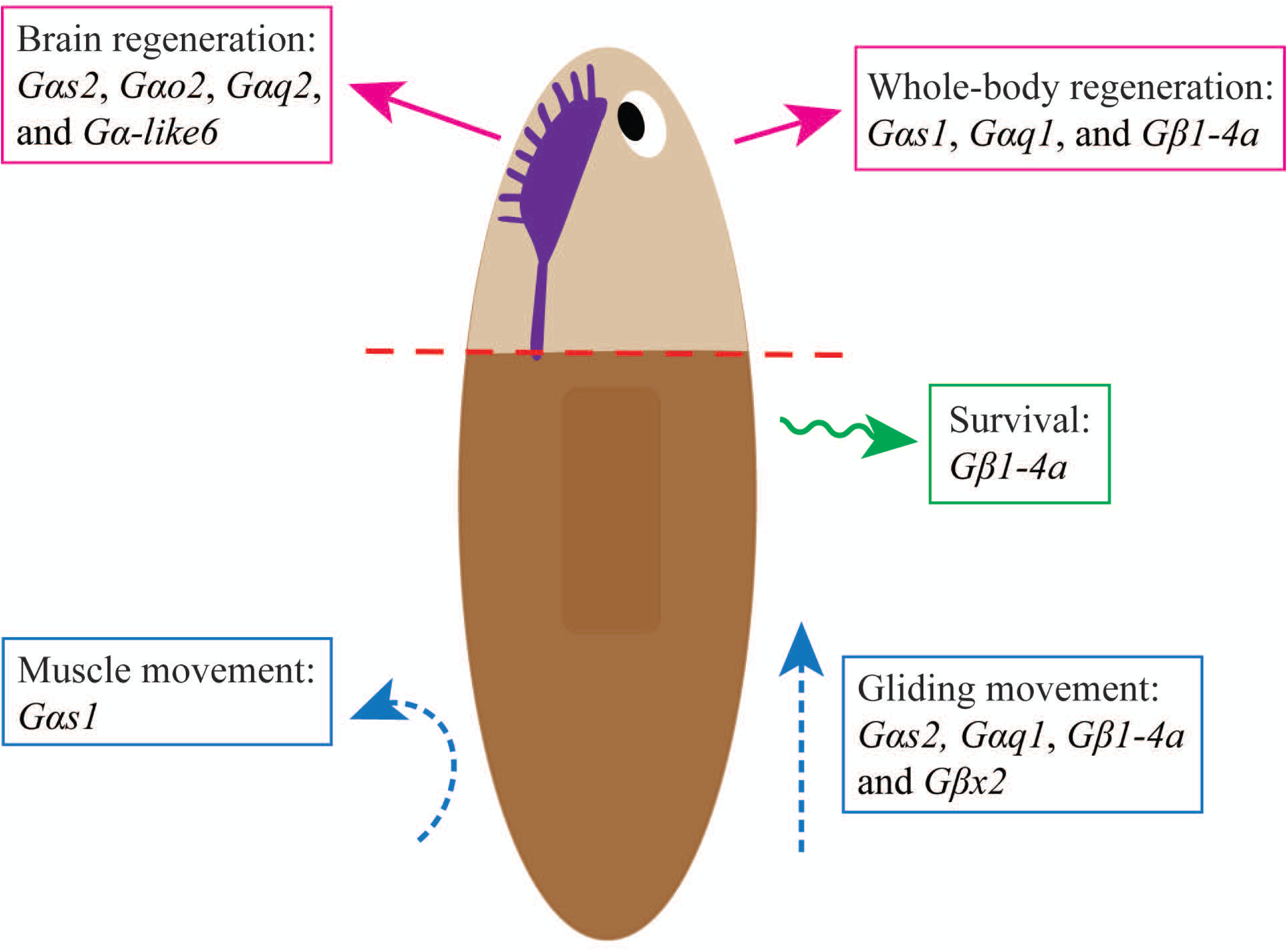
Planarian heterotrimeric G proteins play diverse roles in regeneration, physiology, and behavior. Graphical summary of roles described in this work for heterotrimeric G proteins in *S. mediterranea* (for details, see Figs. 2-3).

Due to the functionally similar but divergent effects of *Gαq1* or *Gβ1-4a*, we reason that these subunits could be activated downstream of a common GPCR but then activate different downstream pathways to support tissue regeneration (Brock et al. 2003; Tang and Gilman 1991; Inglese et al. 1995). This model is supported by coexpression of *Gαq1* or *Gβ1-4a* in subsets of muscle cells and neurons in single cell sequencing data (Fincher et al. 2018) (Supplemental Table S1 and S3). Additionally, we show through RT-qPCR that while targeting *Gβ1-4a* does not impact expression of *Gαq1*, expression of *Gβ1-4a* is significantly reduced in *Gαq1(RNAi)* animals (Supplemental Fig. S7).

### Gαq1 provides a putative connection between planarian GPCR signaling and defined polarity axes

The phenotypic similarities between *Gαq1(RNAi)* and *follistatin(RNAi)* animals and the ability to rescue by *activin* double knockdown indicate that the *Gαq*1 protein likely cooperates with Follistatin during regeneration. Our results suggest that Gαq1 could function upstream to promote *follistatin* expression at the anterior pole. Alternatively, Activin signals belong to the transforming growth factor-β (TGF-β) family, and recent work describes the potential for GPCRs to modulate TGF-β pathways through transactivation (Burch et al. 2012; Hinck et al. 2016; Schafer and Blaxall 2017). Therefore, Gαq1 could potentially influence the Activin/Follistatin axis through a noncanonical mechanism. Further exploring relationships between *Gαq1* and pathway components will help define the nature of the Gαq1/Follistatin cooperation.

Additionally, because *Gαq1(RNAi)* animals displayed functional wound-induced *follistatin* expression, our results also support the notion that *follistatin* expression from the anterior pole is needed to drive successful head regeneration (Roberts-Galbraith and Newmark 2013; Gaviño et al. 2013; Tewari et al. 2018). Therefore, results from future work with *Gαq1* could inform the nature of anterior identity establishment. If *Gαq1*’s mechanism somehow influences the ability of cells to perceive or respond to Activin ligands, it could suggest more complex molecular processes for anterior establishment. Potential roles for *Gαq1* (and GPCRs) in modulating the Activin pathway and promoting polarity reestablishment will require further investigation.

### Identification of gcr052 increases our understanding of the complexity of serotonin’s role in planarian locomotion

In addition to characterizing planarian heterotrimeric G proteins with roles in regeneration, this work also contributes to knowledge of mechanisms governing planarian movement. The current model for planarian gliding is that serotonergic neurons directly innervate ventral epidermal cells and coordinate the beating of motile cilia (Currie and Pearson 2013; März et al. 2013). Furthermore, experiments with mianserin, a pharmacological inhibitor of serotonin receptors, also implicated GPCRs in cilia coordination (Kuang et al. 2002; Currie and Pearson 2013). In this work, we identify 2 G protein subunits (*Gαs2* and *Gβx2*) that affect locomotion in *S. mediterranea* (Fig. 2). We further identified *gcr052* (Saberi et al. 2016), which encodes a putative serotonin GPCR, as a potential specific mediator of gliding motion. This receptor also appears to be *ser85* (from [Zamanian 2011]), supporting the validity of both approaches.

*gcr052* is expressed in *rootletin*^+^ (Glazer et al. 2010) ciliated epidermal cells (Fincher et al. 2018), supporting the model that serotonin directly influences ciliary coordination (Supplemental Table S3). However, the heterotrimeric G protein subunit that phenocopies the inching phenotype (*Gβx2*) is not enriched in epidermal cell clusters, but instead is coexpressed with *gcr052* in neurons (Fincher et al. 2018) (Supplemental Table S3). Although we cannot rule out undetected epidermal transcripts or protein for *Gβx2*, our results indicate that the serotonin signaling to other neurons may also be important for planarian locomotion.

Additional assays may reveal further roles of heterotrimeric G proteins in behavior and sensation. These proteins act in diverse biological processes, such as sensation, in other animals (Yarfitz and Hurley 1994; Jones and Reed 1989; Wong et al. 1996). Additionally, 8 planarian G protein subunits show expression enrichment in proposed sensory structures called the brain branches (Agata et al. 1998; Okamoto et al. 2005), further supporting this notion (Fig. 1; Supplemental Fig. S8). Using the G protein group as a primary screening strategy may be a beneficial starting point for future study of GPCRs in planarian sensory neurobiology.

### Planarian heterotrimeric G proteins can suggest candidate receptors for future planarian GPCR research

Because GPCRs represent one of the largest receptor families in many organisms, including humans (Fredriksson et al. 2003) and planarians (Zamanian et al. 2011; Saberi et al. 2016), approaches to narrow relevant GPCRs can prove to be valuable. Our investigation into planarian heterotrimeric G protein subunits produced functionally distinct and measurable phenotypes, supporting the idea that planarian heterotrimeric G protein subunits could provide a practical approach to identify and study roles of GPCRs.

To develop a G protein subunit-driven candidate approach, we formulated a pipeline that identifies candidate GPCR genes using phenotypes from our work along with published single cell sequencing datasets (Fig. 2D; Supplemental Table S3) (Fincher et al. 2018; Swapna et al. 2018; Plass et al. 2018). Our work with *Gβx2* and *gcr052* demonstrates the utility of characterizing heterotrimeric G proteins as a first step to identifying the associated GPCR and understanding the cellular mechanism (Wise et al. 2004; Oh et al. 2006; Civelli et al. 2013; Ngo et al. 2016). In the future we plan to apply this approach to narrow candidate GPCRs that work with relevant heterotrimeric G proteins to promote polarity establishment and successful regeneration. Identification of novel signaling pathways with key roles in regeneration will help us understand how information about injury is converted into cellular responses to coordinate and drive regeneration.

## Materials and Methods

### Animal Maintenance

Planarians from an asexual strain of the species *S. mediterranea* (CIW4; [Alvarado et al. 2002]) were kept in 1X Monjuïc salts (1.6 mmol/L NaCl, 1 mmol/L CaCl_2_, 1 mmol/L MgSO_4_, 0.1 mmol/L MgCl_2_, 0.1 mmol/L KCl and 1.2 mmol/L NaHCO_3_ prepared in ELGA PURELAB [ELGA LabWater, Woodridge, IL] ultrapure water) (Cebrià and Newmark 2005) at 18°C in the dark. Animals were fed beef liver puree weekly or biweekly. Animals were cut periodically to expand their numbers and generate properly sized (∼2-5 mm) individuals for experiments. Animals were starved for a minimum of one week before experiments.

### Gene Identification

Gα subunit-like transcripts were mined using the Guanine nucleotide-binding domain (PF00503 [Coleman et al. 1994]), Gβ subunit-like transcripts were mined using the WD40-repeat-containing domain preceded by N-terminal alpha helix (IPR001632 [Wall et al. 1995]), and Gγ subunit-like transcripts were mined using the GGL domain (PF00631 [Snow et al. 1998]). Each relevant functional domain (from Pfam [El-Gebali et al. 2019] or InterPro [Mitchell et al. 2019]) was searched within the translated *S. mediterranea* transcripts dataset dd_Smed_v6 (Brandl et al. 2016; Rozanski et al. 2019), then redundant transcripts were removed. To ensure the retrieved Gγ subunit-like sequences were not Regulator of G protein Signaling (RGS) proteins, the absence of an RGS domain (PF00615 [Chen et al. 2001; Longenecker et al. 2001]) was validated (Supplemental Table S1).

### Protein alignment and Phylogenetic Analysis

Amino acid sequences were predicted using the web-based translation tool at the Swiss ExPASy (Expert Protein Analysis System) Molecular Biology Server (Swiss Institute of Bioinformatics, University of Lausanne, Switzerland) (Gasteiger et al. 2003). Protein sequences were aligned to reference sequences from other animals (Supplemental Table S4) using Clustal Omega O(1.2.4) (Sievers and Higgins 2014) and secondary structures were predicted with ESpript3.0 (Robert and Gouet 2014), using well-characterized structure examples (PDB ID: 1GP2). Phylogeny was analyzed using www.phylogeny.fr (Dereeper et al. 2008). The “A la Carte” option was selected with MUSCLE for alignment (Edgar 2004) and PhyML for construction of the phylogenetic tree (Guindon et al. 2010). For the PhyML analysis, 100 bootstrap replicates were performed, and the WAG model of amino acid substitution was applied.

### Molecular cloning

For genes of interest, primers were designed using Primer3 (Rozen and Skaletsky) to amplify an ∼700 bp region of the corresponding gene from asexual *S. mediterranea* cDNA (Supplemental Table S5). The PCR products were then cloned into the vector, pJC53.2 (Collins et al. 2010) using standard molecular biology protocols.

### RNA interference (RNAi) experiments

dsRNA was transcribed in vitro from PCR products amplified from pJC53.2 using standard molecular methods (Collins et al. 2010; Rouhana et al. 2013). Concentration of dsRNA was determined using either a Nanophotometer NP80 (Implen, Munich, Germany) or by band intensity after gel electrophoresis. For a typical experiment, 10-12 animals were fed 1-3μg dsRNA mixed in ∼30μL food (beef liver paste, 4:1 liver:salts mixture), and 1 μL green food dye was added to validate that the animals ate. The mixture was doubled for larger experiments. Negative control worms were fed dsRNA matching *green fluorescent protein* (*GFP)* or bacterial genes (*Chloramphenicol resistance gene* [*Cm*^*R*^] and *toxin CcdB* [*ccdB]*). Animals were kept in 60-100 mm Petri dishes. After eating, the animals were washed, transferred to fresh dishes, and the liquid was supplemented with 1:1000 gentamicin sulfate (50 mg/mL stock [Gemini Bio, West Sacramento, CA]). Animals were fed dsRNA ∼once per week for three total feedings (more feedings given in long-term RNAi experiments [Fig. 3 and 4]) then were processed. Live images during experiments were obtained using a Zeiss Axiocam 506 Color camera mounted on a Zeiss Axio ZoomV.16 microscope (ZEISS Microscopy, Jena, Germany). Live images and video were also captured on an iPhone 6 and processed in iMovie (Apple Inc., Cupertino, California). For the flipping behavior assay (Fig. 3B-C), recordings were taken of each individual live animal for up to 5 minutes after being put on their dorsal side and observed for the time it took the animals flip to their ventral side.

### In situ hybridization (ISH)

Single-stranded antisense riboprobes were transcribed with Digoxygenin (Dig-11-UTP) (Sigma-Aldrich, St. Louis, MO) using standard molecular methods (Collins et al. 2010). Animals were fixed, hybridized with riboprobes, and stained as previously described (King and Newmark 2013), with the following modifications: animals were killed in a 10% N-Acetyl Cysteine solution and treated with a 2 μg/mL Proteinase K solution. Regenerating animals were treated with the Proteinase K/post fixation steps (as opposed to a boiling step). After the hybridization step, 56 °C washes were as follows: one 20-minute Wash hyb (25% Formamide, 3.5X SSC, 0.525 M NaCl, 0.0525 M Na Citrate, 0.1% Triton X-100, and pH 7.0) wash, three 20-minute 2X SSCx (2X SSC [0.3 M NaCl, 0.03 M Na Citrate, pH 7.0] and 0.1% Triton X-100) washes, four 20-minute 0.2X SSCx (0.2X SSC and 0.1% Triton X-100) washes. We also replaced MABT with TNTx (0.1 M Tris pH 7.5, 0.15 M NaCl, and 0.3% Triton X-100). After antibody incubation, animals were washed in TNTx for 5 minutes (1 wash), 10 minutes (1 wash), and six 20-minute washes. The fixation step after sample development was omitted. Other key reagents include anti-digoxygenin conjugated with an alkaline phosphatase (Anti-Dig-AP 1:2000 dilution), nitro-blue tetrazolium (NBT), and 5-bromo-4-chloro-3’-indolyphosphate (BCIP) (all Sigma-Aldrich, St. Louis, MO). Animals were mounted in 80% glycerol and imaged with a Zeiss Axiocam 506 Color camera mounted on a Zeiss Axio Zoom. V16 microscope (ZEISS Microscopy, Jena, Germany).

### Image Quantification

For regeneration assays, the normalized areas of the brains (Figs) or blastemas (figs) were measured from fixed sample images by tracing the structures with FIJI (Fiji is just ImageJ) imaging software (Schindelin et al. 2012), as described previously (Roberts-Galbraith et al. 2016). For growth assays, animal lengths were measured from live images in FIJI. Data were statistically analyzed and visualized using Prism -GraphPad Version 7.0 software (GraphPad Software, San Diego, CA). Tests employed were Unpaired T-Test with Welch’s Correction (Fig. 3C), one-way ANOVA (Fig. 2E; 4D), and Brown-Forsythe and Welch ANOVA (Fig. 6G).

### Quantitative Reverse Transcription Polymerase Chain Reaction (RT-qPCR)

RNA was extracted from animals using Trizol Reagent (Thermo Fisher Scientific, Waltham, MA) as per the manufacturer’s protocol (Liu and Rink 2018). Samples were treated with RQ1 RNase-free DNase (Promega Corporation, Madison, WI) for 15 minutes at 37°C. cDNA was synthesized from RNA using an iScript kit (Bio-Rad, Hercules, CA). RT-qPCR reactions were completed using SYBR Green PCR Master Mix (Bio-Rad, Hercules, CA) in a QuantStudio® 3 real-time PCR system (Applied Biosystems, Foster City, CA). Primers were generated by Primer3 (Rozen and Skaletsky) and targeted sequences ∼100bp in length (Supplemental Table S5). RT-qPCR primers were designed to match a region of the transcripts not included in dsRNA constructs using Benchling software (Benchling, San Francisco, CA). Transcript abundance for genes of interest was normalized using the control gene, *β tubulin* (Collins et al. 2010). Experiments were performed in biological and technical triplicate (n=12 animals per biological replicate). Data were statistically analyzed and visualized using Prism -GraphPad Version 7.0 software (GraphPad Software, San Diego, CA).

## Acknowledgements

We thank Bidushi Chandra, Kendall Clay, Samantha Hack, and Taylor Medlock-Lanier for feedback on this manuscript. We also thank Britessia Smith for technical support and helpful discussions. We thank Alejandro Sánchez Alvarado and Shane Merryman (Stowers Institute) for providing planarians and for instructions on recirculation system construction. Finally, we thank White Oak Pastures (Bluffton, GA) for beef liver. This work was supported by the University of Georgia start-up funding, NSF (IOS-1942822), the McKnight Foundation, and the Alfred P. Sloan Foundation.

## Author Contributions

Conceived and designed the experiments: JEJ and RRG. Performed the experiments: JEJ. Analyzed and interpreted the data: JEJ and RRG. Wrote the manuscript: JEJ and RRG.

## References

Agata K, Soejima Y, Kato K, Kobayashi C, Umesono Y, Watanabe K. 1998. Structure of the Planarian Central Nervous System (CNS) Revealed by Neuronal Cell Markers. Zoological Science 15: 433–440.

Alvarado AS, Newmark PA, Robb SMC, Juste R. 2002. The Schmidtea mediterranea database as a molecular resource for studying platyhelminthes, stem cells and regeneration. Development 129: 5659–5665.

Anantharaman V, Abhiman S, de Souza RF, Aravind L. 2011. Comparative genomics uncovers novel structural and functional features of the heterotrimeric GTPase signaling system. Gene 475: 63–78.

Baguna J, Salo E, Auladell C. 1989. Regeneration and pattern formation in planarians. III. that neoblasts are totipotent stem cells and the cells. Development 107: 77–86.

Bates CA, Meyer RL. 1996. Heterotrimeric G protein activation rapidly inhibits outgrowth of optic axons from adult and embryonic mouse, and goldfish retinal explants. Brain Research 714: 65–75.

Brandl H, Moon H, Vila-Farré M, Liu S-Y, Henry I, Rink JC. 2016. PlanMine – a mineable resource of planarian biology and biodiversity. Nucleic Acids Research 44: D764–D773.

Brock C, Schaefer M, Reusch HP, Czupalla C, Michalke M, Spicher K, Schultz G, Nu□rnberg B. 2003. Roles of Gβγ in membrane recruitment and activation of p110γ/p101 phosphoinositide 3-kinase γ. Journal of Cell Biology 160: 89–99.

Burch ML, Osman N, Getachew R, Al-aryahi S, Poronnik P, Zheng W, Hill MA, Little PJ. 2012. G protein coupled receptor transactivation: Extending the paradigm to include serine/threonine kinase receptors. The International Journal of Biochemistry & Cell Biology 44: 722–727.

Cebrià F, Kudome T, Nakazawa M, Mineta K, Ikeo K, Gojobori T, Agata K. 2002. The expression of neural-specific genes reveals the structural and molecular complexity of the planarian central nervous system. Mechanisms of Development 116: 199–204.

Cebrià F, Newmark PA. 2005. Planarian homologs of netrin and netrin receptor are required for proper regeneration of the central nervous system and the maintenance of nervous system architecture. Development 132: 3691–3703.

Chen Z, Wells CD, Sternweis PC, Sprang SR. 2001. Structure of the rgRGS domain of p115RhoGEF. Nature Structural Biology 8: 805–809.

Choi HY, Saha SK, Kim K, Kim S, Yang G-M, Kim B, Kim J-H, Cho S-G. 2015. G protein-coupled receptors in stem cell maintenance and somatic reprogramming to pluripotent or cancer stem cells. BMB Reports 48: 68–80.

Civelli O, Reinscheid RK, Zhang Y, Wang Z, Fredriksson R, Schiöth HB. 2013. G Protein–Coupled Receptor Deorphanizations. Annual Review of Pharmacology and Toxicology 53: 127–146.

Cloutier JK, McMann CL, Oderberg IM, Reddien PW. 2021. activin-2 is required for regeneration of polarity on the planarian anterior-posterior axis. PLOS Genetics 17: e1009466.

Coleman D, Berghuis A, Lee E, Linder M, Gilman A, Sprang. 1994. Structures of active conformations of Gi alpha 1 and the mechanism of GTP hydrolysis. Science 265: 1405–1412.

Collins JJ, Hou X, Romanova E v., Lambrus BG, Miller CM, Saberi A, Sweedler J v., Newmark PA. 2010. Genome-Wide Analyses Reveal a Role for Peptide Hormones in Planarian Germline Development. PLoS Biology 8: e1000509.

Currie KW, Pearson BJ. 2013. Transcription factors lhx1/5-1 and pitx are required for the maintenance and regeneration of serotonergic neurons in planarians. Development 140: 3577–3588.

Dereeper A, Guignon V, Blanc G, Audic S, Buffet S, Chevenet F, Dufayard J-F, Guindon S, Lefort V, Lescot M, et al. 2008. Phylogeny.fr: robust phylogenetic analysis for the non-specialist. Nucleic Acids Research 36: W465–W469.

Doze VA, Perez DM. 2013. GPCRs in Stem Cell Function. pp. 175–216.

Edgar RC. 2004. MUSCLE: multiple sequence alignment with high accuracy and high throughput. Nucleic Acids Research 32: 1792–1797.

Eisenhoffer GT, Kang H, Alvarado AS. 2008. Molecular Analysis of Stem Cells and Their Descendants during Cell Turnover and Regeneration in the Planarian Schmidtea mediterranea. Cell Stem Cell 3: 327–339.

El-Gebali S, Mistry J, Bateman A, Eddy SR, Luciani A, Potter SC, Qureshi M, Richardson LJ, Salazar GA, Smart A, et al. 2019. The Pfam protein families database in 2019. Nucleic Acids Research 47: D427–D432.

Fincher CT, Wurtzel O, de Hoog T, Kravarik KM, Reddien PW. 2018. Cell type transcriptome atlas for the planarian Schmidtea mediterranea. Science 360.

Fredriksson R, Lagerström MC, Lundin L-G, Schiöth HB. 2003. The G-Protein-Coupled Receptors in the Human Genome Form Five Main Families. Phylogenetic Analysis, Paralogon Groups, and Fingerprints. Molecular Pharmacology 63: 1256–1272.

Garland SL. 2013. Are GPCRs Still a Source of New Targets? Journal of Biomolecular Screening 18: 947–966.

Gasteiger E, Gattiker A, Hoogland C, Ivanyi I, Appel RD, Bairoch A. 2003. ExPASy: The proteomics server for in-depth protein knowledge and analysis. Nucleic acids research 31: 3784–8.

Gaviño MA, Wenemoser D, Wang IE, Reddien PW. 2013. Tissue absence initiates regeneration through Follistatin-mediated inhibition of Activin signaling. eLife 2.

Glazer AM, Wilkinson AW, Backer CB, Lapan SW, Gutzman JH, Cheeseman IM, Reddien PW. 2010. The Zn Finger protein Iguana impacts Hedgehog signaling by promoting ciliogenesis. Developmental Biology 337: 148–156.

Guindon S, Dufayard J-F, Lefort V, Anisimova M, Hordijk W, Gascuel O. 2010. New Algorithms and Methods to Estimate Maximum-Likelihood Phylogenies: Assessing the Performance of PhyML 3.0. Systematic Biology 59: 307–321.

Guo H, Du X, Zhang Y, Wu J, Wang C, Li M, Hua X, Zhang XA, Yan J. 2019. Specific miRNA-G Protein-Coupled Receptor Networks Regulate Sox9a/Sox9b Activities to Promote Gonadal Rejuvenation in Zebrafish. Stem Cells 37: 1189–1199.

Gurley KA, Rink JC, Alvarado AS. 2008. β-Catenin Defines Head Versus Tail Identity During Planarian Regeneration and Homeostasis. Science 319: 323–327.

Hinck AP, Mueller TD, Springer TA. 2016. Structural Biology and Evolution of the TGF-β Family. Cold Spring Harbor Perspectives in Biology 8: a022103.

Hopkins AL, Groom CR. 2002. The druggable genome. Nature Reviews Drug Discovery 1: 727–730.

Iglesias M, Almuedo-Castillo M, Aboobaker AA, Saló E. 2011. Early planarian brain regeneration is independent of blastema polarity mediated by the Wnt/β-catenin pathway. Developmental Biology 358: 68–78.

Iglesias M, Gomez-Skarmeta JL, Saló E, Adell T. 2008. Silencing of Smed -β catenin1 generates radial-like hypercephalized planarians. Development 135: 1215–1221.

Inglese J, Koch W, Touhara K, Lefkowitz R. 1995. Gβγ interactions with PH domains and Ras-MAPK signaling pathways. Trends in Biochemical Sciences 20: 151–156.

Jansen G, Thijssen KL, Werner P, van derHorst M, Hazendonk E, Plasterk RHA. 1999. The complete family of genes encoding G proteins of Caenorhabditis elegans. Nature Genetics 21: 414–419.

Jones DT, Reed RR. 1989. G olf[: an Olfactory Neuron Specific-G Protein Involved in Odorant Signal Transduction. Science 244: 790–795.

Kakugawa S, Langton PF, Zebisch M, Howell SA, Chang T-H, Liu Y, Feizi T, Bineva G, O’Reilly N, Snijders AP, et al. 2015. Notum deacylates Wnt proteins to suppress signalling activity. Nature 519: 187–192.

King RS, Newmark PA. 2013. In situ hybridization protocol for enhanced detection of gene expression in the planarian Schmidtea mediterranea. BMC Developmental Biology 13: 8.

Kiseleva EV, Sidorova MV, Gorbacheva LR, Strukova SM. 2014. Peptide-agonist of protease-activated receptor (PAR 1), similar to activated protein C, promotesproliferation in keratinocytes and wound healing of epithelial layer. Biomeditsinskaya Khimiya 60: 702–706.

Krishnan A, Almén MS, Fredriksson R, Schiöth HB. 2012. The Origin of GPCRs: Identification of Mammalian like Rhodopsin, Adhesion, Glutamate and Frizzled GPCRs in Fungi. PLoS ONE 7: e29817.

Kuang S, Doran SA, Wilson RJA, Goss GG, Goldberg JI. 2002. Serotonergic sensory-motor neurons mediate a behavioral response to hypoxia in pond snail embryos. Journal of Neurobiology 52: 73–83.

Labbé RM, Irimia M, Currie KW, Lin A, Zhu SJ, Brown DDR, Ross EJ, Voisin V, Bader GD, Blencowe BJ, et al. 2012. A Comparative Transcriptomic Analysis Reveals Conserved Features of Stem Cell Pluripotency in Planarians and Mammals. STEM CELLS 30: 1734–1745.

Lagerström MC, Schiöth HB. 2008. Structural diversity of G protein-coupled receptors and significance for drug discovery. Nature Reviews Drug Discovery 7: 339–357.

Langenhan T, Barr MM, Bruchas MR, Ewer J, Griffith LC, Maiellaro I, Taghert PH, White BH, Monk KR. 2015. Model Organisms in G Protein–Coupled Receptor Research. Molecular Pharmacology 88: 596–603.

Lapan SW, Reddien PW. 2012. Transcriptome Analysis of the Planarian Eye Identifies ovo as a Specific Regulator of Eye Regeneration. Cell Reports 2: 294–307.

Li S, Yang C, Zhang L, Gao X, Wang X, Liu W, Wang Y, Jiang S, Wong YH, Zhang Y, et al. 2016. Promoting axon regeneration in the adult CNS by modulation of the melanopsin/GPCR signaling. Proceedings of the National Academy of Sciences 113: 1937–1942.

Liu S-Y, Rink JC. 2018. Total RNA Isolation from Planarian Tissues. pp. 259–265.

Longenecker KL, Lewis ME, Chikumi H, Gutkind JS, Derewenda ZS. 2001. Structure of the RGS-like Domain from PDZ-RhoGEF. Structure 9: 559–569.

Lozano BC. 2015. The Identification and the Functional Validation of Eye Development and Regeneration Genes in Schmidtea Mediterranea.

Malpe MS, McSwain LF, Kudyba K, Ng CL, Nicholson J, Brady M, Qian Y, Choksi V, Hudson AG, Parrott BB, et al. 2020. G-protein signaling is required for increasing germline stem cell division frequency in response to mating in Drosophila males. Scientific Reports 10: 3888.

März M, Seebeck F, Bartscherer K. 2013. A Pitx transcription factor controls the establishment and maintenance of the serotonergic lineage in planarians. Development 140: 4499–4509.

Mitchell AL, Attwood TK, Babbitt PC, Blum M, Bork P, Bridge A, Brown SD, Chang H-Y, El-Gebali S, Fraser MI, et al. 2019. InterPro in 2019: improving coverage, classification and access to protein sequence annotations. Nucleic Acids Research 47: D351–D360.

Mo J-S. 2017. The role of extracellular biophysical cues in modulating the Hippo-YAP pathway. BMB Reports 50: 71–78.

Nakamura T, Takio K, Eto Y, Shibai H, Titani K, Sugino H. 1990. Activin-Binding Protein from Rat Ovary Is Follistatin. Science 247: 836–838.

Ngo T, Kufareva I, Coleman JL, Graham RM, Abagyan R, Smith NJ. 2016. Identifying ligands at orphan GPCRs: current status using structure-based approaches. British journal of pharmacology 173: 2934–51.

O’Connor JT, Stevens AC, Shannon EK, Akbar FB, LaFever KS, Narayanan NP, Gailey CD, Hutson MS, Page-McCaw A. 2021. Proteolytic activation of Growth-blocking peptides triggers calcium responses through the GPCR Mthl10 during epithelial wound detection. Developmental Cell 56: 2160–2175.e5.

Oh DY, Kim K, Kwon HB, Seong JY. 2006. Cellular and Molecular Biology of Orphan G Protein[Coupled Receptors. pp. 163–218.

Okamoto K, Takeuchi K, Agata K. 2005. Neural Projections in Planarian Brain Revealed by Fluorescent Dye Tracing. Zoological Science 22: 535–546.

Oldham WM, Hamm HE. 2008. Heterotrimeric G protein activation by G-protein-coupled receptors. Nature Reviews Molecular Cell Biology 9: 60–71.

Owlarn S, Bartscherer K. 2016. Go ahead, grow a head! A planarian’s guide to anterior regeneration. Regeneration 3: 139–155.

Pascual-Carreras E, Sureda-Gómez M, Barrull-Mascaró R, Jordà N, Gelabert M, Coronel-Córdoba P, Saló E, Adell T. 2021. WNT-FRIZZLED-LRP5/6 Signaling Mediates Posterior Fate and Proliferation during Planarian Regeneration. Genes 12: 101.

Petersen CP, Reddien PW. 2009a. A wound-induced Wnt expression program controls planarian regeneration polarity. Proceedings of the National Academy of Sciences 106: 17061–17066.

Petersen CP, Reddien PW. 2011. Polarized notum Activation at Wounds Inhibits Wnt Function to Promote Planarian Head Regeneration. Science 332: 852–855.

Petersen CP, Reddien PW. 2008. Smed-β catenin-1 Is Required for Anteroposterior Blastema Polarity in Planarian Regeneration. Science 319: 327–330.

Petersen CP, Reddien PW. 2009b. Wnt Signaling and the Polarity of the Primary Body Axis. Cell 139: 1056–1068.

Pierce KL, Premont RT, Lefkowitz RJ. 2002. Seven-transmembrane receptors. Nature Reviews Molecular Cell Biology 3: 639–650.

Plass M, Solana J, Wolf FA, Ayoub S, Misios A, Glažar P, Obermayer B, Theis FJ, Kocks C, Rajewsky N. 2018. Cell type atlas and lineage tree of a whole complex animal by single-cell transcriptomics. Science 360.

Raz AA, Wurtzel O, Reddien PW. 2021. Planarian stem cells specify fate yet retain potency during the cell cycle. Cell Stem Cell 28: 1307–1322.e5.

Reddien PW. 2018. The Cellular and Molecular Basis for Planarian Regeneration. Cell 175: 327–345.

Reddien PW, Bermange AL, Kicza AM, Sánchez Alvarado A. 2007. BMP signaling regulates the dorsal planarian midline and is needed for asymmetric regeneration. Development 134: 4043–4051.

Reddien PW, Oviedo NJ, Jennings JR, Jenkin JC, Alvarado AS. 2005. SMEDWI-2 Is a PIWI-Like Protein That Regulates Planarian Stem Cells. Science 310: 1327–1330.

Robert X, Gouet P. 2014. Deciphering key features in protein structures with the new ENDscript server. Nucleic Acids Research 42: W320–W324.

Roberts-Galbraith RH, Brubacher JL, Newmark PA. 2016. A functional genomics screen in planarians reveals regulators of whole-brain regeneration. eLife 5.

Roberts-Galbraith RH, Newmark PA. 2013. Follistatin antagonizes Activin signaling and acts with Notum to direct planarian head regeneration. Proceedings of the National Academy of Sciences 110: 1363–1368.

Rouhana L, Weiss JA, Forsthoefel DJ, Lee H, King RS, Inoue T, Shibata N, Agata K, Newmark PA. 2013. RNA interference by feeding in vitro–synthesized double[stranded RNA to planarians: Methodology and dynamics. Developmental Dynamics 242: 718–730.

Rozanski A, Moon H, Brandl H, Martín-Durán JM, Grohme MA, Hüttner K, Bartscherer K, Henry I, Rink JC. 2019. PlanMine 3.0—improvements to a mineable resource of flatworm biology and biodiversity. Nucleic Acids Research 47: D812–D820.

Rozen S, Skaletsky H. Primer3 on the WWW for General Users and for Biologist Programmers. In Bioinformatics Methods and Protocols, pp. 365–386, Humana Press, New Jersey.

Saberi A, Jamal A, Beets I, Schoofs L, Newmark PA. 2016. GPCRs Direct Germline Development and Somatic Gonad Function in Planarians. PLOS Biology 14: e1002457.

Schafer AE, Blaxall BC. 2017. G Protein Coupled Receptor-mediated Transactivation of Extracellular Proteases. Journal of Cardiovascular Pharmacology 70: 10–15.

Schindelin J, Arganda-Carreras I, Frise E, Kaynig V, Longair M, Pietzsch T, Preibisch S, Rueden C, Saalfeld S, Schmid B, et al. 2012. Fiji: an open-source platform for biological-image analysis. Nature methods 9: 676–82.

Scimone ML, Lapan SW, Reddien PW. 2014. A forkhead Transcription Factor Is Wound-Induced at the Planarian Midline and Required for Anterior Pole Regeneration. PLoS Genetics 10: e1003999.

Shimizu T, Hisamoto N. 2020. Factors regulating axon regeneration via JNK MAP kinase in Caenorhabditis elegans. The Journal of Biochemistry 167: 433–439.

Sievers F, Higgins DG. 2014. Clustal Omega. Current Protocols in Bioinformatics 48.

Smrcka A v. 2008. G protein βγ subunits: Central mediators of G protein-coupled receptor signaling. Cellular and Molecular Life Sciences 65: 2191–2214.

Snow BE, Krumins AM, Brothers GM, Lee S-F, Wall MA, Chung S, Mangion J, Arya S, Gilman AG, Siderovski DP. 1998. A G protein subunit-like domain shared between RGS11 and other RGS proteins specifies binding to G 5 subunits. Proceedings of the National Academy of Sciences 95: 13307–13312.

Swapna LS, Molinaro AM, Lindsay-Mosher N, Pearson BJ, Parkinson J. 2018. Comparative transcriptomic analyses and single-cell RNA sequencing of the freshwater planarian Schmidtea mediterranea identify major cell types and pathway conservation. Genome Biology 19: 124.

Syrovatkina V, Alegre KO, Dey R, Huang X-Y. 2016. Regulation, Signaling, and Physiological Functions of G-Proteins. Journal of Molecular Biology 428: 3850–3868.

Tang W-J, Gilman AG. 1991. Type-Specific Regulation of Adenylyl Cyclase by G Protein βγ Subunits. Science 254: 1500–1503.

Tewari AG, Stern SR, Oderberg IM, Reddien PW. 2018. Cellular and Molecular Responses Unique to Major Injury Are Dispensable for Planarian Regeneration. Cell Reports 25: 2577–2590.e3.

Tu KC, Cheng L-C, TK Vu H, Lange JJ, McKinney SA, Seidel CW, Sánchez Alvarado A. 2015. Egr-5 is a post-mitotic regulator of planarian epidermal differentiation. eLife 4.

Vogg MC, Owlarn S, Pérez Rico YA, Xie J, Suzuki Y, Gentile L, Wu W, Bartscherer K. 2014. Stem cell-dependent formation of a functional anterior regeneration pole in planarians requires Zic and Forkhead transcription factors. Developmental Biology 390: 136–148.

Wagner DE, Wang IE, Reddien PW. 2011. Clonogenic Neoblasts Are Pluripotent Adult Stem Cells That Underlie Planarian Regeneration. Science 332: 811–816.

Wall MA, Coleman DE, Lee E, Iñiguez-Lluhi JA, Posner BA, Gilman AG, Sprang SR. 1995. The structure of the G protein heterotrimer Giα1β1γ2. Cell 83: 1047–1058.

Wenemoser D, Lapan SW, Wilkinson AW, Bell GW, Reddien PW. 2012. A molecular wound response program associated with regeneration initiation in planarians. Genes & Development 26: 988–1002.

Wise A, Gearing K, Rees S. 2002. Target validation of G-protein coupled receptors. Drug Discovery Today 7: 235–246.

Wise A, Jupe SC, Rees S. 2004. The Identification of Ligands at Orphan G-Protein Coupled Receptors. Annual Review of Pharmacology and Toxicology 44: 43–66.

Witchley JN, Mayer M, Wagner DE, Owen JH, Reddien PW. 2013. Muscle Cells Provide Instructions for Planarian Regeneration. Cell Reports 4: 633–641.

Wong GT, Gannon KS, Margolskee RF. 1996. Transduction of bitter and sweet taste by gustducin. Nature 381: 796–800.

Wurtzel O, Cote LE, Poirier A, Satija R, Regev A, Reddien PW. 2015. A Generic and Cell-Type-Specific Wound Response Precedes Regeneration in Planarians. Developmental Cell 35: 632–645.

Yarfitz S, Hurley JB. 1994. Transduction mechanisms of vertebrate and invertebrate photoreceptors. The Journal of biological chemistry 269: 14329–32.

Yu F-X, Zhao B, Guan K-L. 2015. Hippo Pathway in Organ Size Control, Tissue Homeostasis, and Cancer. Cell 163: 811–828.

Zamanian M. 2011. Genomic and functional characterization of G protein-coupled receptors in the human pathogen Schistosoma mansoni and the model planarian Schmidtea mediterranea. Ames.

Zamanian M, Kimber MJ, McVeigh P, Carlson SA, Maule AG, Day TA. 2011. The repertoire of G protein-coupled receptors in the human parasite Schistosoma mansoni and the model organism Schmidtea mediterranea. BMC Genomics 12: 596.

Zeng A, Li H, Guo L, Gao X, McKinney S, Wang Y, Yu Z, Park J, Semerad C, Ross E, et al. 2018. Prospectively Isolated Tetraspanin+ Neoblasts Are Adult Pluripotent Stem Cells Underlying Planaria Regeneration. Cell 173: 1593–1608.e20.

Ziegler K, Kurz CL, Cypowyj S, Couillault C, Pophillat M, Pujol N, Ewbank JJ. 2009. Antifungal Innate Immunity in C. elegans: PKCδ Links G Protein Signaling and a Conserved p38 MAPK Cascade. Cell Host & Microbe 5: 341–352.

Zugasti O, Bose N, Squiban B, Belougne J, Kurz CL, Schroeder FC, Pujol N, Ewbank JJ. 2014. Activation of a G protein–coupled receptor by its endogenous ligand triggers the innate immune response of Caenorhabditis elegans. Nature Immunology 15: 833–838.

