## Supplemental Figures for "Heterotrimeric G proteins regulate planarian regeneration and behavior"

Supplemental_Fig_S1.tif

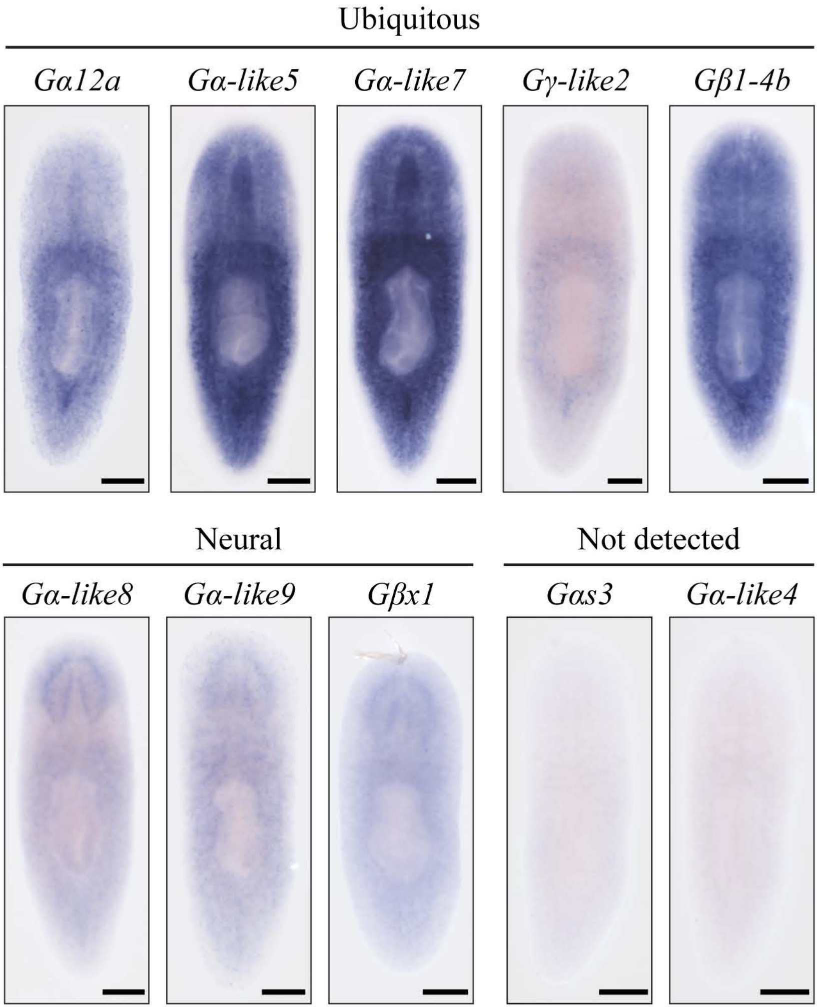


**Supplemental Figure S1. Additional G protein subunit expression patterns.** Representative images of G protein subunit ISH that were not included in Fig. 1 of the main text. Images are grouped by pattern. Scale = 200μm.

Supplemental_Fig_S2.tif

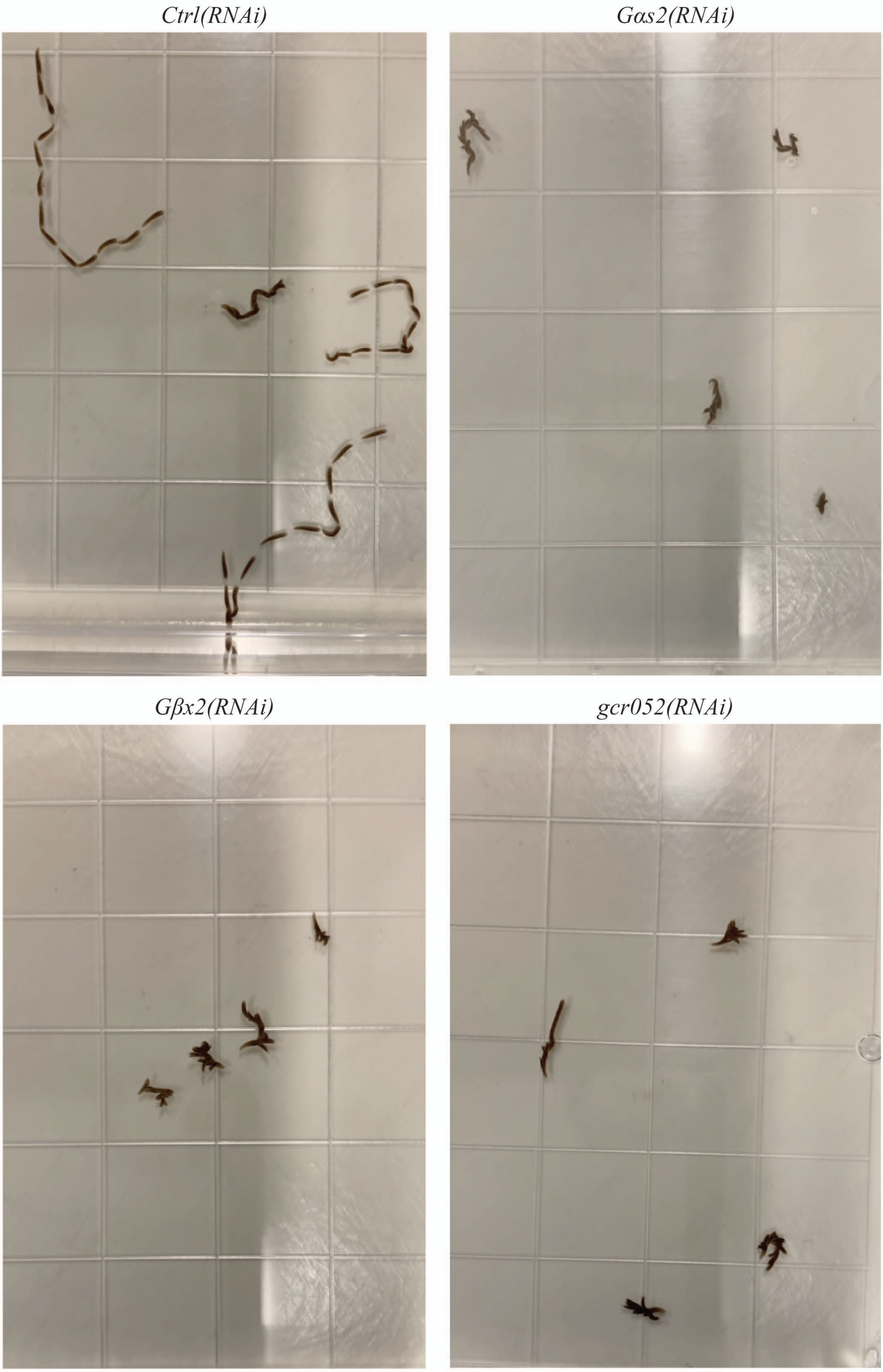


**Supplemental Figure S2. RNAi targeting *Gas2*, *Gβx2*, and *gcr052* results in less forward distance traveled over time.** Overlay images of 5 second increments showing forward distance traveled by control, *Gas2(RNAi)*, *Gβx2(RNAi)*, and *gcr052(RNAi)* animals over the span of 1 minute. Each grid square = 13x13 mm.

Supplemental_Fig_S3.tif

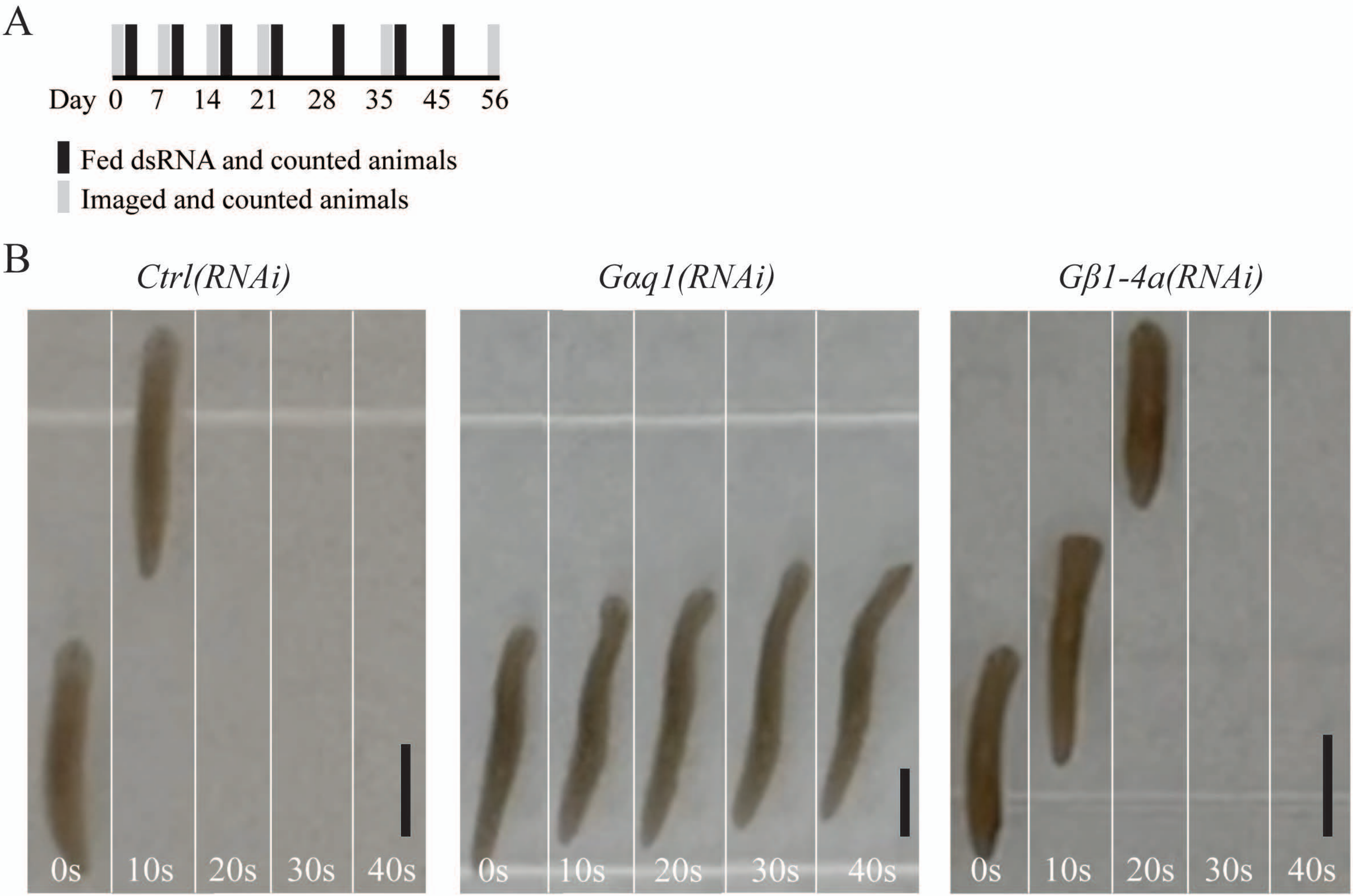


**Supplemental Figure S3. *Gαq1* and *Gβ1-4a* influence animal movement.** (**A**) Long-term RNAi paradigm used on animals in image stills. Behavior videos were taken during the growth/survival assay described in Fig. 4. (**B**) Image stills from videos capturing locomotion displayed by non-amputated control, *Gαq1(RNAi)* and *Gβ1-4a(RNAi)* animals after 28 days of RNAi. Scale = 2 mm.

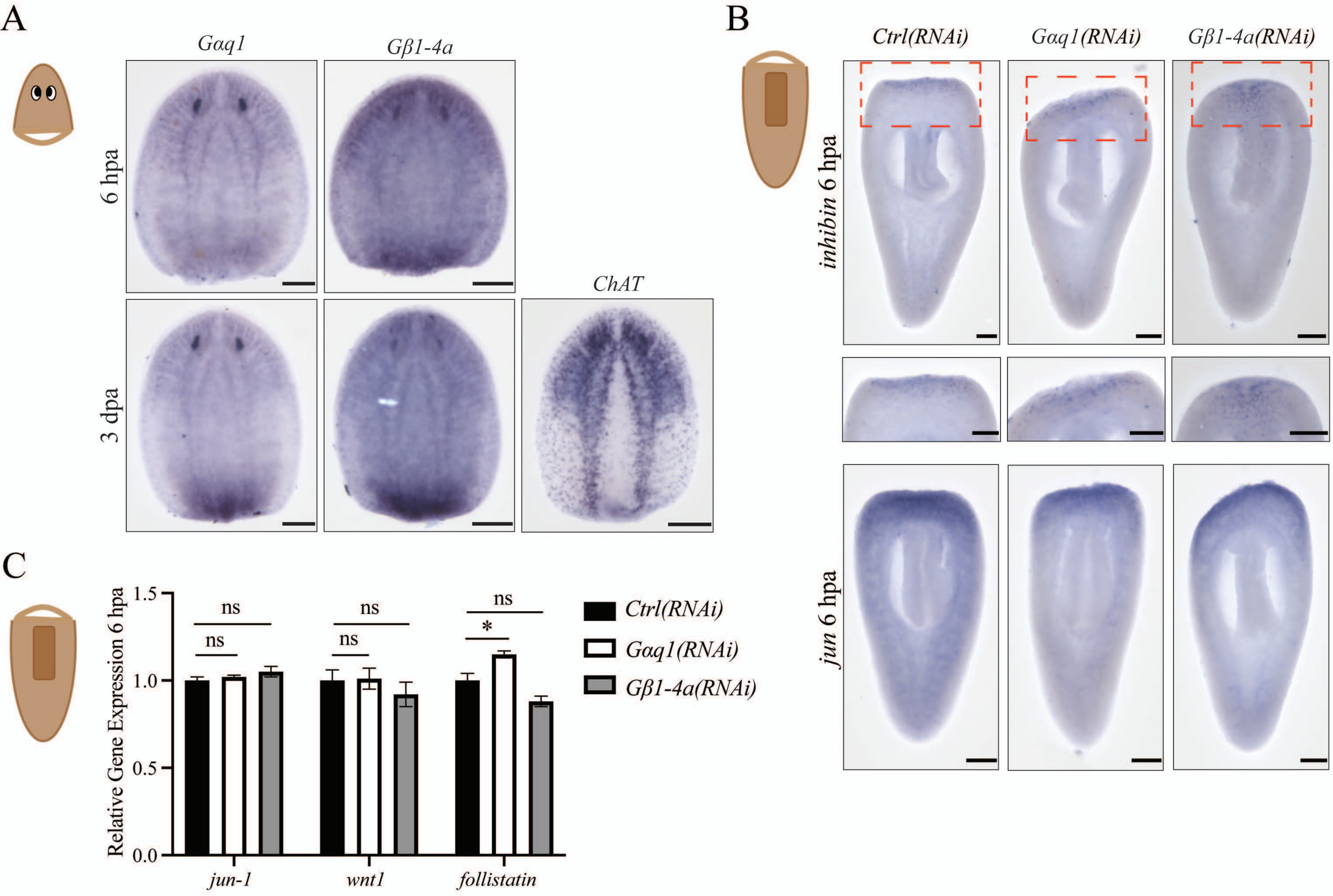


**Supplemental Figure S4.** ***Gαq1*** **and** ***Gβ1-4a*** **are upregulated after injury but are not required for described wound response programs. (A)** Representative images of *Gαq1* and *Gβ1-4a* ISH in in untreated animals 6 hpa and 3 dpa. Example image of *ChAT* ISH at 3 dpa, for reference. **(B)** Representative images of *inhibin* and *jun-1* ISH in *Gαq1(RNAi)* and *Gβ1-4a(RNAi)* animals 6 hpa. Zoomed images are the red-dashed box regions in total body images. **(C)** Relative expression levels of wound response markers 6 hpa, measured by RT-qPCR. Error bars represent SEM. Differences were analyzed with one-way ANOVA (* = P value ≤ 0.05). Visual schematics of amputation scenarios included with each data. Scale = 200μm.

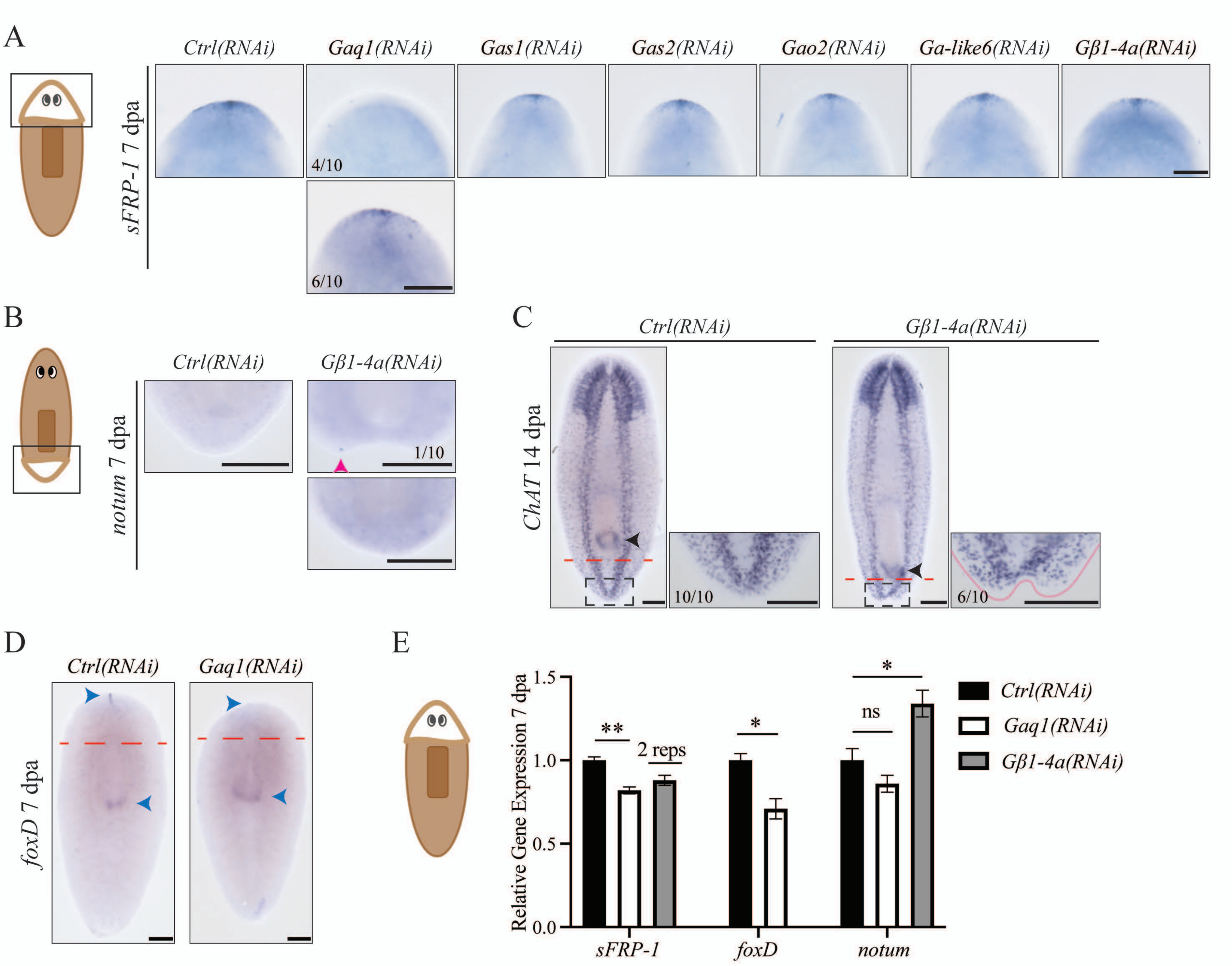


**Supplemental Figure S5. Additional characterization of *Gαq1(RNAi)* and *Gβ1-4a(RNAi)* animals.** (**A**) Visual schematic of image perspective and representative images of *sFRP-1* ISH in animals treated with RNAi targeting *Gαq1*, *Gαs1*, *Gαs2*, *Gαo2*, *Gα-like6*, or *Gβ1-4a* 7 dpa. (**B**) Visual schematic of image perspective and representative images of *notum* ISH in the posterior of *Gβ1-4a(RNAi)* animals 7 dpa. Magenta arrowhead indicates detected expression in the regenerating tail region in 1/10 animals. (**C**) Representative images of *ChAT* ISH in *Gβ1-4a(RNAi)* animals 14 dpa. Red dashed lines indicate the amputation site and black arrowheads indicate expression in the pharynx. (**D**) Representative whole-body images (zoomed inserts at anterior pole in Fig. 6E) of *foxD* ISH in *Gαq1(RNAi)* animals 7 dpa. Red dashed lines indicate the amputation site and blue arrowheads indicate the two main domains of *foxD* expression (at the anterior pole and pharynx (Roberts-Galbraith and Newmark 2013; Scimone et al. 2014)). (**E**) Visual schematic of amputation scenario and relative expression levels of anterior pole markers 7 dpa, measured by RT-qPCR. Data for *sFRP-1* and *foxD* analyzed with Unpaired t test, and data for notum analyzed with One-way ANOVA. Error bars represent SEM. * = P value ≤ 0.05. ** = P value ≤ 0.005. Scale = 200µm.

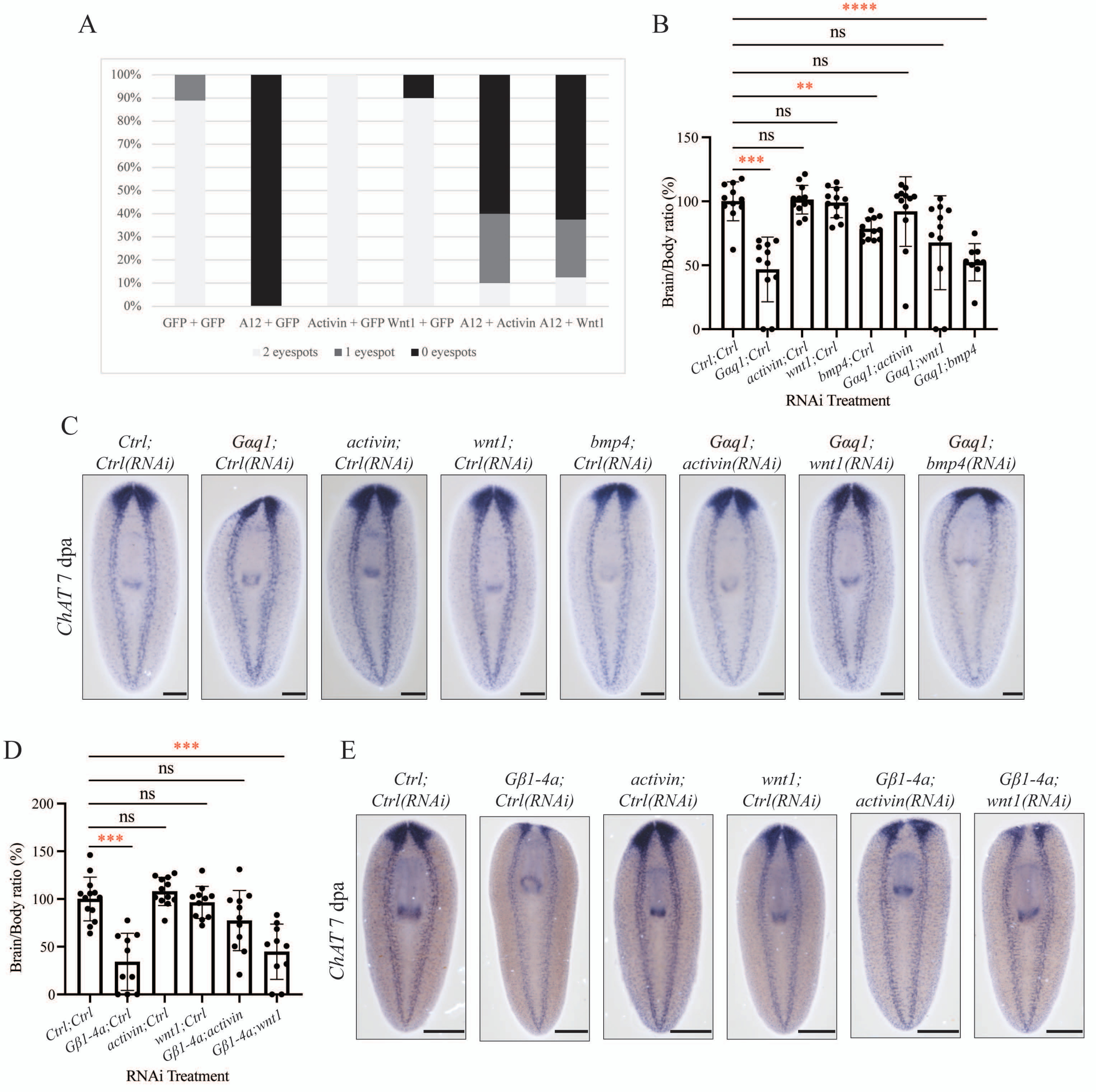


**Supplemental Figure S6. *Gαq1* and *Gβ1-4a* regeneration phenotypes are rescued to varying degrees by co-targeting of posterior-promoting pathway signals.** (**A**) Table representing proportions of animals with one, two, or no eyespots in the RNAi animals from the rescue experiments in Fig. 6, 7 dpa. (**B**) Bar graph showing results from quantification of brain/body ratios in second repetition of rescue experiments, including *bmp4* control. Differences in sample means analyzed with Brown-Forsythe and Welch ANOVA. Error bars represent standard deviation. ** = P value ≤ 0.006. *** = P value = 0.0005. **** = P value ≤ 0.0001. Experiment used the same RNAi paradigm and brain quantification method as described in Fig. 2. (**C**) Representative images showing *ChAT* expression 7 dpa from rescue experiments in (B). (**D**) Bar graph showing results from quantification of brain/body ratios from rescue experiments with *Gβ1-4a(RNAi)* animals. Differences in sample means were statistically analyzed with Brown-Forsythe and Welch ANOVA. Error bars represent SEM. *** = P value ≤ 0.0006. (**E**) Representative images showing *ChAT* expression 7 dpa from rescue experiments in (D). Scale = 200μm (C) and 500 μm (E).

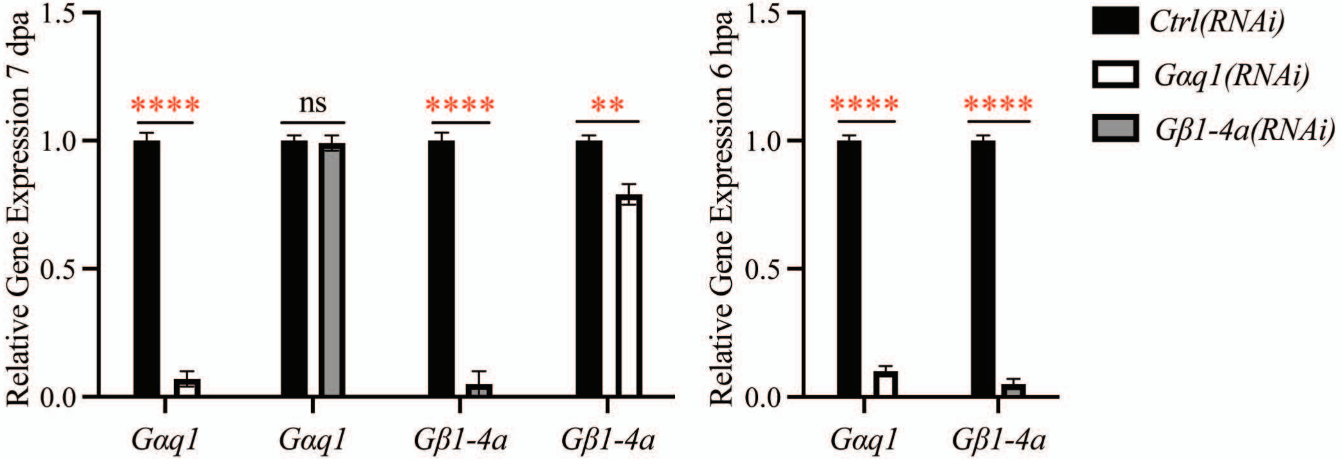


**Supplemental Figure S7. *Gαq1* and *Gβ1-4a* expression in *Gαq1(RNAi)* and *Gβ1-4a(RNAi)* animals.** Relative expression levels of *Gαq1* and *Gβ1-4a* in *Gαq1(RNAi)* and *Gβ1-4a(RNAi)* animals, measured by RT-qPCR. Differences in sample means were statistically analyzed with Unpaired t test. Error bars represent SEM. ** = P value ≤ 0.01 and **** = P value ≤ 0.0001. All RT-qPCR data throughout this work utilize the cDNA used to produce the data displayed here.

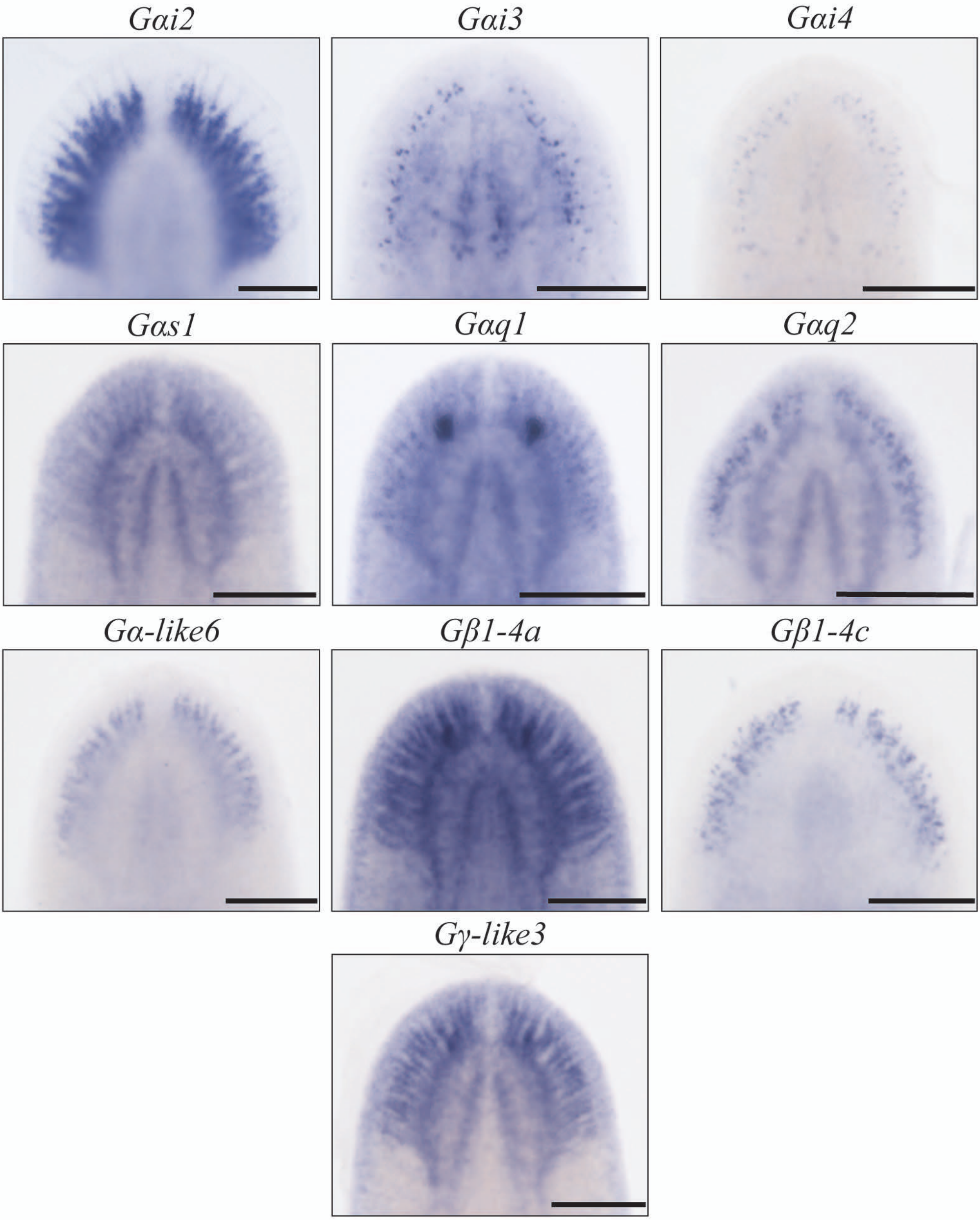


**Supplemental Figure S8. Many heterotrimeric G protein-encoding genes in *S. mediterranea* display enriched expression in the brain branches.** Images zoomed at the head region of ISH for G protein subunits that showed brain branch expression. Staining patterns displayed include those that visualize the branches extending toward the periphery of the animal and individual cell clusters showing regionalized localization in the branches. Scale = 200μm.
