## Supplementary material for "Heterotrimeric G proteins regulate planarian regeneration and behavior": Supp. File 1

```
                                                                   α1                β1
RatGnai1[Ai]                                              ..............................
                      1                            10     ▲  ▲▲   20▲▲   ▲▲    ▲▲ 30
RatGnai1[Ai]     ........MGCTL...............SAEDKA.AVERSKMIDRNLREDGEKAAREVKL
22522            ........MNCNFKISSHLN......TINQISTQKSVLRSREIDQQLMQDQELRKKEIQI
2931             .......MLPCIPK.RSRF........IGDVDIKHQKAKSREIDKLMKREKRKFLTRHSI
8947             .......MFTCFKNSKKKF........YKEIDLKSQKSFSKLIDKSLKIERRKNLMKQHI
33116            ............................................................
9665             ......MPAICGGKSA.................STEDKSNKEIDKKLKKDKKTFLMTHRI
6167             ......MALSCAAF............KTETEEQKNRKESNKRIEKCIEKEKRSFKSTHRL
5082             ........MACCN...............SEEFKE.QSRINKEIEKQLKRDKKDARRELKL
3755             ........MGCCL...............SELDKE.QKRINKEIEKQLKKDKKESRRELKL
1773             ........MGCPL...............GKPDEIEQNEISHKIDKELKLEAKHSYRIVKL
13658            ............................................................
9715             ........MGCAM...............SLDERI.ALNKSKDIDKKLKEDGLQLQRDIKL
1656             ........MGCTM...............SKDERD.ALDKSRNIDKKLKEDGMQAAKDVKL
12112            ........MGCAG...............SVPDKD.QQEKNKEIEQQLRQDAEKAMKEVKL
19937            ........MGCIN...............STKNVI.QEEAHKEIEKQLRIDQEKAMREIKL
47050complete    ........MGCIN...............SSHDEP.QQELNKNIEKQLRIDAEKAMKEVKL
4493             .......MNTCISKVETDEQREKR.....VLLEENRRKNNQIEKELALYKNKKLSCLKL
HumanGNAS[As]    ........MGCLGNSK.........TEDQRNEEKAQREANKKIEKQLQKDKQVYRATHRL
HumanGNA12[A12]  MSGVVRTLSRCLLPAEAGGARERRAGSGARDAEREARRRSRDIDALLARERRAVRRLVKI
HumanGNAQ[Aq]    ..MTLESIMACCL...............SEEAKE.ARRINDEIERQLRRDKRDARRELKL
HumanGNAI1[Ai]   ........MGCTL...............SAEDKA.AVERSKMIDRNLREDGEKAAREVKL
HumanGNAO1[Ao]   ........MGCTL...............SAEERA.ALERSKAIEKNLKEDGISAAKDVKL

                             α2                α3              α4
RatGnai1[Ai]     →      TT
                    40        50          60        70         80        90
RatGnai1[Ai]     LLLGAGESGKSTIVKQMKIIH..EAGYSEEE.CKQYKAVVYSNTIQSIIAIIRAMGRLKI
22522            LVLGNRNSGKTTFLKQFRIHH..GDGYPYLQ.RAILVPDILSNLADAIHIVMDNMARWDL
2931             LLLGTGESGKSTFLKQMKLIN..GKKFTSAE.LAGFKDTIYDNVYKGILFLLHARSSLNI
8947             LLLGTGESGKSTFLKQMKIIC..GQKFTVYE.IESFKDIIYDNIFKGVLFLIDARAQLNI
33116            ................MKLLSSINQTFPDVY.RAKFIPEIQRNLVQALTAILQFMDQIKL
9665             PVVGAGESGKSTLIKQMQILY..INGFNDEE.RKSKIKDIRGNIKDGMLAVTGAMKTLKP
6167             LLLGAGESGKSTIVKQMKILH..IDGFSNEE.KKQKAEDIRKNLRDAILTITSAMSNINP
5082             LLLGTGESGKSTFIKQMRIIH..GTGYSDDD.KRSFIKLIYQNIYLATYTLIRAMEVLKI
3755             LLLGTGESGKSTFIKQMRIIH..GSGYSDED.KRSFIKLVYQNIYLAIYTLILAMENLAI
1773             LLLGTGESGKSTIVKQMKLIV..NNGHSQAE.RLGFRSLIFSNTIQSLIVIIRAMSKLDI
13658            ...............................................MIDILYAVHHLNI
9715             LLLGAGESGKSTIMKQMKIIH..EGGFSSTD.NHHYKPIVYSNTIQSVAAILRAMHKLNI
1656             LLLGAGESGKSTIVKQMKIIH..EGGFTQED.NKQYKPVVYSNTTQSMIAILRAMTTLGI
12112            LLLGAGECGKSTILKQMTIIH..GKGYPEEE.RKEYVPIIYANVVQSMVTILNAMETLGI
19937            LLLGAGECGKSTILKQMSIIH..GGGYSDEE.KKEFIPIIYANIVQSMGAILKAMDILEI
47050complete    LLLGAGECGKSTILKQMSIIH..GDGYSESE.RKKFIPIIYSNIIQSVVTILKAMETLEI
4493             LLLGTGESGKSTILKQMKIIH..VNGFSKTE.KIEFISNIKRNVRDAIMAIVSMNKLQI
HumanGNAS[As]    LLLGAGESGKSTIVKQMRILH..VNGFNGDSEKATKVQDIKNNLKEAIETIVAAMSNLVP
HumanGNA12[A12]  LLLGAGESGKSTFLKQMRIIH..GREFDQKA.LLEFRDTIFDNILKGSRVLVDARDKLGI
HumanGNAQ[Aq]    LLLGTGESGKSTFIKQMRIIH..GSGYSDED.KRGFTKLVYQNIFTAMQAMIRAMDTLKI
HumanGNAI1[Ai]   LLLGAGESGKSTIVKQMKIIH..EAGYSEEE.CKQYKAVVYSNTIQSIIAIIRAMGRLKI
HumanGNAO1[Ao]   LLLGAGESGKSTIVKQMKIIH..EDGFSGED.VKQYKPVVYSNTIQSLAAIVRAMDTLGI

                              α5                                   α6
RatGnai1[Ai]
                    100                 110                   120
RatGnai1[Ai]     DFGDAARADDAR.............QLFVLAGA....AEEGF.........MTAELAGVI
22522            RFEDPYVQTLAKDFHNNCPRSPALSNNPHNLFKSIQNNSPRPFNRFCKTVNRITDFCLHI
2931             NFENYEETSAEAI....R.IEDHFKRNKEIHKEKA..TTGQI....IWSEEEFLKLVDDF
8947             DWNDNETAN.SAE....E.IERYFDETKLIHRQRR..QSKQL....LWKENEFLEIVQRL
33116            DFKHPNAHLHTAKKELYE.IRDQIDKNPDTIEKYA..NSTEM....D....VRNNFYDNC
9665             PVDLEHAEN..Q.............TLLDFIQHNA..VKPDF....T....YTKEFFDYC
6167             PTKLGNPEN..Q.............KFLDYMQHTA..SKPDF....R....YPSEFYEYC
5082             PYENPDNLEYAK.............DLRDIDY.....ETVTT.........FEPSHIVAT
3755             SYSNPENLSYID.............EIKNIDY.....ESVST.........LEPDHVIAI
1773             SFTNPERLNDAQ.............TLLQLAGT.V..ESNGP.........LNEELYSAM
13658            HFESSAREADEI.............LVNNTITT.G..MDEKP.........LNAQLCNAL
9715             SFENSAREADEI.............LVNNIMTT.M..KDQEP.........FTPQLNSAL
1656             SFGDIDRGADAK.............VVSDVIQA.M..EDTEP.........FSEELLEAM
12112            SMENEAMIEESK.............IVQKYMRN....AEEGE.........FPKELGSAL
19937            PLENANLGTEKN.............TVHESMKN....AEEGG.........MTINTIKAL
47050complete    KLNE.HLHSDKL.............LVIDQVKD....VDNGV.........LRQDAVESI
4493             TPKKFDLSEKIK................FLQT.A..FSEQY....T....YPNTFFDVV
HumanGNAS[As]    PVELANPEN..Q.............FRVDYILSVM..NVPDF....D....FPPEFYEHA
HumanGNA12[A12]  PWQYSENEKHGM.............FLMAFENKAG..LPVEP....A....TFQLYVPAL
HumanGNAQ[Aq]    PYKYEHNKAHAQ.............LVREVDV.....EKVSA.........FENPYVDAI
HumanGNAI1[Ai]   DFGDSARADDAR.............QLFVLAGA....AEEGF.........MTAELAGVI
HumanGNAO1[Ao]   EYGDKERKADAK.............MVCDVVSR.M..EDTEP.........FSAELLSAM
```

```
                    α7                         α8       α9          α10
RatGnai1[Ai]
                    130       140          150       160       170       180
RatGnai1[Ai]        KRLWKDSGVQACFNRSREYQ...LNDSAAYYLNDLDRIAQPNYIPTQQDVLRTRVKTTGI
22522               KNISDQDDFQECLDC.KEFFKPSITYADLYFIQNIDRLIQADYTPTLQDIMVMRKPTYSV
2931                KLIWNDKGIKEAFNQRSKLMTESFSENSRFYLNKLDKIRVKDYQPSNEDIVWSRKPTDSI
8947                KAIWNDKSIQQTFLRRSEIITESFSENTRYYLNKIDEIGTLNYFPTDEDIVWTRKPTESI
33116               KALWRDPNILDTFQRSNEYQ...LIDSAQYFLDKIDLIRQPDYKPSDDDVLQCRTKTLGI
9665                YTLWIDAGVQATFARANEYQ...LIDSTKYFLDQVQTIAKDDYLPTDQDLLRCRVLTSGI
6167                AVLWADKGVLETYERSNEYQ...LIDCAKYFLDQALILGQQNYTPTEQDILRCRVLTSGI
5082                RSLWNDSGVKECYDRRREYQ...LTDSAKYYLDDLDRIVTPDYLPTLQDILRVRVPTTGI
3755                KSLWADVGVKECYDRRREFQ...LTDSAKYYLDSLDRIATPNFLPTLQDILRVRVPTTGI
1773                LNLWQDLGIQECFQRSAEYQ...LIDSAEYYLNSLERLSNSNYVPTEQDILRTRVKTAGI
13658               KSLWSDVGVQECFRTFNEYD...LNYSTQYFLDDIDRISAVDYEPTVQDILRSKMTNYGV
9715                KSLWSDIGVQECFRRSNEYQ...LNDSAQYFLDNVDRISAVDYKPSDQDILRTRIKTTGI
1656                KRLWADPGVQECFGRSNEYQ...LNDSAKYFLDDLDRLGAKDYMPTEQDILRTRVKTTGI
12112               ALLWKDDSVQNCYARAKEYQ...LNDSAGYYLDSLDRLSEPSYIPTEQDVLRSRVKTTGI
19937               TVLWKDTGVQEAFNRSKEYQ...LNDSAEYYLIDLERIASDNYIPTVQDILRSRIKTTGI
47050complete       KKLWQNVEVQQCVQRAKEYN...LNDSAEYYLNDIDRLRGDNYLPNDQDILRSRVKTTGI
4493                SELWQDEDVQASFLRSNEYQ...LIDSAKYFLDHIHIIRQNDYIPSLQDILRCRKMTSGI
HumanGNAS[As]       KALWEDEGVRACYERSNEYQ...LIDCAQYFLDKIDVIKQADYVPSDQDLLRCRVLTSGI
HumanGNA12[A12]     SALWRDSGIREAFSRRSEFQ...LGESVKYFLDNLDRIGQLNYFPSKQDILLARKATKGI
HumanGNAQ[Aq]       KSLWNDPGIQECYDRRREYQ...LSDSTKYYLNDLDRVADPAYLPTQQDVLRVRVPTTGI
HumanGNAI1[Ai]      KRLWKDSGVQACFNRSREYQ...LNDSAAYYLNDLDRIAQPNYIPTQQDVLRTRVKTTGI
HumanGNAO1[Ao]      MRLWGDSGIQECFNRSREYQ...LNDSAKYYLDSLDRIGAADYQPTEQDILRTRVKTTGI
```

```
                    β2           β3               α11  η1        β4      η2
RatGnai1[Ai]        ------->T.....T------->                      ------->
                      190             200            210       220       230
RatGnai1[Ai]        VETHFTFK.....DLHFKMFDVGGQRSERKKWIHCF.EGVTAIIFCVALSDYDLVLAE..
22522               KDFIFNLG.....GITLRFIDLGGYKCQRKKFIHYF.EGVSTVFYFCALSEYSEIIEE..
2931                IEEEIDIH....GA.CIVFIDVGGQTKERKKWCQVF.QDMNSVLFLIASSHFDEFYYDKM
8947                IETEIEVS....NDIILVFIDVGGQTKERRKWCQLF.HDMKSVLFLIACSHFDEEYMDRF
33116               HTEIINYN.....NVNFELVDVGGQRDQRAKWIEALSDGVTAVIFLTDVSAYDMVLAE..
9665                FETKFTMN.....KVCFHMFDVGGQREERRKWIQVF.SDVTAVIFVCACSGYDMTLRE..
6167                FETKFTVD.....KVNFHMFDVGGQRDERRKWIQCF.NDVTAIIFVTACSSYNMVLRE..
5082                IEYPFDLD.....SIIFRMVDVGGQRSERRKWIHCF.EGVMTMIYVASISEYDQFLIE..
3755                IEYPFDLD.....SIIFRMVDVGGQRSERRKWIHCF.ENVTSIMFLVALSEYDQGLVE..
1773                IETTVEFK.....ELIMKIVDVGGQRAERKKWIHCF.DNVDAIIFIVALNEYDLRLRE..
13658               CECQIKYK.....ERMFTIFDTSVRLS.DRKWRICF.EDPSALIFCVSLTEYLK......
9715                VEIQFKFK.....NMNFKIFDVGGQRSERKKWIHCF.EDVTAIIFCVSLSEYDQVLVE..
1656                VEVHFMFK.....NMNFKLFDVGGQRSERKKWIHCF.EDVTAIIFCVAMSEYDQVLHE..
12112               VETHFHFK.....DLDFKVFDVGGQRSERKKWMHCF.EGVTAILFLVAMSEYDLKLVE..
19937               IETYFHFK.....DLDFKVFDVGGQRSERKKWMHCF.EGVTAILFLVALSEYDLRLVE..
47050complete       IETEFHFK.....NLDFKVYDVGGQRSERKKWMHCF.EGVTAILFLIAMSEYDLKLVE..
4493                TEISFEVREKKKNKVNFKVFDVGGQQGERKRWIQLF.GEVTAILFLADCSSFDQTLRE..
HumanGNAS[As]       FETKFQVD.....KVNFHMFDVGGQRDERRKWIQCF.NDVTAIIFVVASSSYNMVIRE..
HumanGNA12[A12]     VEHDFVIK.....KIPFKMVDVGGQRSQRQKWFQCF.DGITSILFMVSSSEYDQVLME..
HumanGNAQ[Aq]       IEYPFDLQ.....SVIFRMVDVGGQRSERRKWIHCF.ENVTSIMFLVALSEYDQVLVE..
HumanGNAI1[Ai]      VETHFTFK.....DLHFKMFDVGGQRSERKKWIHCF.EGVTAIIFCVALSDYDLVLAE..
HumanGNAO1[Ao]      VETHFTFK.....NLHFRLFDVGGQRSERKKWIHCF.EDVTAIIFCVALSGYDQVLHE..
```

```
                         α12            η3       β5          α13          η4
RatGnai1[Ai]                                   ------->
                              250       260       270       280       290
RatGnai1[Ai]        .DEEMNRMHESMKLFDSICNNKWFTDTSIILFLNKKDLFEEKIK..KSPLTICYPEYAGS
22522               ..TTKNKLNNSLENFEEIINNKYLWRKDFVIFLNKSDLFKSKVE..NLSINVFFSDFEGN
2931                SKEYRNKLREAMRVFEDLINVQYFLSVSVIIFFNKTDVLTEKVTNKISDIRSEFTEYPEN
8947                TLQKRNKLKEAMFVFEELINQNAFLRVSVLLFFNKTDILSEKIKSHQSNIGKDFQDFPGE
33116               .DQTTNRLRESVSLLGQVWTKNPLRDKSIILFLNKQDKLEMKVRKGRTQIETYFPEFKQE
9665                .AQKQNRLRECISLFSEVWGNRYLRQTSIILFLNKQDLFVQKLTSGKTKIVDFFPEFAEY
6167                .DASQNRLRESLELLKSIWNNRWLRNISVILFLNKQDVLKEKVLAGKSKIEDYFPDYLRY
5082                .DNEINRMFESIKLFDSVCNNQWFSKCSIILFLNKTDLFKLKIV..KSPLTVCFPEYKGN
3755                .SDNDNRMEESKALFRTIITYPWFHNASVILFLNKKDLLEEKII..YSHLVDYFPEYEGP
1773                .DPEVNRMMESLRLFDSMCNNVVFKDTCMILFLNKRDLFEVKIK..KSPLSICFEEYADE
13658               ....EDEMACSMRYFNSIIQFKWFQNSSFILFLNKKDLFKRKLL..DHPITCCFPEYTGL
9715                .DDATNRMQESLFLFESICNNVWFLHTSFILFLNKKDLFLEKLQ..ICPITFCFPEYKGP
1656                .DETTNRMQESLKLFDSICNNKWFTQTSIILFLNKKDLFAEKIK..RSPLTVCFSEYTGR
12112               .DSSTNRMHESMRLFDSICNSQWFVNTSIILFLNKKDLFGEKVV..KSPLTVCFPEYTGA
19937               .DSTTNRMHESMKLFDSICNSPWFVNTSVILFLNKVDLFEIKIQ..RSPLTICFPEYLGP
47050complete       .DQTTNRMHESMKLFESICNSQWFTSTSIILFLNKKDLFMKKIE..ISPITICFKDYNGP
4493                .DRSKNRLIDSLEVFYQAWMNRYLQNVPIIVFVNKIDMLELKIQNEHS.IESMINEITNL
HumanGNAS[As]       .DNQTNRLQEALNLFKSIWNNRWLRTISVILFLNKQDLLAEKVLAGKSKIEDYFPEFARY
HumanGNA12[A12]     .DRRTNRLVESMNIFETIVNNKLFFNVSIILFLNKMDLLVEKVK..TVSIKKHFPDFRGD
HumanGNAQ[Aq]       .SDNENRMEESKALFRTIITYPWFQNSSVILFLNKKDLLEEKIM..YSHLVDYFPEYDGP
HumanGNAI1[Ai]      .DEEMNRMHESMKLFDSICNNKWFTDTSIILFLNKKDLFEEKIK..KSPLTICYPEYAGS
HumanGNAO1[Ao]      .DETTNRMHESLMLFDSICNNKFFIDTSIILFLNKKDLFGEKIK..KSPLTICFPEYTGP
```

```
RatGnai1[Ai]

RatGnai1[Ai]        ............................................................
22522               PS..........................................................
2931                FDPHNIV.....................................................
8947                SDPYNLV.....................................................
33116               KCKHIFELIKDLKAARGKRSKKESDIWEKYFSYFLPQMRDQKKTENPEDATRQQRET...
9665                ............................................................
6167                ............................................................
5082                ............................................................
3755                ............................................................
1773                ............................................................
13658               ............................................................
9715                ............................................................
1656                ............................................................
12112               ............................................................
19937               ............................................................
47050complete       ............................................................
4493                TPHSTKNSAKQLNNRRSSLNINHKTNN........DNMTSCSKCENSHPITDNNKCIHLC
HumanGNAS[As]       ............................................................
HumanGNA12[A12]     ............................................................
HumanGNAQ[Aq]       ............................................................
HumanGNAI1[Ai]      ............................................................
HumanGNAO1[Ao]      ............................................................


RatGnai1[Ai]                                        ℓ...........................

RatGnai1[Ai]        ..............................NTY...........................
22522               ............................................................
2931                ..............................Q...........................VQ
8947                ..............................D...........................VQ
33116               .......QIIN..............EYS..QIQSQVAKVFNDGSVDKWLTDGDDDDIFSH
9665                ..............................QSK............NTFDS..........
6167                ..............................TLP............ADVQC..........
5082                ..............................NNF...........................
3755                ..............................QRD...........................
1773                ..............................NTY...........................
13658               ..............................QEY...........................
9715                ..............................QEY...........................
1656                ..............................QTY...........................
12112               ..............................NIY...........................
19937               ..............................NTF...........................
47050complete       ..............................HNY...........................
4493                PESNSNSKICDRKISIRTIPSCSSRRFSKDNNFEFVAQTFSENPWGKYQPSNEECKEFVN
HumanGNAS[As]       ..............................TTP............EDATP..........
HumanGNA12[A12]     ..............................PHR...........................
HumanGNAQ[Aq]       ..............................QRD...........................
HumanGNAI1[Ai]      ..............................NTY...........................
HumanGNAO1[Ao]      ..............................NTY...........................


                            α14
RatGnai1[Ai]        ...............................ℓℓℓℓℓℓℓℓℓℓℓℓℓ
                                                     300         310

RatGnai1[Ai]        ................................EEAAAYIQCQFEDL..N.......KRKD
22522               ..............................DVEDGINFFRMKFLSLKPA.......NAPK
2931                QFLVDSFVSFVDNPGA...............HP.RSQYSAGNQNQGR.ASIAG.QAGVPK
8947                MFIVDSFVKLIDNPYG...............NDKRASYIRSNSVSGN.KRVVSPVPMQIP
33116               LMKNDEFISFH..........KRIMNITFYQVVSITFFIENQFLHQC.........EQTS
9665                ..........Q..............SDEPSEVTYAKNFVKDKFMST.TK.......KEEG
6167                ..........H.............NHIEEAEFTRAKYFFRDEFLKITTT.......GNDG
5082                ................................EDASEYVRMTFEML..N.......K.KK
3755                ...............................AEAARDFILKMFIEL..N.......P.DQ
1773                ................................EKAVTYIKFKFESL..N.......KYRQ
13658               ................................DPSIEYIEKRFRNE..V.......K.DS
9715                ................................DSSVSYIEQCFRSK..N.......K.DV
1656                ................................EEAAAYIQASFEAK..N.......N.SP
12112               ................................EEASAYIQSQFEEL..N.......KKKE
19937               ................................VDTSNYIREKFEGL..N.......KKKN
47050complete       ................................EETSNFIRLKFEDL..N.......KSKD
4493                SFPVDQ.VDLHSSGTHAAKKKSKDVHNIHPDTIKTACYIKNIFAQITRNHPKNCDMKISN
HumanGNAS[As]       ..........E..............PGEDPRVTRAKYFIRDEFLRISTA.......SGDG
HumanGNA12[A12]     ...............................LEDVQRYLVQCFDRK..R.......R.NR
HumanGNAQ[Aq]       ...............................AQAAREFILKMFVDL..N.......P.DS
HumanGNAI1[Ai]      ................................EEAAAYIQCQFEDL..N.......KRKD
HumanGNAO1[Ao]      ................................EDAAAYIQAQFESK..N.......R.SP
```

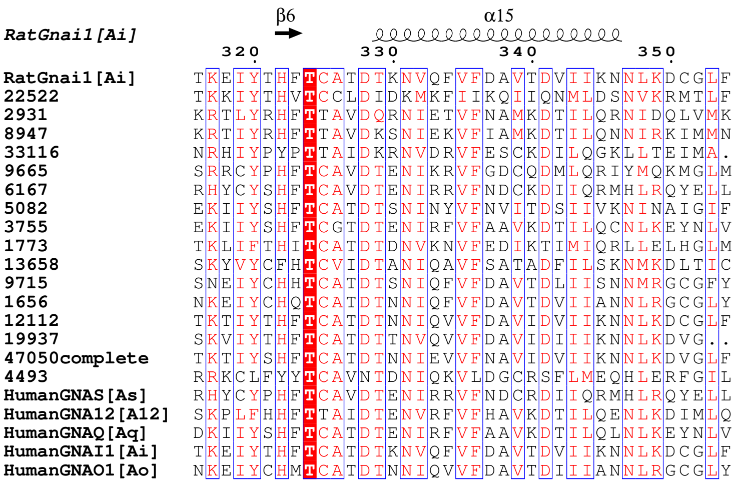

RatGnai1[Ai]
β6
α15
320
330
340
350
RatGnai1[Ai] TKEIYTHFTCATDTKNVQFVFDAVTDVIIKNNLKDCGLF
22522 TKKIYTHVTCCLDIDKMKFIIKQIIQNMLDSNVKRMTLF
2931 KRTLYRHFTTAVDQRNIETVFNAMKDTILQRNIDQLVMK
8947 KRTIYRHFTTAVDKSNIEKVFIAMKDTILQNNIRKIMMN
33116 NRHIYPYPTTAIDKRNVDRVFESCKDILQGKLLTEIMA.
9665 SRRCYPHFTCAVDTENIKRVFGDCQDMLQRIYMQKMGLM
6167 RHYCYSHFTCAVDTENIRRVFNDCKDIIQRMHLRQYELL
5082 EKIIYSHFTCATDTSNINYVFNVITDSIIVKNINAIGIF
3755 EKIIYSHFTCGTDTENIRFVFAAVKDTILQCNLKEYNLV
1773 TKLIFTHITCATDTDNVKNVFEDIKTIMIQRLLELHGLM
13658 SKYVYCFHTCVIDTANIQAVFSATADFILSKNMKDLTIC
9715 SNEIYCHHTCATDTSNIQFVFDAVTDLIISNNMRGCGFY
1656 NKEIYCHQTCATDTNNIQFVFDAVTDVIIANNLRGCGLY
12112 TKTIYTHFTCATDTNNIQVVFDAVIDVIIKNNLKDCGLF
19937 SKVIYTHFTCATDTTNVQVVFDAVIDIIIKNNLKDVG..
47050complete TKTIYSHFTCATDTNNIEVVFNAVIDVIIKNNLKDVGLF
4493 RRKCLFYYTCAVNTDNIQKVLDGCRSFLMEQHLERFGIL
HumanGNAS[As] RHYCYPHFTCAVDTENIRRVFNDCRDIIQRMHLRQYELL
HumanGNA12[A12] SKPLFHHFTTAIDTENVRFVFHAVKDTILQENLKDIMLQ
HumanGNAQ[Aq] DKIIYSHFTCATDTENIRFVFAAVKDTILQLNLKEYNLV
HumanGNAI1[Ai] TKEIYTHFTCATDTKNVQFVFDAVTDVIIKNNLKDCGLF
HumanGNAO1[Ao] NKEIYCHMTCATDTNNIQVVFDAVTDIIIANNLRGCGLY

```
Ratbeta1
Ratbeta1          ..................................................MSELDQLRQE
2872              ....................................METSSNHVNAEEEISVEEKKLTLE
3309              ................................MDKTIENPCADDNGDGEMCAEEHKLLDE
DrosophilabetaE   ..............................................MPKIDPETQKLYDE
31894             ................................................MMSEDRDAIRQE
7509              ..................................................MTEYDRLTDE
Humanbeta3        ..................................................MGEMEQLRQE
Mousebeta3        ..................................................MGEMEQLRQE
Mousebeta2        ..................................................MSELEQLRQE
Humanbeta2        ..................................................MSELEQLRQE
Humanbeta1        ..................................................MSELDQLRQE
Mousebeta1        ..................................................MSELDQLRQE
Humanbeta4        ..................................................MSELEQLRQE
Mousebeta4        ..................................................MSELEQLRQE
Drosophilabeta1   ..................................................MNELDSLRQE
Celegansbeta1     ..................................................MSELDQLRQE
1184              ..................................................MTELDQLRQE
1615              ..................................................MSELDQIRQE
10481             ...................................MS.........VEAPINNDGDLSKE
Celegansbeta5     ......................MSTIAGESSSSSKMPENSQP..TTTEKGSEYLEQLANE
Drosophilabeta5   ...................................MSEAAVPASNANANATEKMASLVRE
Humanbeta5        MCDQTFLVNVFGSCDKCFKQRALRPVFKKSQQLSYCSTCAEIMATEGLHENETLASLKSE
Mousebeta5        MCDQTFLVNVFGSCDKCFKQRALRPVFKKSQQLNYCSTCAEIMATDGLHENETLASLKSE
```

## WD 1

```
                  α1                                  β1         β2
Ratbeta1                                                                 TT
                           20        30        40        50        60        70
Ratbeta1          AEQLKNQIRDARKACADATLSQITNNIDPVGRIQMRTRRTLRGHLAKIYAMHWGTDSRLL
2872              IEKIKEQLDIARQAMQDTDYFKMTEHVKELGRIKFKVRKQLHGHLSKITGICWAQDSQLL
3309              IESVKERLDSVRQEVQDTDFFQMTEHVKEVGRLKFKTCRQLRAHLSKVTSLDWASDSQLL
DrosophilabetaE   INGMIQKFKDDQKSKADCTLADKCGDMGDVPKIRFSSKKILKGHINKVNSVHFAGDSRHC
31894             IEDLKARIREQRAAVSDSTLEEITKGQLGINKLLIRLRYTLRGHLSKIYALHWAGDSRNV
7509              IRGLRDKIEKTRKKCADTTLPNVTKSIQSLPRIQPKTRRTLSGHLAKIYAMQWCKDSRNL
Humanbeta3        AEQLKKQIADARKACADVTLAELVSGLEVVGRVQMRTRRTLRGHLAKIYAMHWATDSKLL
Mousebeta3        AEQLKKQIADARKACADITLAELVSGLEVVGRVQMRTRRTLRGHLAKIYAMHWATDSKLL
Mousebeta2        AEQLRNQIRDARKACGDSTLTQITAGLDPVGRIQMRTRRTLRGHLAKIYAMHWGTDSRLL
Humanbeta2        AEQLRNQIRDARKACGDSTLTQITAGLDPVGRIQMRTRRTLRGHLAKIYAMHWGTDSRLL
Humanbeta1        AEQLKNQIRDARKACADATLSQITNNIDPVGRIQMRTRRTLRGHLAKIYAMHWGTDSRLL
Mousebeta1        AEQLKNQIRDARKACADATLSQITNNIDPVGRIQMRTRRTLRGHLAKIYAMHWGTDSRLL
Humanbeta4        AEQLRNQIQDARKACNDATLVQITSNMDSVGRIQMRTRRTLRGHLAKIYAMHWGYDSRLL
Mousebeta4        AEQLRNQIQDARKACNDATLVQITSNMDSVGRIQMRTRRTLRGHLAKIYAMHWGYDSRLL
Drosophilabeta1   AESLKNAIRDARKAACDTSLLQAATSLEPIGRIQMRTRRTLRGHLAKIYAMHWGNDSRNL
Celegansbeta1     AEQLKSQIREARKSANDTTLATVASNLEPIGRIQMRTRRTLRGHLAKIYAMHWASDSRNL
1184              SEQLKNQIREARKAAADTTLAQATSSIEAVGRIQMRTRRTLRGHLAKIYAMHWSSDSRNL
1615              CEQLKNQIREARKAAADTSLAQASSDIDSVGRIQMKTRRTLRGHLAKIYAMHWGTDSRNL
10481             ADTLKKKLVEDRNRLKDGDLATLSQKLEPVPSLNIKVRRLLKGHQGRVLSLAWSYDKRHL
Celegansbeta5     AEELRKKLDQERHKLNDIPIQQAAERLDVMGALGVKQRRILKGHVGKVLCMDWSLDKRHI
Drosophilabeta5   AENLKTKLEEERQKLNDVNLSNIAERLEQIAYVNIKPRKVLKGHQAKVLCTDWSPDKRHI
Humanbeta5        AESLKGKLEEERAKLHDVELHQVAERVEALGQFVMKTRRTLKGHGNKVLCMDWCKDKRRI
Mousebeta5        AESLKGKLEEERAKLHDVELHQVAERVEALGQFVMKTRRTLKGHGNKVLCMDWCKDKRRI
```

## WD 2

```
                  β3         β4           β5             β6         β7         β8
Ratbeta1                         TTT                              TT         TT      TT TT
                          80        90        100       110       120       130
Ratbeta1          VSASQDGKLIIWDSYTTNKVHAIPLRSSWVMTCAYAPSGNYVACGGLDNICSIYNLKTRE
2872              VSASQDGKLFIWDTHTANKIHMIPLEIPWVIGCGISPNMKLVCSGGLDNVISVFNIANI.
3309              VTASQDGKLIVWDTYSGNKLHMIPLNIAWVIGCGFAPGCRFVCSGGLDNVISIFDLQSA.
DrosophilabetaE   VTGSLDGKLIIWDTWTANKVQIIPLRSAWVMTVAFSPSGNFVACGGMDNQCTVYDVNNRD
31894             VSASQDGKLIIWDSYTTNKIHAIPLRSSWVMTCALADSGDYVACGGLDNTCTVYNLRLRE
7509              VSASQDGKLIVWDGFTGYKIHAIPLRSSWVMT............................
Humanbeta3        VSASQDGKLIVWDSYTTNKVHAIPLRSSWVMTCAYAPSGNFVACGGLDNMCSIYNLKSRE
Mousebeta3        VSASQDGKLIVWDTYTTNKVHAIPLRSSWVMTCAYAPSGNFVACGGLDNMCSIYNLKSRE
Mousebeta2        VSASQDGKLIIWDSYTTNKVHAIPLRSSWVMTCAYAPSGNFVACGGLDNICSIYSLKTRE
Humanbeta2        VSASQDGKLIIWDSYTTNKVHAIPLRSSWVMTCAYAPSGNFVACGGLDNICSIYSLKTRE
Humanbeta1        VSASQDGKLIIWDSYTTNKVHAIPLRSSWVMTCAYAPSGNYVACGGLDNICSIYNLKTRE
Mousebeta1        VSASQDGKLIIWDSYTTNKVHAIPLRSSWVMTCAYAPSGNYVACGGLDNICSIYNLKTRE
Humanbeta4        VSASQDGKLIIWDSYTTNKMHAIPLRSSWVMTCAYAPSGNYVACGGLDNICSIYNLKTRE
Mousebeta4        VSASQDGKLIIWDSYTTNKMHAIPLRSSWVMTCAYAPSGNYVACGGLDNICSIYNLKTRE
Drosophilabeta1   VSASQDGKLIVWDSHTTNKVHAIPLRSSWVMTCAYAPSGSYVACGGLDNMCSIYNLKTRE
Celegansbeta1     VSASQDGKLIVWDSYTTNKVHAIPLRSSWVMTCAYAPSGSFVACGGLDNICSIYSLKTRE
1184              VSASQDGKLIVWDGYSTNKVHAIPLRSSWVMTCAYAPSGNYVACGGLDNICSIYSLKTRE
1615              VSASQDGKLIVWDVYTTNKVHAIPLRSSWVMTCAYAPSGGYVACGGLDNICSIYNLKTRE
10481             VSTSQDGKLIVWDGFTNTKEFGLSMATTWVMACAYSPTGAHVACGGLDNKCTIYPLITD.
Celegansbeta5     VSSSQDGKVIVWDGFTTNKEHALTMPTTWVMACAFSPSSQMIACGGLDNKCSVVPLSFE.
Drosophilabeta5   ISSSQDGRLIIWDAFTTNKEHAVTMPTTWIMACAYAPSGNFVACGGLDNKVTVYPITSD.
Humanbeta5        VSSSQDGKVIVWDSFTTNKEHAVTMPCTWVMACAYAPSGCAIACGGLDNKCSVYPLTFDK
Mousebeta5        VSSSQDGKVIVWDSFTTNKEHAVTMPCTWVMACAYAPSGCAIACGGLDNKCSVYPLTFDK
```

# WD 3

```
Ratbeta1            β9  β10  T...T  β11  TT  β12  η1  β13
                    140  150  160  170  180
Ratbeta1            .G.NVRVSRELAGHTGYLSCCRFLDD...NQIVTSSGDTTCALWDIETG.QQTTTFTGHT
2872                .DSLMKPSIELKDHKGYISNCKFIND...SSIISSGDKNCILWDISSGGKAVAQFVGHQ
3309                .SGPEAPCCELNGHDGYISSCRFQND...DTIISASGDKSCGLWNIETN.KMLSSYYGHE
DrosophilabetaE     ASGVAKMVKELMGYEGFLSSCRFLDD...GHLITGSGDMKICHWDLEKG.VKTMDFNGHA
31894               SG.HPKVSRELPGHNGYLSCCRFLGNDEEGYIITSSGDTTCGLWDIEAA.QCTRLFESHT
7509                ............................................................
Humanbeta3          .G.NVKVSRELSAHTGYLSCCRFLDD...NNIVTSSGDTTCALWDIETG.QQKTVFVGHT
Mousebeta3          .G.NVKVSRELSAHTGYLSCCRFLDD...NNIVTSSGDTTCALWDIETG.QQKTVFVGHT
Mousebeta2          .G.NVRVSRELPGHTGYLSCCRFLDD...NQIITSSGDTTCALWDIETG.QQTVGFAGHS
Humanbeta2          .G.NVRVSRELPGHTGYLSCCRFLDD...NQIITSSGDTTCALWDIETG.QQTVGFAGHS
Humanbeta1          .G.NVRVSRELAGHTGYLSCCRFLDD...NQIVTSSGDTTCALWDIETG.QQTTTFTGHT
Mousebeta1          .G.NVRVSRELAGHTGYLSCCRFLDD...NQIVTSSGDTTCALWDIETG.QQTTTFTGHT
Humanbeta4          .G.NVRVSRELPGHTGYLSCCRFLDD...SQIVTSSGDTTCALWDIETG.QQTTTFTGHS
Mousebeta4          .G.NVRVSRELPGHTGYLSCCRFLDD...GQIITSSGDTTCALWDIETA.QQTTTFTGHS
Drosophilabeta1     .G.NVRVSRELPGHGGYLSCCRFLDD...NQIVTSSGDMSCGLWDIETG.LQVTSFLGHT
Celegansbeta1       .G.NVRVSRELPGHTGYLSCCRFLDD...NQIVTSSGDMTCALWDIETG.QQCTAFTGHT
1184                .G.NVRVSRELPGHTGYLSCCRFIDD...NQIVTSSGDVTCGLWDIETG.QQVATFTGHT
1615                .G.NVRVSRELPGHTGYLSCCRFLDD...SQIVTSSGDVTCGLWDIETG.QQLAAFTGHT
10481               .EDPVLKKRLVATHTSYLSCCQFNLS..EYQLLTASGDSTCVLWDVESG.QMIQSFHGHS
Celegansbeta5       .DDIIQKKRQVATHTSYMSCCTFLRS..DNLILTGSGDSTCAIWDVESG.QLIQNFHGHT
Drosophilabeta5     .EEMAAKKRTVGTHTSYMSCCIYPNS..DQQILTGSGDSTCALWDVESG.QLLQSFHGHS
Humanbeta5          NENMAAKKKSVAMHTNYLSACSFTNS..DMQILTASGDGTCALWDVESG.QLLQSFHGHG
Mousebeta5          NENMAAKKKSVAMHTNYLSACSFTNS..DMQILTASGDGTCALWDVESG.QLLQSFHGHG
```

# WD 4 WD 5

```
Ratbeta1            β14  TT  β15  TT  β16  β17  β18  TT
                    190  200  210  220  230  240
Ratbeta1            GDVMSLSLAPD..TRLFVSGACDASAKLWDVRE.GMCRQTFTGHESDINAICFFPN.GNA
2872                NDVTAIALSSSSR.SDMFVSVSSDKSCRIWDIRS.QRCVQIFEGHTEDVNGVEFFPKDNYG
3309                NDVTCLDISRKD.VNIFVTASADKTCRLWDVRIPNRYVQVFEGHEQDVNGISFFPESSYG
DrosophilabetaE     GDIAGLSLSPD..MKTYITGSVDKTAKLWDVRE.EGHKQMFFGHDMDVSSVCYHPS.GFG
31894               GDVMSVDVTRD..NKLFISGACDASVKMWDIRS.GNCVQTFTGHESDINAVTYFPN.EQL
7509                ............................................................
Humanbeta3          GDCMSLAVSPD..FNLFISGACDASAKLWDVRE.GTCRQTFTGHESDINAICFFPN.GEA
Mousebeta3          GDCMSLAVSPD..YKLFISGACDASAKLWDVRE.GTCRQTFTGHESDINAICFFPN.GEA
Mousebeta2          GDVMSLSLAPD..GRTFVSGACDASIKLWDVRD.SMCRQTFIGHESDINAVAFFPN.GYA
Humanbeta2          GDVMSLSLAPD..GRTFVSGACDASIKLWDVRD.SMCRQTFIGHESDINAVAFFPN.GYA
Humanbeta1          GDVMSLSLAPD..TRLFVSGACDASAKLWDVRE.GMCRQTFTGHESDINAICFFPN.GNA
Mousebeta1          GDVMSLSLAPD..TRLFVSGACDASAKLWDVRE.GMCRQTFTGHESDINAICFFPN.GNA
Humanbeta4          GDVMSLSLSPD..MRTFVSGACDASSKLWDIRD.GMCRQSFTGHVSDINAVSFFPN.GYA
Mousebeta4          GDVMSLSLSPD..LKTFVSGACDASSKLWDIRD.GMCRQSFTGHISDINAVSFFPS.GYA
Drosophilabeta1     GDVMALSLAPQ..CKTFVSGACDASAKLWDIRE.GVCKQTFPGHESDINAVTFFPN.GQA
Celegansbeta1       GDVMSLSLSPD..FRTFISGACDASAKLWDIRD.GMCKQTFPGHESDINAVAFFPS.GNA
1184                GDVMSLSLAPD..MRTFVSGACDASAKLWDLRD.GQCKQTFPGHESDINAITFFPN.GQA
1615                GDVMSLSLAPD..HRTFVSGACDASAKLWDLRD.GKCKQTFPGHESDINAVTFFPN.GFG
10481               GDVMSLDLSPSESGRVFISGSCDRCVNVWDMRT.GQCVQVYEGHESDVNSVRFYPS.GDA
Celegansbeta5       GDVFAIDVPKCDTGNTFISAGADKHSLVWDIRS.GQCVQSFEGHEADINTVRFHPN.GDA
Drosophilabeta5     GDVMAIDLAPNETGNTFVSGSCDRMAFIWDMRS.GHVVQSFEGHQSDVNSVKFHPC.GDA
Humanbeta5          ADVLCLDLAPSETGNTFVSGGCDKKAMVWDMRS.GQCVQAFETHESDINSVRYYPS.GDA
Mousebeta5          ADVLCLDLAPSETGNTFVSGGCDKKAMVWDMRS.GQCVQAFETHESDVNSVRYYPS.GDA
```

# WD 6

```
Ratbeta1            β19  TT  β20  β21  β22  TT  β23  TT  β24  TT
                    250  260  270  280  290  300
Ratbeta1            FATGSDDATCRLFDLRADQELMTYSHDNIICGITSVSFSKSGRLLLAGYDDFNCNVWDAL
2872                FVTSSDDGTCRLWDSRADQSIAVYTDDYITCGSTSVSLSKSGRLLIAGYDDYNCHVWDVL
3309                FVSASDDKACRLWDIRSDQCIAIYTDDYIKSGSTSVTISKSGRVLIAGYDDFICHVWDLL
DrosophilabetaE     FASCSEDQTARMYDLRADQQIAQYEPPQKNTGFTSCALSTSGRYLMCGGIEGNVHSWDTM
31894               IGTASDDATCRLFDIRADQELALYSLDSIICGITSIQFSLSGRMLFAGYDDFNCQVWDTL
7509                ............................................................
Humanbeta3          ICTGSDDASCRLFDLRADQELICFSHESIICGITSVAFSLSGRLLFAGYDDFNCNVWDSM
Mousebeta3          ICTGSDDASCRLFDLRADQELTAYSQESIICGITSVAFSLSGRLLFAGYDDFNCNVWDSL
Mousebeta2          FTTGSDDATCRLFDLRADQELLMYSHDNIICGITSVAFSRSGRLLLAGYDDFNCNIWDAM
Humanbeta2          FTTGSDDATCRLFDLRADQELLMYSHDNIICGITSVAFSRSGRLLLAGYDDFNCNIWDAM
Humanbeta1          FATGSDDATCRLFDLRADQELMTYSHDNIICGITSVSFSKSGRLLLAGYDDFNCNVWDAL
Mousebeta1          FATGSDDATCRLFDLRADQELMTYSHDNIICGITSVSFSKSGRLLLAGYDDFNCNVWDAL
Humanbeta4          FATGSDDATCRLFDLRADQELLLYSHDNIICGITSVAFSKSGRLLLAGYDDFNCNVWDTL
Mousebeta4          FATGSDDATCRLFDLRADQELLLYSHDNIICGITSVAFSKSGRLLLAGYDDFNCSVWDAL
Drosophilabeta1     FATGSDDATCRLFDIRADQELAMYSHDNIICGITSVAFSKSGRLLLAGYDDFNCNVWDTM
Celegansbeta1       FATGSDDATCRLFDIRADQELAMYSHDNIICGITSVAFSKSGRLLFAGYDDFNCNVWDSM
1184                FATGSDDATCRLFDIRADQEIGMFSHDNIICGITSVAFSKSGRLLLGGYDDFNCNVWDTL
1615                FATGSDDATCRLFDIRSDQEIGMYSNDSIICGITSVAFSKSGRLLLGGYDDFNCNVWDTL
10481               FATGSDDATCRLFDLRADSEICVYKKDSVLFGCNAVDFSLSGRLLFAGYNDYIMNIWDVL
Celegansbeta5       FATGSDDATCRLFDLRADRQVCVYEKESILFPVNGVDFSLSGRILFAGYGDYRVGVWDSL
Drosophilabeta5     IATGSDDSSCRLYDMRADREVAVFAKESIIFGVNSVDFSVSGRLLFAGYNDYTVNLWDTL
Humanbeta5          FASGSDDATCRLYDLRADREVAIYSKESIIFGASSVDFSLSGRLLFAGYNDYTINVWDVL
Mousebeta5          FASGSDDATCRLYDLRADREVAIYSKESIIFGASSVDFSLSGRLLFAGYNDYTINVWDVL
```

## WD 7

```
                      β25          β26          β27        β28
Ratbeta1            T                      TT           TT
                                 310         320          330          340
Ratbeta1            KADRAGVLAGHDNRVSCLGVTDDGMAVATGSWDSFLKIWN......
2872                REERVGIMSAHDGRVSCVSVSPNGVGIATGSWDSSCLIWTSKPSQ.
3309                REERVGLLSAHDGRISGVKVSPDGVAIATCSWDTTCLVWTARKKGK
DrosophilabetaE     KQRHTGTLSGHENRITCISLCPNGMCLASTSWDQQVRLWL......
31894               KQDRIGVLASHDNRVSCLGVSKNGEALCTGSWDSTLKIWN......
7509                ..............................................
Humanbeta3          KSERVGILSGHDNRVSCLGVTADGMAVATGSWDSFLKIWN......
Mousebeta3          KCERVGILSGHDNRVSCLGVTADGMAVATGSWDSFLKIWN......
Mousebeta2          KGDRAGVLAGHDNRVSCLGVTDDGMAVATGSWDSFLKIWN......
Humanbeta2          KGDRAGVLAGHDNRVSCLGVTDDGMAVATGSWDSFLKIWN......
Humanbeta1          KADRAGVLAGHDNRVSCLGVTDDGMAVATGSWDSFLKIWN......
Mousebeta1          KADRAGVLAGHDNRVSCLGVTDDGMAVATGSWDSFLKIWN......
Humanbeta4          KGDRAGVLAGHDNRVSCLGVTDDGMAVATGSWDSFLRIWN......
Mousebeta4          KGGRSGVLAGHDNRVSCLGVTDDGMAVATGSWDSFLRIWN......
Drosophilabeta1     KAERSGILAGHDNRVSCLGVTENGMAVATGSWDSFLRVWN......
Celegansbeta1       RQERAGVLAGHDNRVSCLGVTEDGMAVCTGSWDSFLKIWN......
1184                KQDRAGVLAGHDNRVSCLGVSEDGMAVCTGSWDSFLRIWN......
1615                KQDRAGVLAGHDNRVSCLGVSEDGMAVCTGSWDSFLRVWN......
10481               KSNRVAILNGHENRISCLKTSPDGTAVCTGSWDSTLRIWA......
Celegansbeta5       KCARHSVLYGHENRISCLRTSPDGTAVCSASWDCTIRIWA......
Drosophilabeta5     KSERVCLLYGHENKVSCVQVSPDGTALSTGSWDYTIRVWA......
Humanbeta5          KGSRVSILFGHENRVSTLRVSPDGTAFCSGSWDHTLRVWA......
Mousebeta5          KGSRVSILFGHENRVSTLRVSPDGTAFCSGSWDHTLRVWA......
```

```
                                              α1                        α2
CattleGNG2 [G2]
                           1       10        20                  30
CattleGNG2 [G2]     ......MASNNTASIAQARKLVEQLK..........MEANIDRIKV........SKAAAD
1191                M........NKSSTVSNDRKLSKNIKKEAKENFVNRLQIKINKKQINSQLADKLDETTEM
1090                .....MQPDKVIRKMAELNKEISILK..........NNLRVKRIPV........SQAVQE
2571                ........MPAKRDVAKAMVELETLR..........ESSQVKRLPL........SVTLKE
1237                ...........MSGISQQKKLVEQLR..........AEAAINRRPV........SASVQE
2883                MSAAATEENPRQNEIDMKARELEYLK..........NILKLERLQL........SKTIKD
HumanGNGT1 [Gt1]    MP...VINIEDLTEKDKLKMEVDQLK..........KEVTLERMLV........SKCCEE
HumanGNG2 [G2]      ......MASNNTASIAQARKLVEQLK..........MEANIDRIKV........SKAAAD
HumanGNG3 [G3]      MK..GETPVNSTMSIGQARKMVEQLK..........IEASLCRIKV........SKAAAD
HumanGNG4 [G4]      MK..EGMSNNSTTSISQARKAVEQLK..........MEACMDRVKV........SQAAAD
HumanGNG5 [G5]      ........MSGSSSVAAMKKVVQQLR..........LEAGLNRVKV........SQAAAD
HumanGNG7 [G7]      ........MSATNNIAQARKLVEQLR..........IEAGIERIKV........SKAASD
HumanGNG8 [G8]      .......MSNNMAKIAEARKTVEQLK..........LEVNIDRMKV........SQAAAE
HumanGNG10 [G10]    ........MSSGASASALQRLVEQLK..........LEAGVERIKV........SQAAAE
HumanGNG11 [G11]    MP...ALHIEDLPEKEKLKMEVEQLR..........KEVKLQRQQV........SKCSEE
HumanGNG12 [G12]    ....MSSKTASTNNIAQARRTVQQLR..........LEASIERIKV........SKASAD
HumanGNG13 [G13]    .........MEEWDVPQMKKEVESLK..........YQLAFQREMA........SKTIPE
HumanRGS7           SPTHTPTPETKPPTEDELQQQIKYWQ..........IQLDRHRLKM........SKVADS

CattleGNG2 [G2]                        TTT
                       40        50         60         ☆70
CattleGNG2 [G2]     LMAYCEAHAKEDPLLTPVP.ASENPFREKKF..FCAIL......................
1191                ILENCNQEQSN.......PSNKEKPKKQKKL..KNLEIEK...IINKNITSFGHSYENIF
1090                LIIYCNENSTHDIWMNP...LIKSPFSSTHWSICGML.......................
2571                LIEFCNTNYKVDFFLNP...MEENPFKSHRKPICSII.......................
1237                MIQYMDENKDKDCLVSGFANKKDNPFQEKGG..CIIL.......................
2883                FMEYCYNNANSDPLLYP...LKENPFLEKKS..CIIL.......................
HumanGNGT1 [Gt1]    VRDYVEERSGEDPLVKGIP.EDKNPFKELKG..GCVIS......................
HumanGNG2 [G2]      LMAYCEAHAKEDPLLTPVP.ASENPFREKKF..FCAIL......................
HumanGNG3 [G3]      LMTYCDAHACEDPLITPVP.TSENPFREKKF..FCALL......................
HumanGNG4 [G4]      LLAYCEAHVREDPLIIPVP.ASENPFREKKF..FCTIL......................
HumanGNG5 [G5]      LKQFCLQNAQHDPLLTGVS.SSTNPFRPQKV..CSFL.......................
HumanGNG7 [G7]      LMSYCEQHARNDPLLVGVP.ASENPFKDKKP..C.IIL......................
HumanGNG8 [G8]      LLAFCETHAKDDPLVTPVP.AAENPFRDKRL..FCVLL......................
HumanGNG10 [G10]    LQQYCMQNACKDALLVGVP.AGSNPFREPRS..CALL.......................
HumanGNG11 [G11]    IKNYIEERSGEDPLVKGIP.EDKNPFKE.KG..SCVIS......................
HumanGNG12 [G12]    LMSYCEEHARSDPLLIGIP.TSENPFKDKKT..C.IIL......................
HumanGNG13 [G13]    LLKWIEDGIPKDPFLNPDL.MKNNPWVEKGK..CTIL.......................
HumanRGS7           LLSYTEQYLEYDPFLLPP..DPSNPWLSDDT..TFWELEASKEPSQQRVKRWGFGMDEAL

CattleGNG2 [G2]

CattleGNG2 [G2]     ............................................................
1191                ............................................................
1090                ............................................................
2571                ............................................................
1237                ............................................................
2883                ............................................................
HumanGNGT1 [Gt1]    ............................................................
HumanGNG2 [G2]      ............................................................
HumanGNG3 [G3]      ............................................................
HumanGNG4 [G4]      ............................................................
HumanGNG5 [G5]      ............................................................
HumanGNG7 [G7]      ............................................................
HumanGNG8 [G8]      ............................................................
HumanGNG10 [G10]    ............................................................
HumanGNG11 [G11]    ............................................................
HumanGNG12 [G12]    ............................................................
HumanGNG13 [G13]    ............................................................
HumanRGS7           KDPVGREQFLKFLESEFSSENLRFWLAVEDLKKRPIKEVPSRVQEIWQEFLAPGAPSAIN

CattleGNG2 [G2]

CattleGNG2 [G2]     ............................................................
1191                ............................................................
1090                ............................................................
2571                ............................................................
1237                ............................................................
2883                ............................................................
HumanGNGT1 [Gt1]    ............................................................
HumanGNG2 [G2]      ............................................................
HumanGNG3 [G3]      ............................................................
HumanGNG4 [G4]      ............................................................
HumanGNG5 [G5]      ............................................................
HumanGNG7 [G7]      ............................................................
HumanGNG8 [G8]      ............................................................
HumanGNG10 [G10]    ............................................................
HumanGNG11 [G11]    ............................................................
HumanGNG12 [G12]    ............................................................
HumanGNG13 [G13]    ............................................................
HumanRGS7           LDSKSYDKTTQNVKEPGRYTFEDAQEHIYKLMKSDSYPRFIRSSAYQELLQAKKKSGNSM
```
