## Supplementary figures and images for "Heterotrimeric G proteins regulate planarian regeneration and behavior"

### Supp. File 2

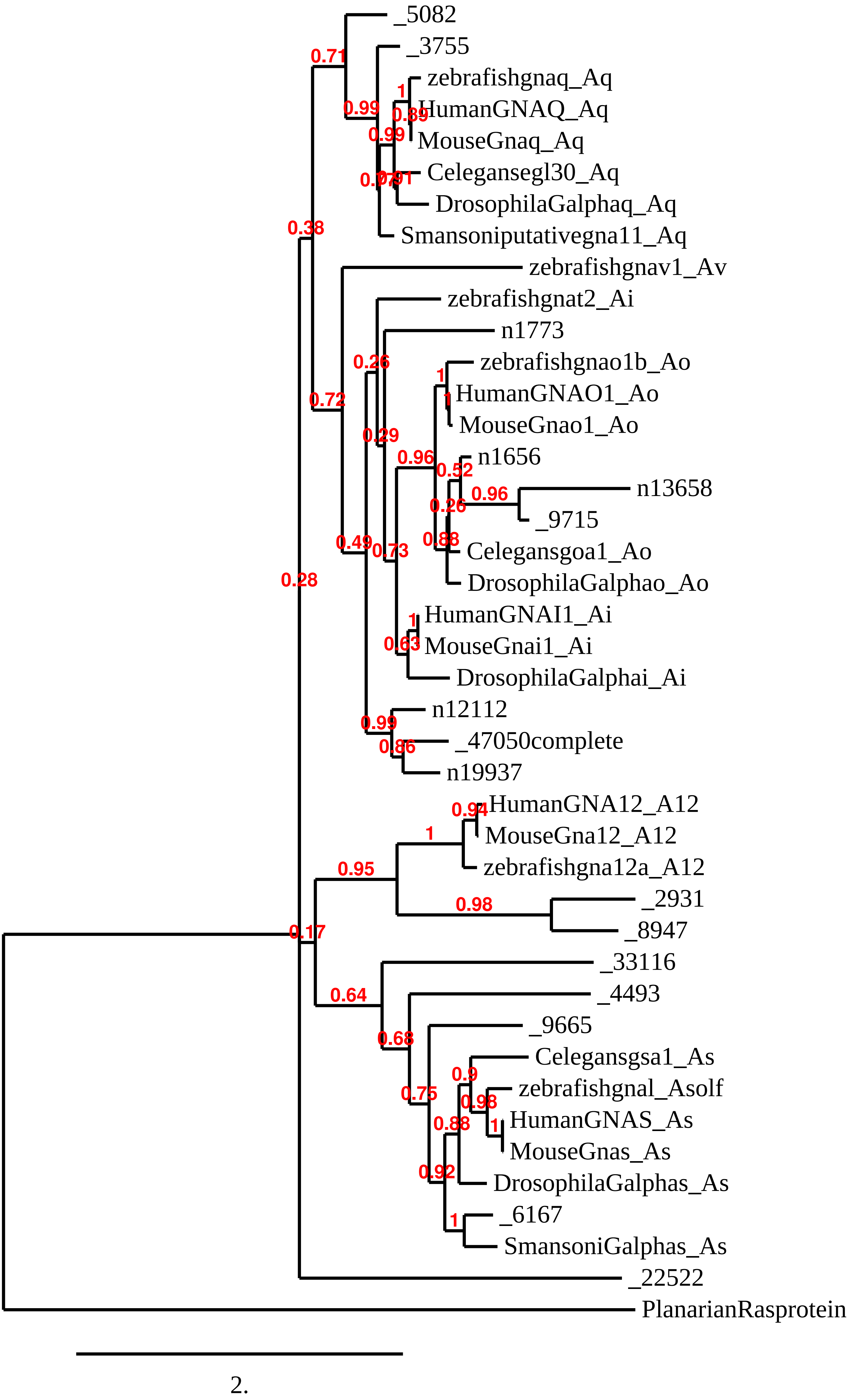

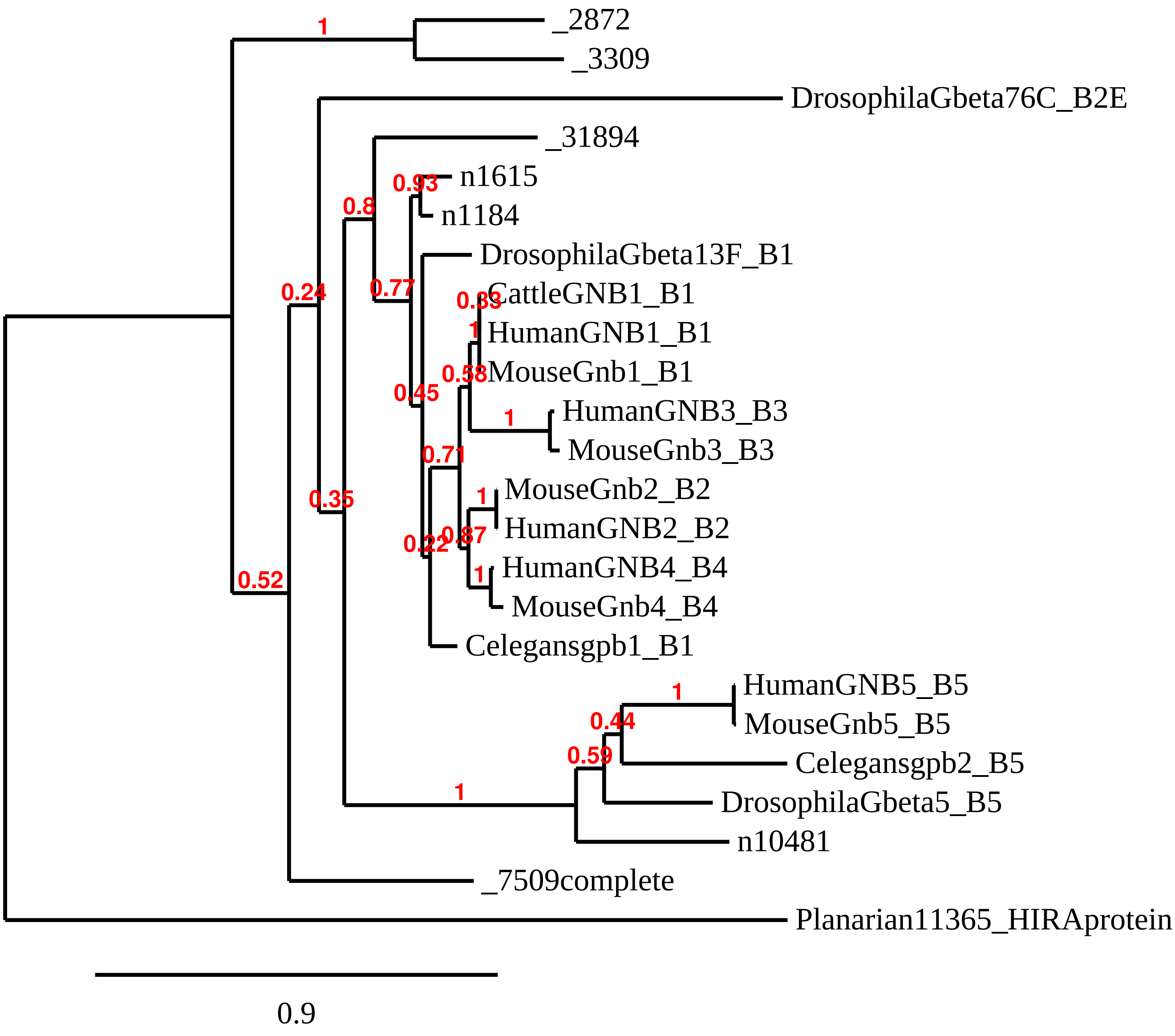

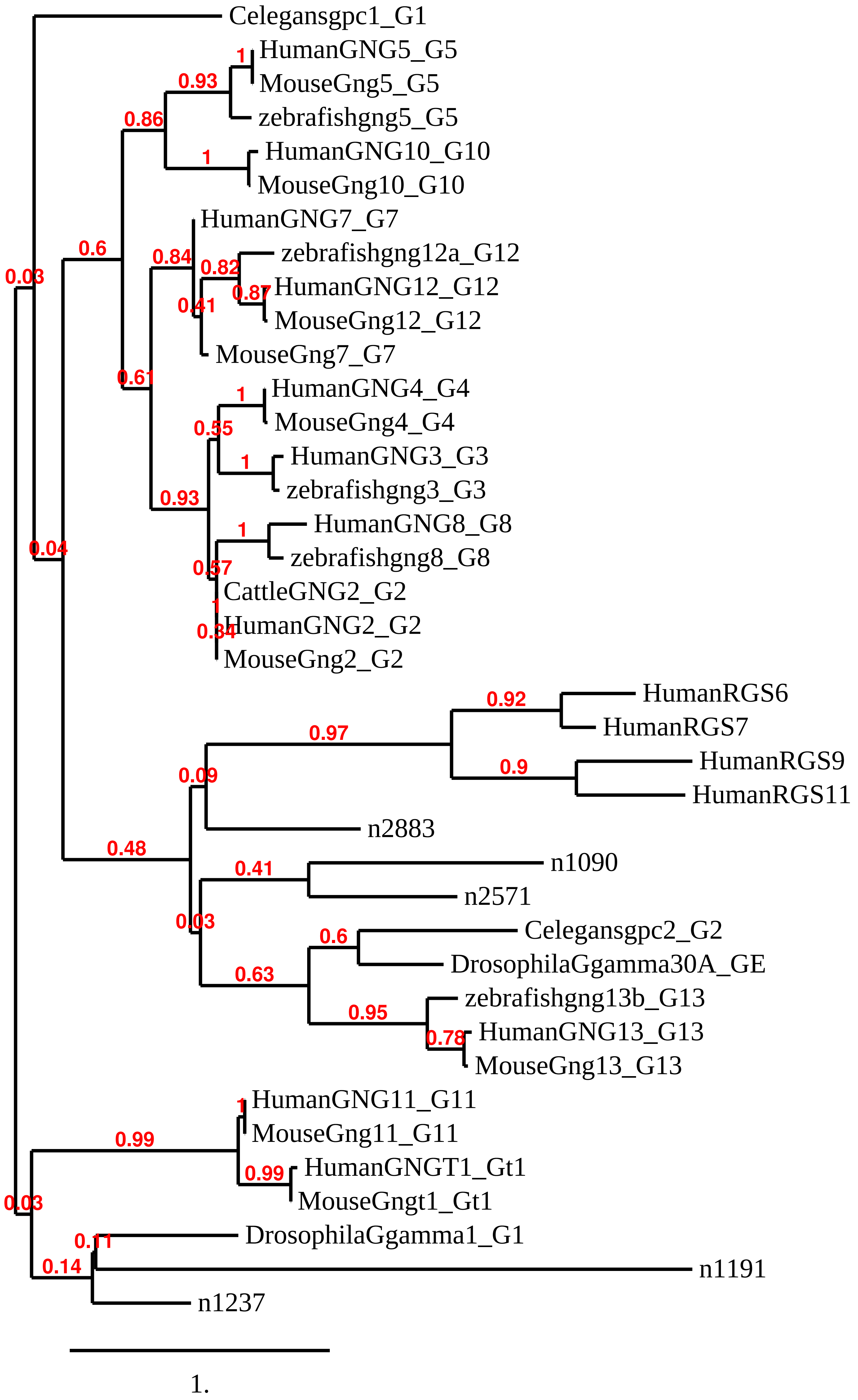
