## Supplementary material for "Heterotrimeric G proteins regulate planarian regeneration and behavior": Supp. File 3

```
                                                α1              β1
RatGnai1[Ai]                                 .        
                      1                  10        20        30
RatGnai1[Ai]        ........MGCTLS...............AEDKA.AVERSKMIDRNLREDGEKAAREVKL
82535               ............................................................
1635                ................................................MVSRIGTPVVAI
16751               ............................................................
2931                .......MLPCIPK.RSRFIG........DVDIKHQKAKSREIDKLMKREKRKFLTRHSI
8947                .......MFTCFKNSKKKFYK........EIDLKSQKSFSKLIDKSLKIERRKNLMKQHI
33116               ............................................................
13658               ............................................................
23262               ............................................................
32381               ............................................................
21781               ............................................................
86418               ............................................................
9715                ........MGCAMS................LDERIALNKSKDIDKKLKEDGLQLQRDIKL
94774               ............................................................
1656                ........MGCTMS................KDERDALDKSRNIDKKLKEDGMQAAKDVKL
22522               ........MNCNFKISSHLNTIN......QISTQKSVLRSREIDQQLMQDQELRKKEIQI
9665                ......MPAICGGKSA.................STEDKSNKEIDKKLKKDKKTFLMTHRI
6167                ......MALSCAAF...KTET.........EEQKNRKESNKRIEKCIEKEKRSFKSTHRL
5082                ........MACCNS...............EEFKE.QSRINKEIEKQLKRDKKDARRELKL
3755                ........MGCCLS...............ELDKE.QKRINKEIEKQLKKDKKESRRELKL
1773                ........MGCPLG...............KPDEIEQNEISHKIDKELKLEAKHSYRIVKL
77832               ........MGCLNT...............TLDQD.QQDRNREIEKQIRCDAEKAMKEVKL
47050complete       ........MGCINS...............SHDEP.QQELNKNIEKQLRIDAEKAMKEVKL
19937               ........MGCINS...............TKNVI.QEEAHKEIEKQLRIDQEKAMREIKL
12112               ........MGCAGS...............VPDKD.QQEKNKEIEQQLRQDAEKAMKEVKL
65197               ............................................................
4493                .......MNTCISKVETDEQREK......RVLLEENRRKNNQIEKELALYKNKKLSCLKL
HumanGNAO1[Ao]      ........MGCTLS................AEERAALERSKAIEKNLKEDGISAAKDVKL
HumanGNAS[As]       ........MGCLGNSKTEDQR.........NEEKAQREANKKIEKQLQKDKQVYRATHRL
HumanGNA12[A12]     MSGVVRTLSRCLLPAEAGGARERRAGSGARDAEREARRRSRDIDALLARERRAVRRLVKI
HumanGNAQ[Aq]       ..MTLESIMACCLS...............EEAKE.ARRINDEIERQLRRDKRDARRELKL
HumanGNAI1[Ai]      ........MGCTLS...............AEDKA.AVERSKMIDRNLREDGEKAAREVKL
```

```
                                α2              α3              α4
RatGnai1[Ai]          →    TT
                          40        50        60        70        80        90
RatGnai1[Ai]        LLLGAGESGKSTIVKQMKII..HEAGYSEEE.CKQYKAVVYSNTIQSIIAIIRAMGRLKI
82535               ............................................................
1635                MLISSG...SDRCMEDFDIM..............KFGVTTLNNMD...............
16751               ............................................................
2931                LLLGTGESGKSTFLKQMKLI..NGKKFTSAE.LAGFKDTIYDNVYKGILFLLHARSSLNI
8947                LLLGTGESGKSTFLKQMKII..CGQKFTVYE.IESFKDIIYDNIFKGVLFLIDARAQLNI
33116               ................MKLLSSINQTFPDVY.RAKFIPEIQRNLVQALTAILQFMDQIKL
13658               ...............................................MIDILYAVHHLNI
23262               ......................................................MTTLNI
32381               ............MAKQIKII..HEGGFKNED.NKQYKPVVRSKTTQSITAILRTITILNI
21781               ............MAKHMTII..YKVGVTNED.NKQYKTVVYNKTPQSMIAIL........
86418               ................MKII..HESGFTNED.NKQYKPVIYNKTIQSMRAILRAITTLNI
9715                LLLGAGESGKSTIMKQMKII..HEGGFSSTD.NHHYKPIVYSNTIQSVAAILRAMHKLNI
94774               ................MKIT..QEGGFTNEY.NNQYKLVVYSSTFQ..............
1656                LLLGAGESGKSTIVKQMKII..HEGGFTQED.NKQYKPVVYSNTTQSMIAILRAMTTLGI
22522               LVLGNRNSGKTTFLKQFRIH..HGDGYPYLQ.RAILVPDILSNLADAIHIVMDNMARWDL
9665                PVVGAGESGKSTLIKQMQIL..YINGFNDEE.RKSKIKDIRGNIKDGMLAVTGAMKTLKP
6167                LLLGAGESGKSTIVKQMRIL..HIDGFSNEE.KKQKAEDIRKNLRDAILTITSAMSNINP
5082                LLLGTGESGKSTFIKQMRII..HGTGYSDDD.KRSFIKLIYQNIYLATYTLIRAMEVLKI
3755                LLLGTGESGKSTFIKQMRII..HGSGYSDED.KRSFIKLVYQNIYLAIYTLILAMENLAI
1773                LLLGTGESGKSTIVKQMKLI..VNNGHSQAE.RLGFRSLIFSNTIQSLIVIIRAMSKLDI
77832               LLLGTGECGKSTILKQMTII..HGKGFSEDD.RRQYIPIIISNLIDSINVIINAMDSLEI
47050complete       LLLGAGECGKSTILKQMSII..HGDGYSESE.RKKFIPIIYSNIIQSVVTILKAMETLEI
19937               LLLGAGECGKSTILKQMSII..HGGGYSDEE.KKEFIPIIYANIVQSMGAILKAMDILEI
12112               LLLGAGECGKSTILKQMTII..HGKGYPEEE.RKEYVPIIYANVVQSMVTILNAMETLGI
65197               ............................................................
4493                LLLGTGESGKSTILKQMKII..HVNGFSKTE.KIEFISNIKRNVRDAIMAIVVSMNKLQI
HumanGNAO1[Ao]      LLLGAGESGKSTIVKQMKII..HEDGFSGED.VKQYKPVVYSNTIQSLAAIVRAMDTLGI
HumanGNAS[As]       LLLGAGESGKSTIVKQMRIL..HVNGFNGDSEKATKVQDIKNNLKEAIETIVAAMSNLVP
HumanGNA12[A12]     LLLGAGESGKSTFLKQMRII..HGREFDQKA.LLEFRDTIFDNILKGSRVLVDARDKLGI
HumanGNAQ[Aq]       LLLGTGESGKSTFIKQMRII..HGSGYSDED.KRGFTKLVYQNIFTAMQAMIRAMDTLKI
HumanGNAI1[Ai]      LLLGAGESGKSTIVKQMKII..HEAGYSEEE.CKQYKAVVYSNTIQSIIAIIRAMGRLKI
```

```
                              α5                                 α6
RatGnai1[Ai]
                    100           110                120

RatGnai1[Ai]       DFGD..AARADDARQLFVLAGA.A..........E.EGFMTA................EL
82535              ............................................................
1635               ......PTT...HIYLIA..........................................
16751              ............................................................
2931               NFENYEETSEAEIRI.EDHFKRNKEIHKEKA...TTGQIIWS...........EEEFLKL
8947               DWNDNETAN.SAEEI.ERYFDETKLIHRQRR...QSKQLLWK...........ENEFLEI
33116              DFKHPNAHLHTAKKELYEIRDQI...DKNP............DTIEKYANSTEMDVRNNF
13658              HFES..SAREADEILVNNTITTG...........MDEKPLNA................QL
23262              SFGD..SDRLPDAKIGSDVSQAM...........EVIEPWFE................EF
32381              SFGY..PDRSADSKIGYDVIQAM...........KSLNPFSR................SF
21781              ............................................................
86418              FFGD..PYRSADANIRSDVIKAM...........EVNEPCFE................EF
9715               SFEN..SAREADEILVNNIMTTM...........KDQEPFTP................QL
94774              ............................................................
1656               SFGD..IDRGADAKVVSDVIQAM...........EDTEPFSE................EL
22522              RFED..PYVQTLAKDFHNNCPRSPALSNNPHNLFKSIQNNSPRPFNRF..CKTVNRITDF
9665               PVDLEHAENQTLLDFIQHNAV.............KPDFTYTK................EF
6167               PTKLGNPENQKFLDYMQHTAS.............KPDFRYPS................EF
5082               PYEN..PDNLEYAKDLRDI..D.Y..........ETVTTFEP................SH
3755               SYSN..PENLSYIDEIKNI..D.Y..........ESVSTLEP................DH
1773               SFTN..PERLNDAQTLLQLAGT.V..........ESNGPLNE................EL
77832              SL..........................................................
47050complete      KLNE...HLHSDKLLVIDQVKD.V..........D.NGVLRQ................DA
19937              PLEN..ANLGTEKNTVHESMKN.A..........E.EGGMTI................NT
12112              SMEN..EAMIEESKIVQKYMRN.A..........E.EGEFPK................EL
65197              ............................................................
4493               TPKK..FDLSEKIKFLQTAF..............SEQYTYPN................TF
HumanGNAO1[Ao]     EYGD..KERKADAKMVCDVVSRM...........EDTEPFSA................EL
HumanGNAS[As]      PVELANPENQFRVDYILSVMN.............VPDFDFPP................EF
HumanGNA12[A12]    PWQY..SENEKHGMFLMAFENK.AGL........PVEPATFQ................LY
HumanGNAQ[Aq]      PYKY..EHNKAHAQLVREV..D.V..........EKVSAFEN................PY
HumanGNAI1[Ai]     DFGD..SARADDARQLFVLAGA.A..........E.EGFMTA................EL
```

```
                              α7                   α8      α9          α10
RatGnai1[Ai]
                        130           140           150       160       170       180

RatGnai1[Ai]       AGVIKRLWKDSGVQACFNRSREY...QLNDSAAYYLNDLDRIAQPNYIPTQQDVLRTRVK
82535              .......................MKMSFNNSCCFHSGTQNI.......LLGWDICRFNYP
1635               .............VCCHVANERFRWPLMIGYSLYFLSSIEKFSVTDNFL.......CHSF
16751              ...................................................MGRFRCPRM
2931               VDDFKLIWNDKGIKEAFNQRSKLMTESFSENSRFYLNKLDKIRVKDYQPSNEDIVWSRKP
8947               VQRLKAIWNDKSIQQTFLRRSEIITESFSENTRYYLNKIDEIGTLNYFPTDEDIVWTRKP
33116              YDNCKALWRDPNILDTFQRSNEY...QLIDSAQYFLDKIDLIRQPDYKPSDDDVLQCRTK
13658              CNALKSLWSDVGVQECFRTFNEY...DLNYSTQYFLDDIDRISAVDYEPTVQDILRSKMT
23262              LKASKRLCADSCVQECFNLSNEY...QLSDSAKYFHDDINQLGAGDYMPTTHSNEIQHGS
32381              ............................................................
21781              ............................................................
86418              ............................................................
9715               NSALKSLWSDIGVQECFRRSNEY...QLNDSAQYFLDNVDRISAVDYKPSDQDILRTRIK
94774              ............................................................
1656               LEAMKRLWADPGVQECFGRSNEY...QLNDSAKYFLDDLDRLGAKDYMPTEQDILRTRVK
22522              CLHIKNISDQDDFQECLDCKEFF.KPSITYADLYFIQNIDRLIQADYTPTLQDIMVMRKP
9665               FDYCYTLWIDAGVQATFARANEY...QLIDSTKYFLDQVQTIAKDDYLPTDQDLLRCRVL
6167               YEYCAVLWADKGVLETYERSNEY...QLIDCAKYFLDQALILGQQNYTPTEQDILRCRVL
5082               IVATRSLWNDSGVKECYDRRREY...QLTDSAKYYLDDLDRIVTPDYLPTLQDILRVRVP
3755               VIAIKSLWADVGVKECYDRRREF...QLTDSAKYYLDSLDRIATPNFLPTLQDILRVRVP
1773               YSAMLNLWQDLGIQECFQRSAEY...QLIDSAEYYLNSLERLSNSNYVPTEQDILRTRVK
77832              ............................................................
47050complete      VESIKKLWQNVEVQQCVQRAKEY...NLNDSAEYYLNDIDRLRGDNYLPNDQDILRSRVK
19937              IKALTVLWKDTGVQEAFNRSKEY...QLNDSAEYYLIDLERIASDNYIPTVQDILRSRIK
12112              GSALALLWKDDSVQNCYARAKEY...QLNDSAGYYLDSLDRLSEPSYIPTEQDVLRSRVK
65197              ............................................................
4493               FDVVSELWQDEDVQASFLRSNEY...QLIDSAKYFLDHIHIIRQNDYIPSLQDILRCRKM
HumanGNAO1[Ao]     LSAMMRLWGDSGIQECFNRSREY...QLNDSAKYYLDSLDRIGAADYQPTEQDILRTRVK
HumanGNAS[As]      YEHAKALWEDEGVRACYERSNEY...QLIDCAQYFLDKIDVIKQADYVPSDQDLLRCRVL
HumanGNA12[A12]    VPALSALWRDSGIREAFSRRSEF...QLGESVKYFLDNLDRIGQLNYFPSKQDILLARKA
HumanGNAQ[Aq]      VDAIKSLWNDPGIQECYDRRREY...QLSDSTKYYLNDLDRVADPAYLPTQQDVLRVRVP
HumanGNAI1[Ai]     AGVIKRLWKDSGVQACFNRSREY...QLNDSAAYYLNDLDRIAQPNYIPTQQDVLRTRVK
```

```
                         β2              β3           α11   η1          β4       η2
RatGnai1[Ai]          ────────►T.....T────────►     ℓℓℓℓℓℓℓℓ        ────────►ℓℓℓℓℓ
                            190           200          210         220         230   ★

RatGnai1[Ai]          TTGIVETHFTFK.....DLHFKMFDVGGQRSERKKWIHCF.EGVTAIIFCVALSDYDLVL
82535                 FNGVQEVSCG..................................................
1635                  RSP.........................................................
16751                 PQSVKRI.....................................................
2931                  TDSIIEEEIDIH....GA.CIVFIDVGGQTKERKKWCQVF.QDMNSVLFLIASSHFDEFY
8947                  TESIIETEIEVS....NDIILVFIDVGGQTKERRKWCQLF.HDMKSVLFLIACSHFDEEY
33116                 TLGIHTEIINYN.....NVNFELVDVGGQRDQRAKWIEALSDGVTAVIFLTDVSAYDMVL
13658                 NYGVCECQIKYK.....ERMFTIFDTSVRL.SDRKWRICF.EDPSALIFCVSLTEYLK..
23262                 ............................................................
32381                 ............................................................
21781                 ............................................................
86418                 ............................................................
9715                  TTGIVEIQFKFK.....NMNFKIFDVGGQRSERKKWIHCF.EDVTAIIFCVSLSEYDQVL
94774                 ............................................................
1656                  TTGIVEVHFMFK.....NMNFKLFDVGGQRSERKKWIHCF.EDVTAIIFCVAMSEYDQVL
22522                 TYSVKDFIFNLG.....GITLRFIDLGGYKCQRKKFIHYF.EGVSTVFYFCALSEYSEII
9665                  TSGIFETKFTMN.....KVCFHMFDVGGQREERRKWIQVF.SDVTAVIFVCACSGYDMTL
6167                  TSGIFETKFTVD.....KVNFHMFDVGGQRDERRKWIQCF.NDVTAIIFVTACSSYNMVL
5082                  TTGIIEYPFDLD.....SIIFRMVDVGGQRSERRKWIHCF.EGVMTMIYVASISEYDQFL
3755                  TTGIIEYPFDLD.....SIIFRMVDVGGQRSERRKWIHCF.ENVTSIMFLVALSEYDQGL
1773                  TAGIIETTVEFK.....ELIMKIVDVGGQRAERKKWIHCF.DNVDAIIFIVALNEYDLRL
77832                 ............................................................
47050complete         TTGIIETEFHFK.....NLDFKVYDVGGQRSERKKWMHCF.EGVTAILFLIAMSEYDLKL
19937                 TTGIIETYFHFK.....DLDFKVFDVGGQRSERKKWMHCF.EGVTAILFLVALSEYDLRL
12112                 TTGIVETHFHFK.....DLDFKVFDVGGQRSERKKWMHCF.EGVTAILFLVAMSEYDLKL
65197                 ....................................MHFF.EGVTAILFLVAMSEYDLKL
4493                  TSGITEISFEVREKKKNKVNFKVFDVGGQQGERKRWIQLF.GEVTAILFLADCSSFDQTL
HumanGNAO1[Ao]        TTGIVETHFTFK.....NLHFRLFDVGGQRSERKKWIHCF.EDVTAIIFCVALSGYDQVL
HumanGNAS[As]         TSGIFETKFQVD.....KVNFHMFDVGGQRDERRKWIQCF.NDVTAIIFVVASSSYNMVI
HumanGNA12[A12]       TKGIVEHDFVIK.....KIPFKMVDVGGQRSQRQKWFQCF.DGITSILFMVSSSEYDQVL
HumanGNAQ[Aq]         TTGIIEYPFDLQ.....SVIFRMVDVGGQRSERRKWIHCF.ENVTSIMFLVALSEYDQVL
HumanGNAI1[Ai]        TTGIVETHFTFK.....DLHFKMFDVGGQRSERKKWIHCF.EGVTAIIFCVALSDYDLVL


                                  α12              η3      β5         α13          η4
RatGnai1[Ai]                ℓℓℓℓℓℓℓℓℓℓℓℓℓℓℓℓ      ℓℓℓ  ────────►  ℓℓℓℓℓℓℓℓℓℓℓ       ℓℓℓ
                      ★★★  240       250       260       270       280

RatGnai1[Ai]          AED...EEMNRMHESMKLFDSICNNKWFTDTSIILFLNKKDLFEEKIKK..SPLTICYPE
82535                 ............................................................
1635                  .........N..................................................
16751                 ...............................PIKRFS.......................
2931                  YDKMSKEYRNKLREAMRVFEDLINVQYFLSVSVIIFFNKTDVLTEKVTNKISDIRSEFTE
8947                  MDRFTLQKRNKLKEAMFVFEELINQNAFLRVSVLLFFNKTDILSEKIKSHQSNIGKDFQD
33116                 AED...QTTNRLRESVSLLGQVWTKNPLRDKSIILFLNKQDKLEMKVRKGRTQIETYFPE
13658                 ........EDEMACSMRYFNSIIQFKWFQNSSFILFLNKKDLFKRKLLD..HPITCCFPE
23262                 ............................................................
32381                 ............................................................
21781                 ............................................................
86418                 ............................................................
9715                  VED...DATNRMQESLFLFESICNNVWFLHTSFILFLNKKDLFLEKLQI..CPITFCFPE
94774                 ............................................................
1656                  HED...ETTNRMQESLKLFDSICNNKWFTQTSIILFLNKKDLFAEKIKR..SPLTVCFSE
22522                 .EE...TTKNKLNNSLENFEEIINNKYLWRKDFVIFLNKSDLFKSKVEN..LSINVFFSD
9665                  REA...QKQNRLRECISLFSEVWGNRYLRQTSIILFLNKQDLFVQKLTSGKTKIVDFFPE
6167                  RED...ASQNRLRESLELLKSIWNNRWLRNISVILFLNKQDVLKEKVLAGKSKIEDYFPD
5082                  IED...NEINRMFESIKLFDSVCNNQWFSKCSIILFLNKTDLFKLKIVK..SPLTVCFPE
3755                  VES...DNDNRMEESKALFRTIITYPWFHNASVILFLNKKDLLEEKIIY..SHLVDYFPE
1773                  RED...PEVNRMMESLRLFDSMCNNVVFKDTCMILFLNKRDLFEVKIKK..SPLSICFEE
77832                 ............................................................
47050complete         VED...QTTNRMHESMKLFESICNSQWFTSTSIILFLNKKDLFMKKIEI..SPITICFKD
19937                 VED...STTNRMHESMKLFDSICNSPWFVNTSVILFLNKVDLFEIKIQR..SPLTICFPE
12112                 VED...SSTNRMHESMRLFDSICNSQWFVNTSIILFLNKKDLFGEKVVK..SPLTVCFPE
65197                 KED...PNINRMHESMKLFDSISNSKWFVSTSIILFLNKKDLFAEKILV..SPLKECFED
4493                  RED...RSKNRLIDSLEVFYQAWMNRYLQNVPIIVFVNKIDMLELKIQN..EHSIESMIN
HumanGNAO1[Ao]        HED...ETTNRMHESLMLFDSICNNKFFIDTSIILFLNKKDLFGEKIKK..SPLTICFPE
HumanGNAS[As]         RED...NQTNRLQEALNLFKSIWNNRWLRTISVILFLNKQDLLAEKVLAGKSKIEDYFPE
HumanGNA12[A12]       MED...RRTNRLVESMNIFETIVNNKLFFNVSIILFLNKMDLLVEKVKT..VSIKKHFPD
HumanGNAQ[Aq]         VES...DNENRMEESKALFRTIITYPWFQNSSVILFLNKKDLLEEKIMY..SHLVDYFPE
HumanGNAI1[Ai]        AED...EEMNRMHESMKLFDSICNNKWFTDTSIILFLNKKDLFEEKIKK..SPLTICYPE
```

```
RatGnai1[Ai]             ℓ.ℓℓℓ...............................................
                    290
                    .
RatGnai1[Ai]        Y.AGS.NTY.EEA...............................................
82535               ............................................................
1635                ............................................................
16751               ............................................................
2931                YPENFDPHNIVQVQQFLV..........................................
8947                FPGESDPYNLVDVQMFIV..........................................
33116               FKQEKCKHIFELIK.DLKAARGKRSKKESDIWEKYFSYFLPQMRDQKKTENPEDATRQQR
13658               Y.TGL..QEYDPS...............................................
23262               ............................................................
32381               ............................................................
21781               ............................................................
86418               ............................................................
9715                Y.KGP..QEYDSS...............................................
94774               ............................................................
1656                Y.TGR..QTYEEA...............................................
22522               FEGNP..SDVEDG...............................................
9665                FAEYQSKNTFDSQ...............................................
6167                YLRYTLPADVQCHN..............................................
5082                Y.KGN.NNF.EDA...............................................
3755                Y.EGP.QRDAEAA...............................................
1773                Y.ADE.NTY.EKA...............................................
77832               ............................................................
47050complete       Y.NGP.HNY.EET...............................................
19937               Y.LGP.NTF.VDT...............................................
12112               Y.TGA.NIY.EEA...............................................
65197               Y.TGN.NSY.NEA...............................................
4493                EITNLTPHSTKNSAKQLNNRRSSLN.....INHKTN...NDNMTSCSKCENSHPITDNNK
HumanGNAO1[Ao]      Y.TGP..NTYEDA...............................................
HumanGNAS[As]       FARYTTPEDATPE...............................................
HumanGNA12[A12]     F.RGD.PHRLEDV...............................................
HumanGNAQ[Aq]       Y.DGP.QRDAQAA...............................................
HumanGNAI1[Ai]      Y.AGS.NTY.EEA...............................................
```

```
                                         α15
RatGnai1[Ai]        ............................................................

RatGnai1[Ai]        ............................................................
82535               ............................................................
1635                ............................................................
16751               ............................................................
2931                ...............DS......................FV..SFVDNPGAH........
8947                ...............DS......................FV..KLIDNPYGN........
33116               ETQ..........IINE...YSQIQ.............SQVAKVFNDGSVDKWLTDGDDD
13658               ............................................................
23262               ............................................................
32381               ............................................................
21781               ............................................................
86418               ............................................................
9715                ............................................................
94774               ............................................................
1656                ............................................................
22522               ............................................................
9665                ............................................................
6167                ............................................................
5082                ............................................................
3755                ............................................................
1773                ............................................................
77832               ............................................................
47050complete       ............................................................
19937               ............................................................
12112               ............................................................
65197               ............................................................
4493                CIHLCPESNSNSKICDRKISIRTIPSCSSRRFSKDNNFEFVAQTFSENPWGKYQPSNEEC
HumanGNAO1[Ao]      ............................................................
HumanGNAS[As]       ............................................................
HumanGNA12[A12]     ............................................................
HumanGNAQ[Aq]       ............................................................
HumanGNAI1[Ai]      ............................................................
```

```
                                                                   α16
RatGnai1[Ai]        ...........................................
                                                        300           310

RatGnai1[Ai]        .......................................AAYIQCQFEDLNKRK......
82535               ............................................................
1635                ............................................................
16751               ............................................................
2931                ....................................P.RSQYSAGNQNQGR.ASIAG.QA
8947                ....................................DKRASYIRSNSVSGN.KRVVSPVP
33116               DIFSHLMKNDEF.........ISFHKRIMNITFYQVVSITFFIENQFLHQCE........
13658               .......................................IEYIEKRFRNEVKDS......
23262               ............................................................
32381               ............................................................
21781               ............................................................
86418               ............................................................
9715                .......................................VSYIEQCFRSKNKDV......
94774               ............................................................
1656                .......................................AAYIQASFEAKNNSP......
22522               .......................................INFFRMKFLSLKPAN......
9665                .............................SDEPSEVTYAKNFVKDKFMST.TKK......
6167                .............................HIEEAEFTRAKYFFRDEFLKITTTG......
5082                .......................................SEYVRMTFEMLNKKK......
3755                .......................................RDFILKMFIELNPDQ......
1773                .......................................VTYIKFKFESLNKYR......
77832               ............................................................
47050complete       .......................................SNFIRLKFEDLNKSK......
19937               .......................................SNYIREKFEGLNKKK......
12112               .......................................SAYIQSQFEELNKKK......
65197               .......................................SRYIQETFEMLNKKK......
4493                KEFVNSFPVDQVDLHSSGTHAAKKKSKDVHNIHPDTIKTACYIKNIFAQITRNHPKNCDM
HumanGNAO1[Ao]      .......................................AAYIQAQFESKNRSP......
HumanGNAS[As]       .............................PGEDPRVTRAKYFIRDEFLRISTAS......
HumanGNA12[A12]     .......................................QRYLVQCFDRKRRNR......
HumanGNAQ[Aq]       .......................................REFILKMFVDLNPDS......
HumanGNAI1[Ai]      .......................................AAYIQCQFEDLNKRK......
```

```
                             β6             α17
RatGnai1[Ai]                 →
                        320          330          340          350

RatGnai1[Ai]        ...DTKEIYTHFTCATDTKNVQFVFDAVTDVIIKNNLKDCGLF
82535               ...................IIKLVFTGSTIAFENASV......
1635                ...........................................
16751               ...........................................
2931                GVPKKRTLYRHFTTAVDQRNIETVFNAMKDTILQRNIDQLVMK
8947                MQIPKRTIYRHFTTAVDKSNIEKVFIAMKDTILQNNIRKIMMN
33116               .QTSNRHIYPYPTTAIDKRNVDRVFESCKDILQGKLLTEIMA.
13658               ....SKYVYCFHTCVIDTANIQAVFSATADFILSKNMKDLTIC
23262               ...........................................
32381               ...........................................
21781               ...........................................
86418               ...........................................
9715                ....SNEIYCHHTCATDTSNIQFVFDAVTDLIISNNMRGCGFY
94774               ...........................................
1656                ....NKEIYCHQTCATDTNNIQFVFDAVTDVIIANNLRGCGLY
22522               .APKTKKIYTHVTCCLDIDKMKFIIKQIIQNMLDSNVKRMTLF
9665                .EEGSRRCYPHFTCAVDTENIKRVFGDCQDMLQRIYMQKMGLM
6167                .NDGRHYCYSHFTCAVDTENIRRVFNDCKDIIQRMHLRQYELL
5082                ....EKIIYSHFTCATDTSNINYVFNVITDSIIVKNINAIGIF
3755                ....EKIIYSHFTCGTDTENIRFVFAAVKDTILQCNLKEYNLV
1773                ...QTKLIFTHITCATDTDNVKNVFEDIKTIMIQRLLELHGLM
77832               ...........................................
47050complete       ...DTKTIYSHFTCATDTNNIEVVFNAVIDVIIKNNLKDVGLF
19937               ...NSKVIYTHFTCATDTTNVQVVFDAVIDIIIKNNLKDVG..
12112               ...ETKTIYTHFTCATDTNNIQVVFDAVIDVIIKNNLKDCGLF
65197               ...LSKTIYTHFTCATDTNNVQVVFDAVIDVIIKNNLKDCGLF
4493                KISNRRKCLFYYTCAVNTDNIQKVLDGCRSFLMEQHLERFGIL
HumanGNAO1[Ao]      ....NKEIYCHMTCATDTNNIQVVFDAVTDIIIANNLRGCGLY
HumanGNAS[As]       .GDGRHYCYPHFTCAVDTENIRRVFNDCRDIIQRMHLRQYELL
HumanGNA12[A12]     ....SKPLFHHFTTAIDTENVRFVFHAVKDTILQENLKDIMLQ
HumanGNAQ[Aq]       ....DKIIYSHFTCATDTENIRFVFAAVKDTILQLNLKEYNLV
HumanGNAI1[Ai]      ...DTKEIYTHFTCATDTKNVQFVFDAVTDVIIKNNLKDCGLF
```
